# Mapping the regulatory wiring of endothelial cell differentiation reveals a critical role for Hippo mechanosignaling

**DOI:** 10.64898/2026.11.04.763740

**Authors:** Tri C. Nguyen, Hunter D. King, Dulguun Amgalan, Olga Pushkarev, Yanyan Alexandra Mo, Chad J. Munger, Chloe McCreery, Revant Gupta, Tony Zeng, Ronghao Zhou, Stephanie D. Conley, Surya Prakash Rao Batta, Maya U. Sheth, Eugenio Mattei, Anastasiia Masaltseva, Helen Y. Kang, X. Rosa Ma, Ngoc Tran, Liz Gaskell, Noam Shoresh, Valeria Orlova, Anshul Kundaje, Saulius Sumanas, Lay Teng Ang, Kyle M. Loh, Jesse M. Engreitz

## Abstract

At each step of development, stem cells encounter a branching decision to differentiate into one type of cell or another. This decision is executed through the action of competing gene programs that specify different cellular identities. Consequently, creating a comprehensive map of the gene programs that drive each branching step of human cell differentiation remains a long-sought goal of developmental and stem cell biology. Here, we introduce a framework combining genome-wide CRISPR screens, single-cell readouts, and gene program inference to systematically discover and characterize regulators of cell differentiation. This framework includes a method for CRISPR perturbations paired with single-cell RNA-seq called circular capture Perturb-seq (CC-Perturb-seq), which improves gRNA detection from split-pool combinatorial indexing protocols and enables perturbing many candidate regulators to map their effects on gene programs. We applied this framework to map how human pluripotent stem cells differentiate into arterial endothelial cells (ECs), and identified 174 regulators that promote or inhibit EC differentiation. By tracking gene programs across three stages of differentiation, we resolved when and how regulators divert cells toward alternative fates. This map revealed an unexpected requirement for the Hippo mechanosignaling pathway in EC specification. During lateral mesoderm differentiation, Hippo mechanosignaling was required to activate *ETV2*, the master transcriptional regulator of endothelial identity. Inhibiting Hippo blocked *ETV2* even in the presence of VEGF and diverted progenitors to a mesenchymal fate *in vitro*. Likewise, Hippo signaling was required for maximal *etv2* expression in developing zebrafish embryos. Conversely, pharmacologically activating the Hippo pathway increased the purity of both arterial and venous EC differentiation *in vitro*. Our framework thus identifies novel regulators of endothelial cell differentiation, improves stem-cell differentiation protocols via manipulation of mechanosignaling, and provides a generalizable platform for mapping the regulatory wiring of human development.

## Introduction

During directed differentiation of stem cells, some adopt the desired fate, while others deviate.^1^ Understanding *why* is an aspiration of developmental and stem cell biology, and will require mapping the interplay of signaling pathways, transcriptional regulators, and downstream gene programs that execute differentiation.^2,3^ Toward this goal, an ideal map would describe: which genes and downstream gene programs regulate cellular differentiation, at which times, and toward what alternative cell fates? Such a genetic map of differentiation could reveal new molecular mechanisms that drive development and improve our ability to guide stem cell differentiation for regenerative medicine.

An exemplary system where such a map could be transformative is the differentiation of artery endothelial cells (ECs) from human pluripotent stem cells (hPSCs).^4^ Arteries convey blood, nutrients, hormones, and immune cells to all organs.^5^ We recently showed that arterial ECs can be efficiently derived from hPSCs *in vitro* by mimicking the stepwise embryonic progression of pluripotent stem cells to primitive streak to lateral mesoderm to ECs.^4^ This protocol involves supplying key growth factors or inhibitors needed to specify endothelial identity, including VEGF.^6–10^ Yet, the gene programs that enact or counteract these signals remain to be discovered.

New tools could now enable building such a map. Toward discovering genes that regulate differentiation, forward genetic screens can now be applied at unprecedented scale in model organisms ^11–15^ and in differentiating human cells *in vitro*.^16–22^ Toward characterizing the effects of these regulators, Perturb-seq now enables measuring the transcriptional consequences of gene perturbations, by combining CRISPR with single-cell RNA sequencing.^23–26^ However, Perturb-seq has limitations in scale and cost, and these tools have yet to be fully integrated to systematically connect regulators to their downstream gene programs across differentiation.

Here we introduce a framework and technology (CC-Perturb-seq) for mapping the gene programs that drive cellular differentiation, and apply it to discover novel regulators of arterial EC differentiation (**Fig. 1a**). First, we systematically searched for and discovered 174 regulators that promote or repress arterial EC differentiation through a genome-wide CRISPR interference (CRISPRi) screen.^27^ Second, we used CC-Perturb-seq to knock down these regulators and characterize their transcriptional consequences across multiple stages of differentiation. Third, we distilled these transcriptional consequences into gene programs through consensus non-negative matrix factorization (cNMF)^28^, and tracked the effects of regulators across time. A key element of this framework is circular capture (CC)-Perturb-seq, which improves perturbation assignment and scalability of split-pool Perturb-seq^29,30^ by capturing Cas9 gRNAs via circularization of their cDNAs. Together, this framework generated a precise temporal and mechanistic view of when and how differentiation becomes derailed upon the loss of each regulator.

**Figure 1.**
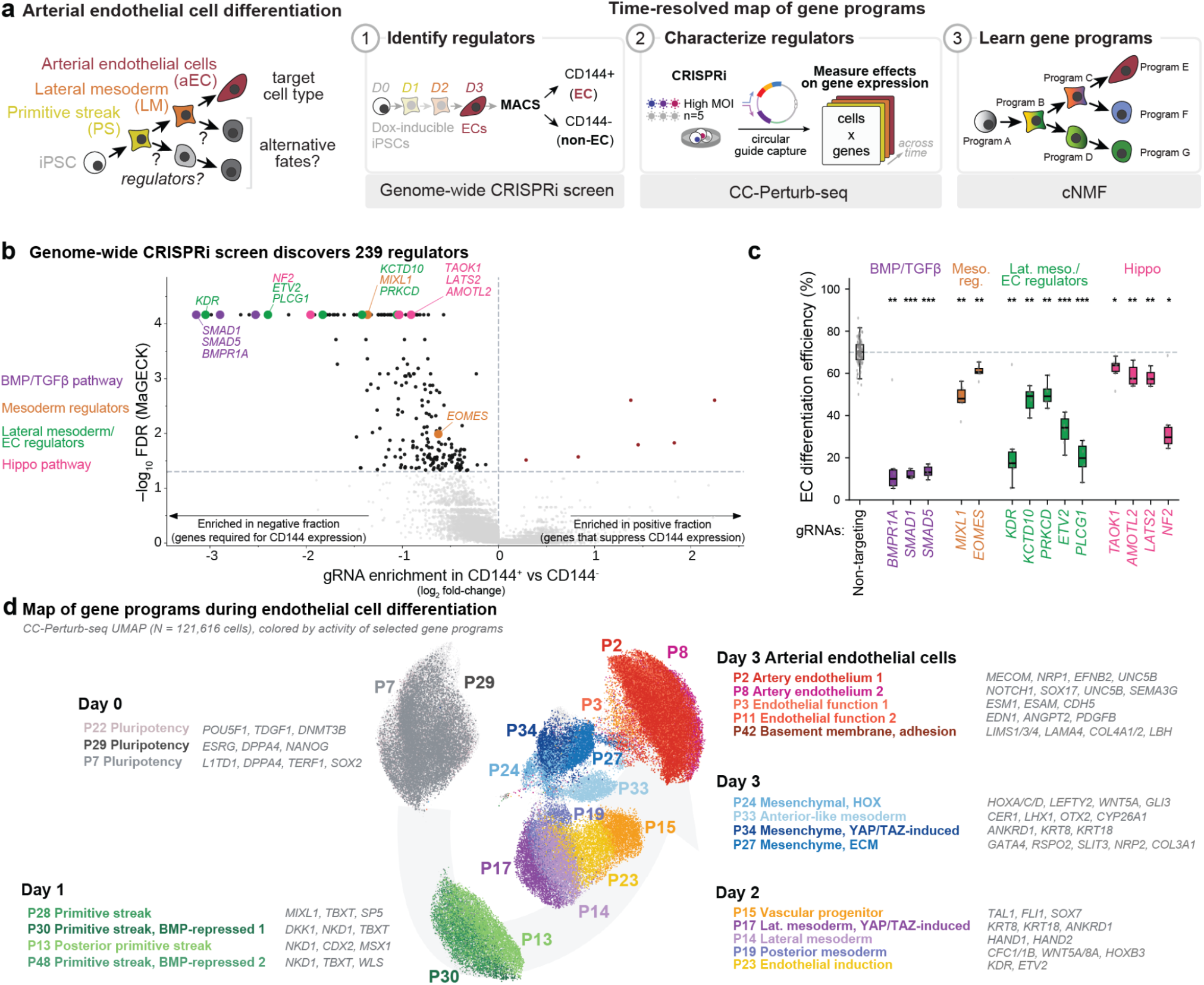
A time-resolved map of regulators and gene programs during arterial endothelial differentiation. **a.** Framework to identify and characterize regulators of arterial EC differentiation. A genome-wide CRISPRi enrichment screen identifies genes that alter CD144+ endothelial differentiation efficiency. A focused high-MOI CC-Perturb-seq experiment characterizes selected genes at Days 0–3 and measures the transcriptional profiles of alternative cell states and fates. Unsupervised matrix factorization with cNMF maps perturbations to coordinated sets of co-expressed genes (“programs”) and the differentiation timepoints at which they act. **b.** Genome-wide CRISPRi screen. Each point is a gene. X-axis: log₂ guide enrichment in CD144+ versus CD144− cells. Y-axis: −log_10_ FDR from MAGeCK. Black: Genes significantly depleted from the CD144+ fraction. Red: Genes significantly enriched in the CD144+ fraction. Selected hits are labeled and colored by functional group. Horizontal dashed line: FDR < 0.05. **c.** Estimated efficiency of EC differentiation (% of CD144+ cells on Day 3) for selected genes from the validation CRISPRi experiment. Dots: Guides. Boxplot: Mean per gene +/- interquartile range. Dashed line: mean estimated EC differentiation efficiency for negative control gRNAs. *FDR*: Benjamini-Hochberg corrected Welch’s t-test. * FDR < 0.05, ** FDR < 0.01, and *** FDR < 0.001 **d.** UMAP of 121,616 CC-Perturb-seq cells from Days 0–3, colored by the selected cNMF program with the highest activity in each cell. Labels describe program annotation and representative marker genes.

Unexpectedly, this temporally-resolved map of gene programs revealed that the Hippo mechanosignaling pathway was critical for human EC differentiation. Upon Hippo inhibition, differentiating lateral mesoderm cells failed to acquire EC identity *in vitro*, even when exposed to VEGF; they instead adopted an alternative mesenchymal fate. Conversely, pharmacological activation of the Hippo signaling pathway maximized hPSC differentiation into both arterial and venous ECs. Mechanistically, Hippo and VEGF were combinatorially required to upregulate *ETV2*, the transcriptional master regulator of endothelial identity. Manipulating mechanosignaling could be a broader strategy to enhance hPSC differentiation into additional lineages beyond ECs. Taken together, this demonstrates the power of this framework to discover hitherto unknown regulators and mechanisms of differentiation and improve our ability to direct cellular differentiation. This framework could be more generally applied to understand the gene programs that regulate the branching differentiation steps leading to many other cell types across each stage of development.

## Results

### Identifying regulators of arterial EC differentiation

We developed an approach to systematically map gene programs that regulate the differentiation of arterial EC from hPSCs (**Fig. 1a**). We implemented multiple functional genomics screens in the context of a 3-day hiPSC-to-arterial EC differentiation protocol. This protocol begins with hiPSCs and, emulating the stepwise development of ECs *in vivo*, progresses into primitive streak, lateral mesoderm, and then to arterial ECs (**Fig. 1a**), yielding 71.9% arterial ECs in the genome-wide screen^4^.

To identify genes that affect arterial EC differentiation efficiency, we conducted a genome-wide CRISPRi screen targeting the promoters of 20,528 genes, encompassing 20,655 promoters in total (**Fig. 1a, Fig. S1**). We delivered the library of sgRNAs into hiPSCs expressing doxycycline-inducible dCas9–KRAB, induced CRISPRi with doxycycline for three days before differentiation, and continued knockdown throughout the subsequent three-day arterial EC differentiation. On Day 3, an aliquot of each culture was analyzed by flow cytometry for endothelial cell-surface markers CD144 (otherwise known as VE-cadherin, encoded by *CDH5*) and CD31 to quantify differentiation efficiency (**Fig. S1**). The remaining cells were separated into CD144+ endothelial and CD144− non-endothelial fractions using magnetic-activated cell sorting (MACS) (**Fig. 1a**). We then sequenced sgRNAs recovered from each fraction to determine how individual gene perturbations affected EC differentiation efficiency. We identified 239 genes whose perturbation significantly increased or decreased the proportion of CD144+ cells on Day 3 (MaGECK FDR < 0.05, **Fig. 1b**). Effect sizes correlated in a focused validation screen (Pearson’s r = 0.72, *P* = 1.85 × 10^-40^, **Fig. S1d**).

The 239 candidate regulators included positive control *CDH5* (encoding CD144), known regulators of endothelial cell specification such as *KDR* (encoding VEGFR2)^6,10^ and *ETV2* (encoding the transcriptional master regulator of EC specification)^10,31,32^ (**Fig. 1b,c**), and many genes that have not been previously linked to arterial EC differentiation (**Table S2** and **S5**). These genes included members of the Hippo pathway, which were required for arterial EC differentiation and are explored further below (**Fig. 1c**).

### Mapping gene programs that regulate arterial EC differentiation

To characterize when and how these genes affect arterial EC differentiation, we developed an improved method called circular capture Perturb-seq (CC-Perturb-seq) for scalable mapping of gene programs (**Fig. S2**). As in Perturb-seq,^23–25^ we knock down genes with CRISPRi and read out their effects using single-cell RNA-seq. To improve scalability, we use split-pool combinatorial indexing^33^ to generate single-cell RNA-seq libraries, and improve the efficiency of gRNA detection by directly capturing the Pol III gRNA transcript, first by reverse transcription using a gRNA-specific primer, and then by circularization and inverse PCR of the resulting cDNA (“circular capture”, **Fig. S2a**). By capturing Pol III guides directly, CC-Perturb-seq improves gRNA detection efficiency and provides flexibility to detect gRNAs from many expression vectors (**Fig. S2**).

We used CC-Perturb-seq to knock down each of the 239 candidate regulators and 58 additional genes in hiPSCs, and read out the effects on gene expression at 4 timepoints before and during differentiation towards arterial ECs (**Fig. 1a**). Using the same timeline as the genome-wide screen, we transduced cells at high multiplicity of infection (MOI) with a library comprising 1,806 sgRNAs targeting 297 genes (∼6 sgRNAs per gene), and maintained doxycycline throughout differentiation. We collected cells immediately before differentiation as the Day 0 hiPSC sample and every 24 hours thereafter, profiling Day 1 primitive streak, Day 2 lateral mesoderm and Day 3 arterial ECs. We obtained data for 121,616 high-quality single cells (**Fig. S3**). Library quality and gRNA detection were robust, with an average of 2,472 transcriptome UMIs/cell from 7,850 transcriptome reads/cell, 45 UMIs/gRNA/cell, and an apparent MOI of 5–6 (**Fig. S3**). Across all targeted genes, we obtained on average 50.8% knockdown, which was consistent across timepoints (**Fig. S3f**). Altogether, after excluding genes with a close neighbor where CRISPRi targeting may be ambiguous, CC-Perturb-seq confirmed and characterized the effects of 174 regulators (**Table S1**) that significantly affected arterial EC differentiation (Fisher’s test FDR < 0.05).

We used these CC-Perturb-seq data to map which genes regulate which gene programs at which timepoints during arterial EC specification. Following our previous approach,^34^ we distilled co-regulated genes in this dataset into “gene programs” using cNMF^28^, identified genes whose perturbation affected each program, and annotated each program according to its linked genes and timepoint(s) of highest activity (**Fig. 1a**, **Fig. 2b, Fig. S4**, see **Methods**). We further collected 10x Multiome scRNA+ATAC data at each of the 4 timepoints, and annotated predicted transcription factor binding sites in enhancers linked to each program, using ChromBPNet^35^ and scE2G^36^ (**Fig. S5**).

**Figure 2.**
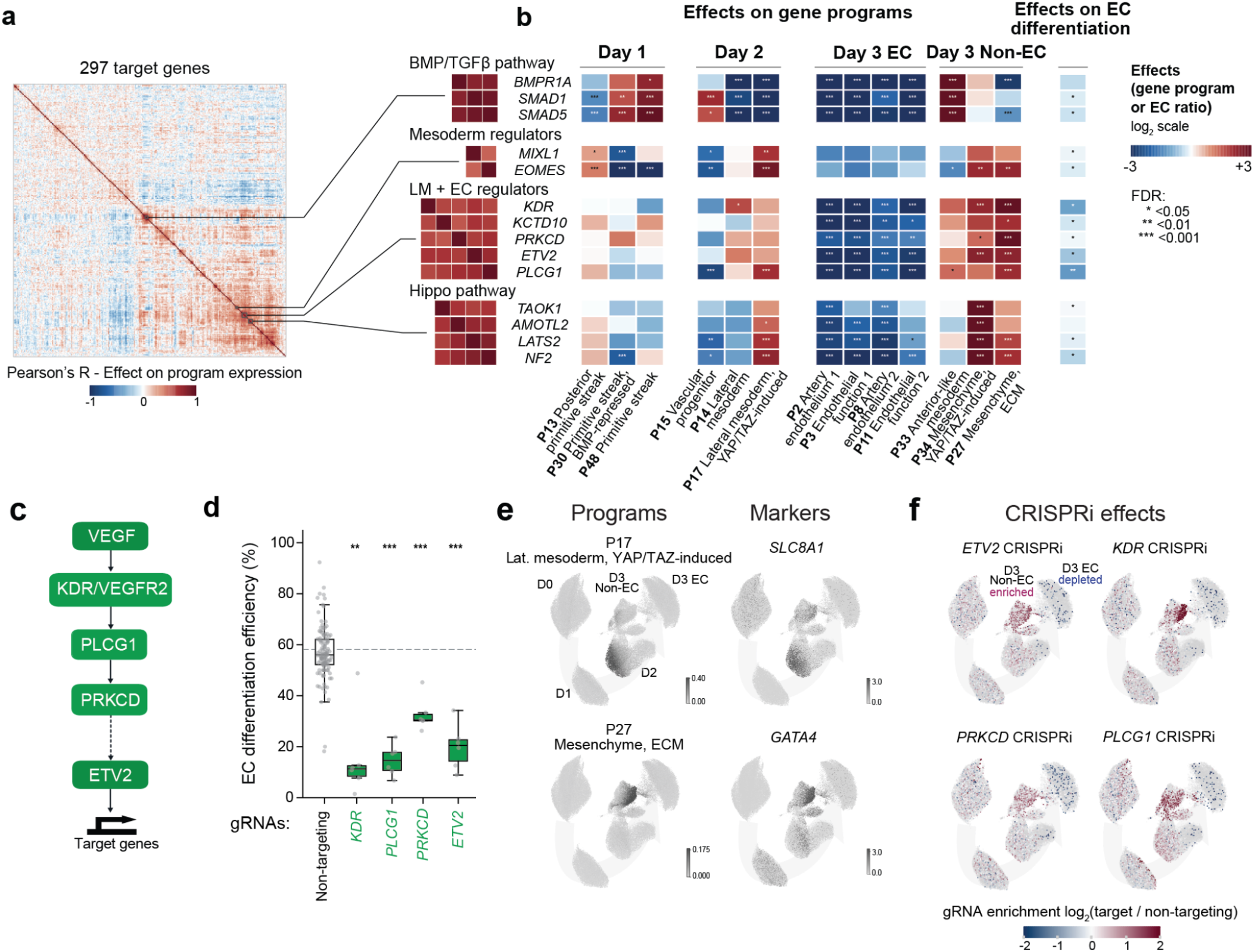
Distinct perturbation programs converge on failed endothelial differentiation. **a.** Heatmaps: pairwise correlations of gene perturbation effects across all cNMF programs and time points. Selected clusters of regulators are expanded at right. **b.** Effects of selected regulators on selected Day 1, Day 2 and Day 3 programs and on Day 3 EC differentiation. Heatmap shows signed perturbation effects. Asterisks denote significance (FDR, conditional randomization test from CC-Perturb-seq). **c.** Schematic of the VEGF–KDR–PLCG1–PKC signaling pathway and its relationship to *ETV2* induction. Solid arrows: established signaling steps. Dashed arrow: *ETV2* is inferred to be downstream of *PRKCD*. **d.** Day 3 EC fraction among (green) cells carrying guides targeting *ETV2*, *KDR*, *PLCG1* or *PRKCD* or (black) cells carrying non-targeting guides, excluding cells with guides targeting any of these 4 genes. Each point represents one guide. Boxes: median and interquartile range. Significance was assessed by Welch’s t-test. * FDR < 0.05, ** FDR < 0.01, and *** FDR < 0.001 **e.** UMAPs showing activity of Programs P17 and P27 and expression of representative marker genes *SLC8A1* and *GATA4* in CC-Perturb-seq data. **f.** Enrichment or depletion of cells with knockdown of *ETV2, KDR, PRKCD* or *PLCG1* across cell types and states in CC-Perturb-seq. Colored dots: Cells carrying guides targeting the indicated gene. Gray dots: All other cells. Red and blue indicate local enrichment and depletion, respectively, in program space, of cells carrying guides targeting the gene relative to cells (i) carry non-targeting control guides and (ii) do not carry guides targeting the gene. For example, cells carrying gRNAs targeting *ETV2* are enriched in the non-EC cell fraction on Day 3.

We obtained a map of 50 gene programs, each with 8–86 upstream regulators (**Fig. 1d, Table S4**). Cell identity programs defined various cell states during differentiation, including pluripotency (*e.g.,* Program 29, expressed on Day 0, with marker genes *ESRG, DPPA4,* and *NANOG*) and primitive streak (*e.g.,* Program 28, expressed on Day 1, with marker genes *MIXL1*, *TBXT,* and *SP5*) (**Fig. 1d**, **Fig. S6**). Later during differentiation, some gene programs were associated with diverse cellular processes in Day 3 endothelial cells including genes involved in angiogenesis, arterial identity, or the basement membrane (**Fig. 1d**). By contrast, other gene programs on Day 3 were associated with alternative non-endothelial fates, containing genes characteristic of mesenchymal or mesoderm-like cells (**Fig. 1d**). This is consistent with how, during hPSC differentiation towards arterial ECs, a minority of cells stray and instead adopt mesenchymal identity^4^.

### Inhibition of EC regulators diverts cells to alternative fates

This map of gene programs allowed us to identify groups of regulators with similar functions, trace the effects of gene perturbations over time, and define alternative fates (**Fig. 2a,b**). We clustered together regulators whose knockdown led to similar transcriptional consequences, reasoning this might define regulators that belonged in the same biological pathway, some of which might represent new pathways.

One cluster of regulators with similar effects on gene programs involved the known regulators *KDR* (VEGFR2) and *ETV2*, along with several other genes that appear to act downstream of VEGFR signaling: *PLCG1* and *PRKCD* (**Fig. 2c**). VEGF signaling through VEGFR2 is known to upregulate *ETV2*^10,37^, the transcriptional master regulator of endothelial specification^31,32,38,39^ The correlated perturbation profiles of these genes therefore suggested that PLCG1 and PKCδ participate in the intracellular signaling linking VEGFR2 activation to *ETV2* upregulation. Knockdown of these genes first affected gene programs at Day 2, specifically by increasing Program 17 (“Lateral mesoderm, YAP/TAZ induced”, **Fig. 2b**). By Day 3, knockdown of these genes led to significant reductions in CD144+ ECs (**Fig. 2d**) and an increase in Program 27—a mesenchymal program expressing the transcription factor *GATA4*^40^ as well as extracellular matrix genes such as *COL3A1*, *COL1A1*, and *FN1* (**Fig. 2e, Fig. S6**). Mechanistically, KDR binds PLCG1^41^, generating diacylglycerol that activates protein kinase C (PKC)^42^ (**Fig. 2c**). Across PKC isoforms, which vary in their requirements in different EC contexts and functions^43–45^, CRISPRi effects pointed to PKCδ (*PRKCD*) as the relevant isoform in this context (**Fig. S7**). Thus, perturbations at multiple points in the VEGF signaling cascade can divert arterial ECs to an alternative mesenchymal cell fate.

Perturbing BMP signaling (*BMPR1A*, *ACVR2A*, or *SMAD1/4/5*) or the primitive streak transcription factors *EOMES* and *MIXL1* likewise compromised EC differentiation, but in each case, differentiation failed for different reasons (**Fig. 2a,b**, **Fig. S8**). Knockdown of BMP signaling diverted cells towards an anterior-like mesoderm identity, consistent with how BMP is posteriorizing *in vivo*.^46^ Conversely, *EOMES* or *MIXL1* knockdown instead redirected cells towards mesenchymal lineages (**Fig. S8**).

Thus, genes with similar effects on overall EC differentiation efficiency often affected distinct gene programs throughout differentiation and led to distinct alternative fates. This map provides a rich resource for understanding the temporal decisions underlying endothelial cell specification.

### Hippo pathway inhibition diverts lateral mesoderm differentiation away from ECs and towards mesenchymal cells

Our map of gene programs revealed a surprising requirement for the Hippo pathway in EC specification (**Fig. 3a**). In brief, the mammalian Hippo pathway comprises several upstream components which are activated by different mechanical inputs including cell density, size, and shape as well as extracellular matrix (ECM) stiffness.^47–50^ These components converge to activate LATS1/2 kinases, which then phosphorylate the transcriptional coactivators YAP/TAZ, thus preventing YAP/TAZ from translocating into the nucleus and interacting with TEAD transcription factors to regulate target genes^47–50^ (**Fig. 3a**). In summary, Hippo pathway activation represses YAP/TAZ, whereas Hippo inhibition leads to nuclear YAP/TAZ localization (**Fig. 3a**). Individually knocking down four components of the Hippo pathway (*NF2*, *LATS2*, *AMOTL2*, or *TAOK1*) by 43–76% impaired EC differentiation (**Fig. 3c**, **Fig. S9**). CC-Perturb-seq revealed that knockdown of Hippo pathway genes decreased the proportion of ECs at Day 3, and reciprocally increased the proportion of cells that express alternative mesenchymal gene programs (**Fig. 3b,c**, **Fig. 2b**). This was unexpected: while the Hippo pathway is known to operate in differentiated ECs and mediates their responses to shear stress and hydrostatic pressure,^51–55^ less is known about the role of Hippo signaling in the incipient specification of ECs from mesoderm.

**Figure 3.**
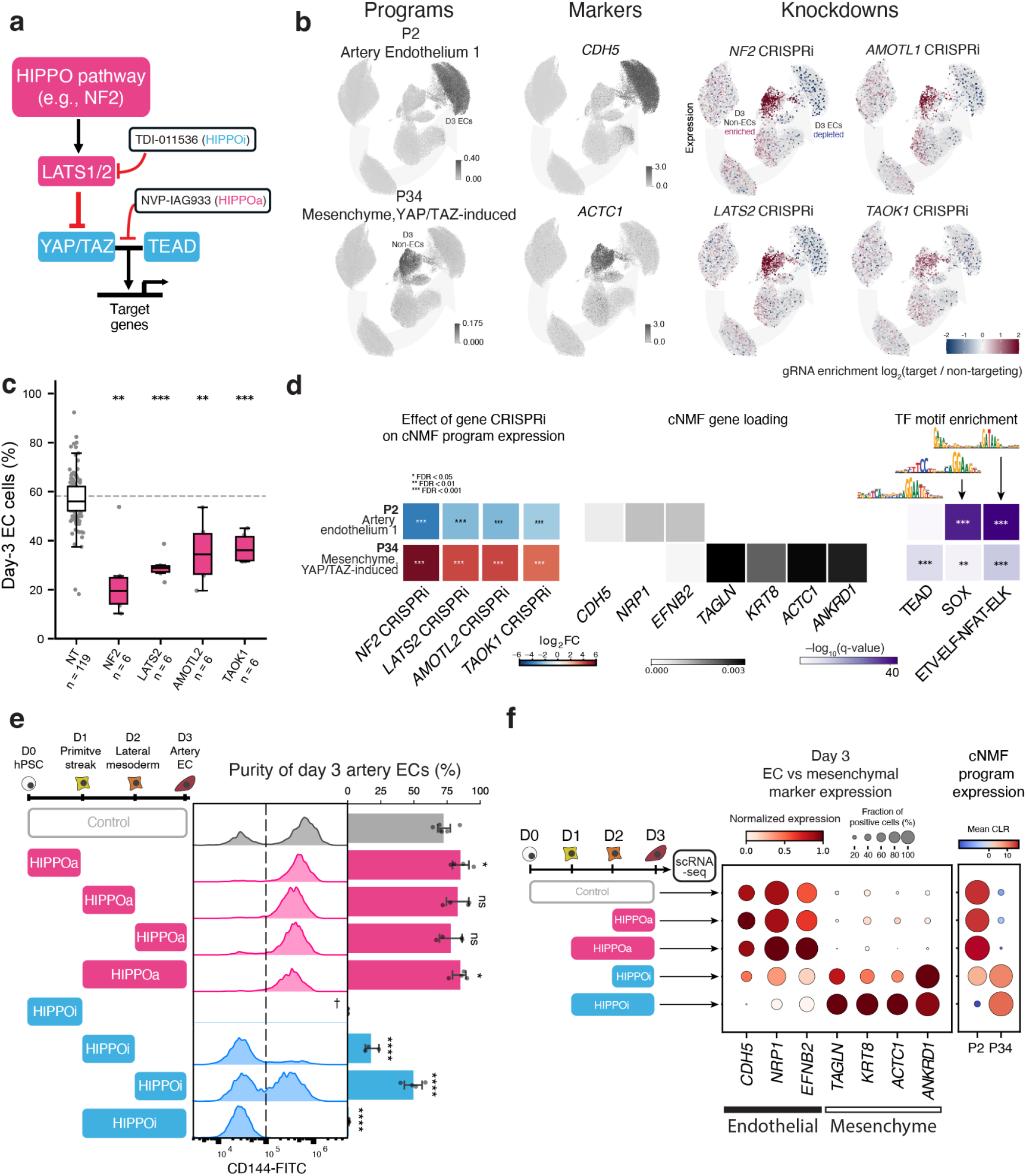
Hippo signaling is required for efficient endothelial differentiation. **a.** Hippo pathway schematic. Upstream components, including NF2, AMOTL2 and TAOK1, activate LATS1/2, which phosphorylate YAP/TAZ and limit TEAD-dependent transcription. TDI-011536 (HIPPOi) inhibits LATS1/2; NVP-IAG933 (HIPPOa) disrupts YAP/TAZ–TEAD binding. **b.** UMAP plots showing endothelial Program 2, mesenchymal Program 34, representative marker genes *CDH5* and *ACTC1*, and local enrichment of *NF2*, *LATS2*, *AMOTL2* and *TAOK1* perturbations. **c.** Day 3 EC fraction for non-targeting guides and guides targeting *NF2*, *LATS2*, *AMOTL2* or *TAOK1*. Points represent guides; boxes show the median and interquartile range. **d.** Annotation of P2 and P34. Left: CRISPRi effects on program activity. Middle: cNMF gene loading for selected marker genes. Right: enrichment of transcription factor motifs in linked enhancers. Asterisks denote adjusted significance. **e.** Flow-cytometry distributions and quantified CD144+ fraction after HIPPOa or HIPPOi treatment during the indicated 24- or 48-hour windows of the three-day differentiation. Points show biological replicates, n=4 total replicates with two replicates harvested from each of two independent experiments. Control arteries include two additional replicates from the second independent experiment. Differences from control were assessed using an additive two-factor ANOVA with condition and plate block as fixed effects, followed by two-sided, model-based Dunnett adjustment across all seven treatment-versus-control comparisons. \**P* < 0.05, \*\**P* < 0.01, \*\*\**P* < 10^-3^, \*\*\*\**P* < 10^-4^, ns: not significant. **f.** Marker-gene and cNMF-program expression in single-cell RNA-seq of Day 3 cultures treated with HIPPOa or HIPPOi during the final 24 or 48 hours. Dot size indicates the fraction of expressing cells and color indicates normalized expression (genes) or mean CLR-transformed usage (programs).

The rich transcriptional data afforded by CC-Perturb-seq provided a detailed view of how knockdown of Hippo pathway genes impacts gene expression (**Fig. 3b,d**). CRISPRi knockdown of any of the four Hippo pathway members led to a significant increase by Day 3 of differentiation in the transcriptional activity of Program 34 (“Mesenchyme, YAP/TAZ-induced”), which expressed known YAP/TAZ target genes (*ANKRD1*, *CCN1*, and *CCN2*)^56^ and mesenchymal markers (*MYOCD*, *ACTA2*, and *ACTC1*). Predicted binding sites for TEAD were indeed strongly enriched in predicted enhancers of genes in Program 34 (**Fig. 3d**, **Fig. S6**). CRISPRi knockdown of *NF2* or *LATS2* also upregulated the transcriptional activity of Program 27 (“Mesenchymal, structural”), which included extracellular matrix genes (*COL1A1*, *COL1A2*, *COL3A1*, and *COL5A1*) characteristic of mesenchymal cells.^57^

To test if and when Hippo signaling is required during EC differentiation, we employed fast-acting and reversible small-molecule modulators of the Hippo pathway. During hPSC differentiation, cells were exposed to a Hippo pathway inhibitor (“HIPPOi”; TDI-011536, an inhibitor of both LATS1 and LATS2 kinases)^58^ (**Fig. 3a**). Treatment effects on Day 3 CD144-positive purity were assessed using an additive two-factor ANOVA with treatment condition and plate block as fixed effects, followed by two-sided Dunnett-adjusted comparisons of each treatment with control (**Fig. 3e**; Methods). Hippo inhibition for 24 hours during lateral mesoderm differentiation (*i.e.*, day 1–2) potently inhibited the subsequent emergence of ECs by Day 3 (**Fig. 3e**, adjusted *P* < 2.20 × 10^-12^). Hippo pathway inhibition for 48 hours (i.e., Day 1–3) completely abolished EC differentiation (**Fig. 3e**, adjusted *P* < 2.51 × 10^-15^). However, Hippo inhibition during the final transition from lateral mesoderm to ECs (Day 2–3) only modestly reduced EC purity (**Fig. 3e**, adjusted *P* = 3.04 × 10^-6^), indicating a weaker requirement during this later interval than during Day 1–2 lateral mesoderm induction. This suggests that Hippo is specifically required during lateral mesoderm differentiation for subsequent EC differentiation.

To further confirm the cellular phenotypes, we profiled cells treated with Hippo pathway modulators with single-cell RNA-seq at various timepoints (**Fig. 3f**, **Fig. S12**). Consistent with the CC-Perturb-seq data entailing CRISPRi perturbations, we found that pharmacological HIPPOi treatment reduced expression of endothelial markers and programs (e.g., *CDH5*, *NRP1*, and *EFNB2*), and increased YAP/TAZ target genes (*ANKRD1*) and mesenchymal markers and programs (e.g., *TAGLN*, *KRT8*, and *ACTC1*) (**Fig. 3f**).

Together, these data show that Hippo signaling during lateral mesoderm differentiation is critical for EC differentiation, and that inhibiting Hippo diverts cells towards a mesenchymal fate.

### Hippo pathway activation improves EC differentiation efficiency

Because inhibition of the Hippo pathway decreased EC differentiation, we wondered whether activation could improve it. We tested this by adding a Hippo pathway agonist (“HIPPOa”; NVP-IAG933, an inhibitor of the YAP/TAZ-TEAD interface^59^) during various intervals of differentiation (**Fig. 3a**). Indeed, Hippo pathway activation during Day 1–2 or Day 1–3 of differentiation increased CD144+ EC purity from a 72.35% baseline to 82.93% or 84.98% respectively (*P* = 0.0786 or 0.0215, respectively) (**Fig. 3e**). HIPPOa significantly increased the expression of EC markers on Day 3 and significantly decreased the expression of mesenchymal markers (**Fig. 3f**).

Because both HIPPOi and HIPPOa treatment from Day 1–2 impacted EC purity in opposing directions, we hypothesized that levels of endogenous Hippo signaling might vary between individual cells during arterial EC differentiation. Indeed, immunostaining on Day 2 of differentiation revealed that some cells displayed YAP1 exclusively in the nucleus—indicative of low Hippo signaling—while others displayed YAP1 in both the nucleus and the cytoplasm—indicative of moderate Hippo signaling (**Fig. S10a**). Thus, Day 2 vascular lateral mesoderm cells exhibit heterogeneous Hippo activity, which likely explains how HIPPOi and HIPPOa guide differentiation more homogeneously into non-ECs or ECs, respectively.

In parallel to arterial ECs, venous ECs can also be derived from a protocol that follows the same path from primitive streak to lateral mesoderm, followed by a two-day venous EC induction step.4 HIPPOa treatment during lateral mesoderm induction (day 1–2) similarly increased the efficiency of venous EC specification, from 63.15% to 83.05% (**Fig. S11a**). Taken together, pharmacological activation of the Hippo pathway improves the differentiation efficiency of both artery and venous ECs *in vitro*.

### Hippo signaling acts with VEGF to activate *ETV2*, the EC master regulator

We next investigated with higher temporal resolution how Hippo gene perturbations affect cell state transitions during Days 1–2 of EC differentiation. *In vivo*, *Hand1*+ lateral mesoderm progenitors encounter a decision to differentiate into ECs or mesenchymal cells; differentiation into ECs entails sequential expression of *ETV2* and *TAL1/SCL* in vascular lateral mesoderm cells, also variously known as angioblasts or endothelial progenitors.^60–63^ Likewise, the 24-hour “lateral mesoderm” induction stage reproduces this sequence of transcription factor activation *in vitro^4^* (**Fig. 4a**). Accordingly, we treated Day-1 primitive streak cells with lateral mesoderm induction media in the presence or absence of HIPPOi or HIPPOa and measured the expression of all three transcription factors every 4 hours. Strikingly, HIPPOi led to sustained expression of *HAND1* and blocked the activation of *ETV2* and *TAL1* (**Fig. 4a**). HIPPOa did the opposite, further elevating *ETV2* and *TAL1* during their respective expression waves (Fig. 4a). These reciprocal effects on HAND1 and ETV2 were also observed by immunostaining (**Fig. S10b**). Hippo signaling is thus required to induce *ETV2*,a master transcription factor both necessary and sufficient for EC identity.^31,32,38,39^

**Figure 4.**
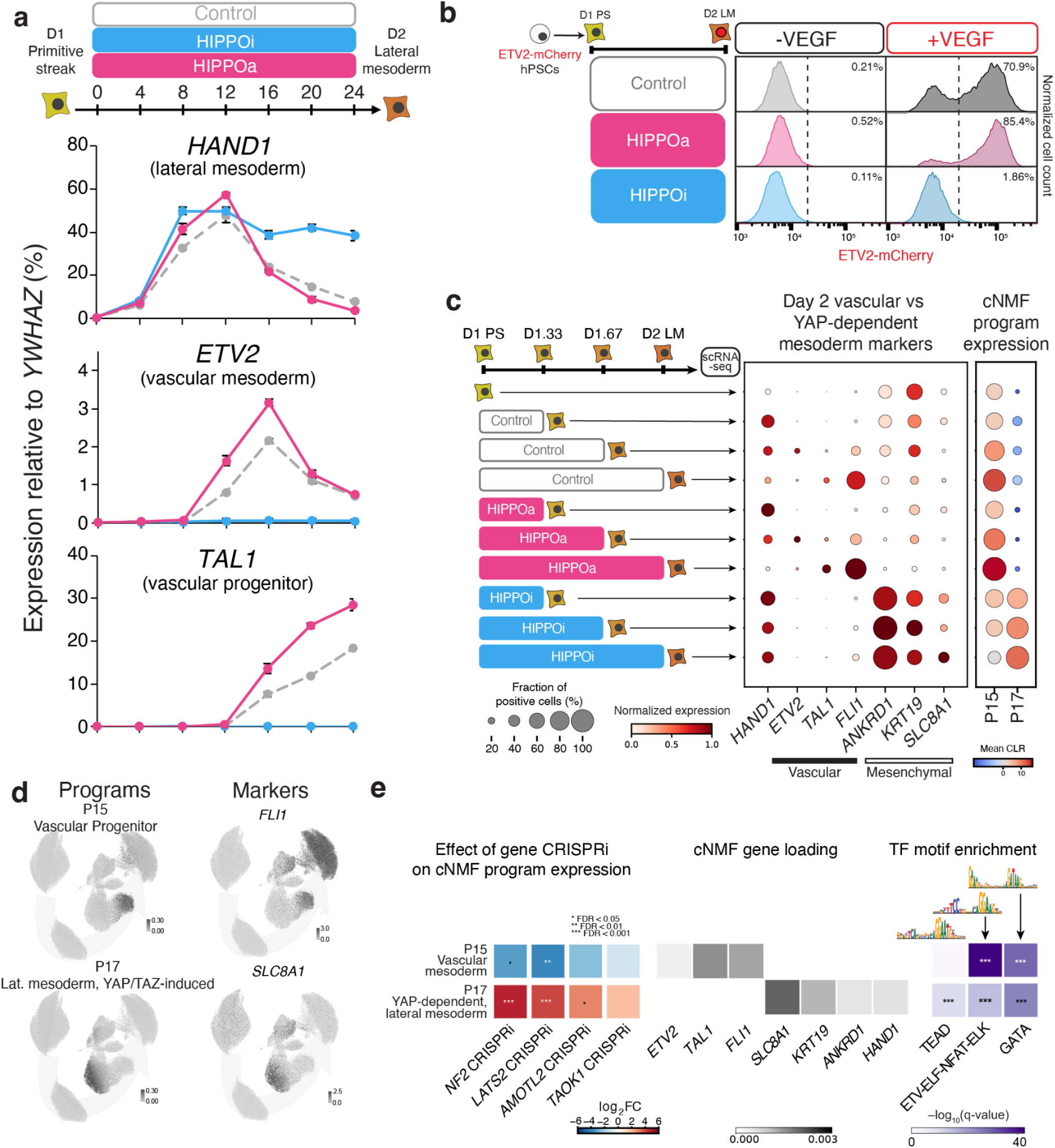
Hippo signaling promotes ETV2 and suppresses alternative lateral-mesoderm fates. **a.** Four-hour qPCR time course during Day 1–2 lateral-mesoderm induction under control, HIPPOa or HIPPOi conditions. Expression of *HAND1*, *ETV2* and *TAL1* was normalized to *YWHAZ*. Points represent average of two biological replicates **b.** ETV2–P2A–nuclear-mCherry reporter expression by flow cytometry after 24 hours of lateral-mesoderm induction with or without VEGF and with control, HIPPOa or HIPPOi treatment. Percentage of reporter-positive cells is indicated in each representative histogram. **c.** Single-cell expression of vascular- and mesenchymal-state markers and projected cNMF programs in cultures sampled every 8 hours (Day 1, Day 1.33, Day 1.67 and Day 2) under control, HIPPOa or HIPPOi treatment. Dot size indicates the fraction of positive cells and color indicates normalized expression (genes) or mean CLR-transformed usage (programs). **d.** UMAP plots showing activity of the Day 2 programs P15 and P17 and expression of representative markers *FLI1* and *SLC8A1* in the CC-Perturb-seq data. **e.** Integrated annotation of P15 and P17. Left, Hippo-pathway perturbation effects on program activity; middle, representative cNMF gene loadings; right, enrichment of transcription-factor motifs in linked enhancers.

VEGF upregulates *ETV2* to confer endothelial identity^10,37,61,64^, and we thus asked how combinations of Hippo and VEGF impact ETV2 expression. We tracked ETV2 expression using an endogenous *ETV2–P2A–mCherry* reporter hPSC line^65^, hereafter termed the *ETV2-mCherry* reporter. Cells were challenged with HIPPOi or HIPPOa, either in the presence or absence of VEGF, during lateral mesoderm induction (**Fig. 4b**). Cells activated *ETV2* only when both Hippo and VEGF were activated; all conditions lacking VEGF or treated with HIPPOi failed to activate *ETV2* (**Fig. 4b**). Given the prevailing model that VEGF is the principal driver of *ETV2* expression^10,37,61,64^, it was surprising that even in the presence of VEGF, cells would not turn on *ETV2* unless Hippo was simultaneously activated. VEGF and Hippo signaling therefore form an “AND” logic gate for activating *ETV2*. Interestingly, HIPPOa enhanced the percentage of *ETV2*+ cells above control (**Fig. 4b**), suggesting that endogenous Hippo activity is submaximal in the standard protocol and that making *ETV2* induction more uniform may explain how HIPPOa improves EC purity. This thus adds a new dimension to our knowledge of how EC identity is incipiently specified from mesoderm by showing that both VEGF and Hippo are conjointly required.

To understand the resulting phenotype of cells upon Hippo perturbations at the end of lateral mesoderm induction, we examined the CC-Perturb-seq data collected at Day 2. CRISPRi knockdown of Hippo pathway genes decreased Program 15 (“Vascular progenitor”) and increased Program 17 (“Lateral mesoderm, YAP/TAZ induced”), reflecting a shift in lateral mesoderm differentiation towards mesenchymal lineages and away from vascular lateral mesoderm (**Fig. 4d,e**). These programs represented distinct cell states observed on Day 2, with Program 15 representing cells expressing *ETV2*, *TAL1*, and other vascular progenitor genes such as *FLI1*, and Program 17 representing cells that aberrantly sustain *HAND1* expression, which also expressed YAP/TAZ-TEAD response genes such as *ANKRD1* and mesenchymal-associated genes *KRT19* and *SLC8A1* (**Fig. 4d,e**). We observed similar reciprocal effects on gene expression in cells treated with HIPPOi/a (**Fig. 4c**). In summary, Hippo signaling acts alongside VEGF to control the bifurcation of *HAND1*+ lateral mesoderm progenitors between vascular and mesenchymal fates.

### Mechanical cues modulate *ETV2* induction

Among developmental pathways, Hippo signaling is idiosyncratic because it is principally activated by mechanical cues such as high cell density or low matrix stiffness.^47–50^ While mechanical forces regulate multiple stages of development,^66^ less is known about how mechanical forces control early mesoderm lineage decisions. To test whether such mechanical cues upstream of Hippo can impact *ETV2* induction, we seeded primitive streak cells onto surfaces of varying stiffness (plastic, or alternatively, hydrogels of 0.5 kPa or 32 kPa stiffness) at varying cell densities, and exposed them to lateral mesoderm induction media (**Fig. 5a**). Consistent with how soft ECMs suppress YAP/TAZ (*i.e.*, activate the Hippo pathway)^47–50^, we observed decreased expression of YAP/TAZ target gene *ANKRD1* in cells plated on surfaces of decreasing stiffness (**Fig. 5b**). Conversely, *ETV2* increased as stiffness decreased (**Fig. 5b**). Higher cell density—which is known to activate Hippo signaling^47–50^—increased the purity of *ETV2*+ cells at each level of stiffness (**Fig. 5c**), consistent with the above observations that Hippo signaling is essential for *ETV2* induction. To test whether the Hippo–YAP/TAZ axis lies downstream of these mechanical inputs and upstream of *ETV2*, we tested whether pharmacological HIPPOa could rescue the reduction in *ETV2* caused by increased matrix stiffness or reduced cell density (**Fig. 5c**). Indeed, HIPPOa treatment increased the percentage of *ETV2+* cells replated on high-stiffness plastic culture dishes to levels comparable or greater than cells replated on 32 kPa or 0.5 kPa substrates in control media (**Fig. 5c**). Further, the difference in *ETV2*+ cell purity between 32 kPa and 0.5 kPa was masked when HIPPOa was added, indicative of HIPPOa rescuing any loss of *ETV2* on the stiffer substrate (**Fig. 5c**). We similarly found in cells plated at lower density that HIPPOa treatment rescued *ETV2* purity to comparable levels to cells seeded at high density (**Fig. S13a**). Taken together, cell density and surface stiffness are two mechanical inputs that impact *ETV2* induction through Hippo signaling.

**Figure 5.**
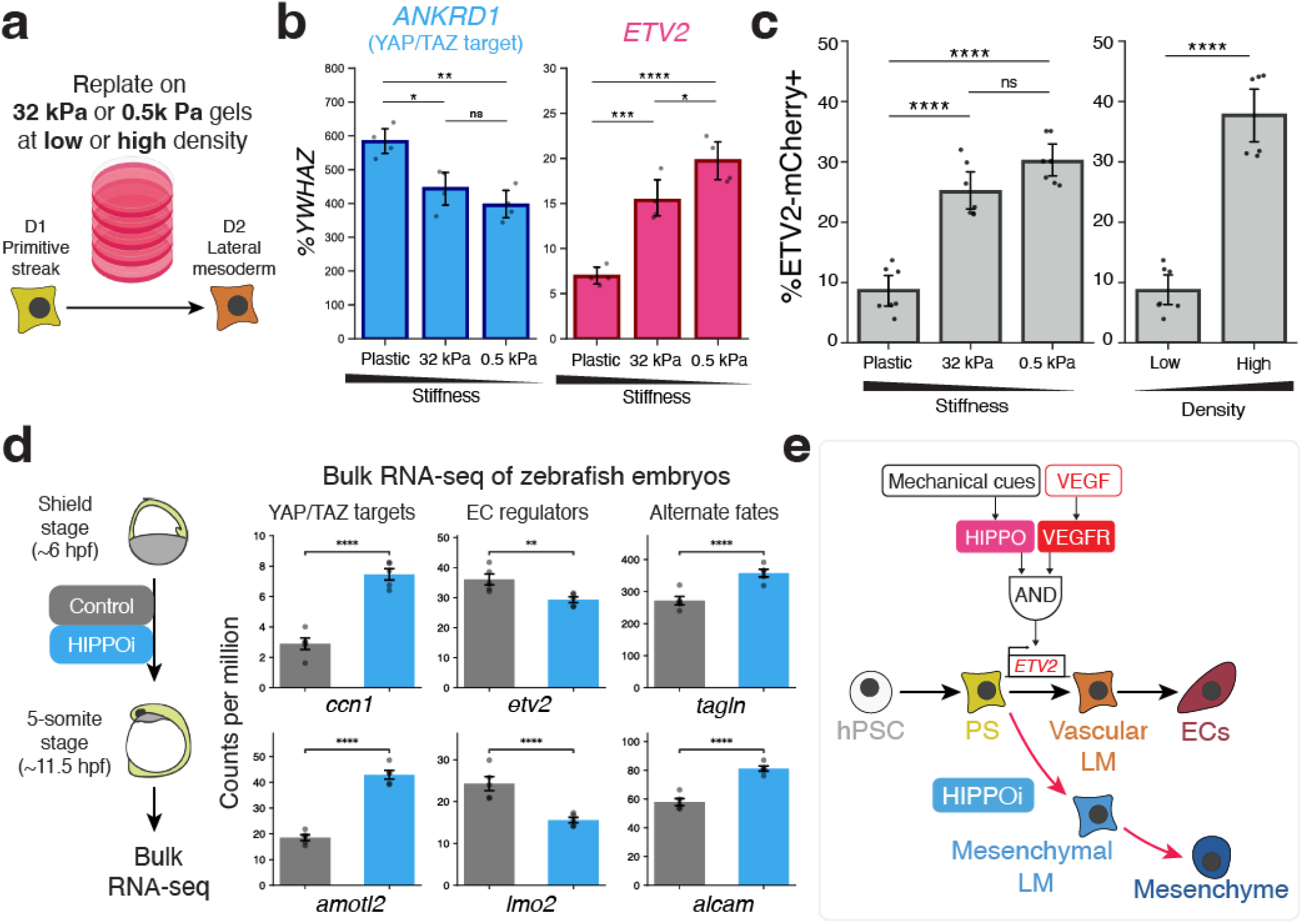
Hippo controls *ETV2* downstream of mechanical cues and promotes *etv2 in vivo*. **a.** ETV2-mCherry reporter hPSCs^65^ were differentiated to day 1 primitive streak then replated in lateral mesoderm media for 24 hours on substrates of varying stiffness. **b.** Expression of *ETV2* and the YAP/TAZ target *ANKRD1* measured by qPCR. For each gene, expression was normalized to the expression of the housekeeping gene *YWHAZ*. \**P* < 0.05, \*\**P* < 0.01, \*\*\**P* < 10^-3^, \*\*\*\**P* < 10^-4^, ns: not significant; Tukey’s Honestly Significant Difference test. n=4 total biological replicates, two replicates harvested from each of two independent experiments. **c.** Quantified ETV2-mCherry+ fraction by flow cytometry of cells replated onto either a standard plastic culture dish or onto substrates of specified stiffness at either low (2.6 × 10^4^ cells/cm^2^, “Sparse”) or high (7.8 × 10^4^ cells/cm^2^, “Dense”) cell density. Left: cells replated on substrates of varying stiffness at low density. Right: cells replated at different densities onto plastic culture dishes. Asterisks represent the same *P* value thresholds as in **b**; Tukey’s Honestly Significant Difference test. n=6 for the high density condition, n=7 for all other conditions; all across 3 independent experiments. **d.** Bulk RNA-seq expression of YAP/TAZ targets, endothelial regulators and alternative mesoderm markers in zebrafish embryos treated with DMSO or 10 μM HIPPOi from shield stage to the 5-somite stage. Points show n=5 biological replicates (embryos). Asterisks represent the same *P* value thresholds as in **b**; adjusted *P* value calculated by Benjamini-Hochberg procedure using DEseq2.^67^ **e.** Summary cartoon of key findings. Upstream mechanical inputs activate Hippo which, together with VEGF signaling, upregulates *ETV2* to promote EC fate. Hippo pathway inhibition diverts cells towards a mesenchymal fate.

### Hippo inhibition suppresses *etv2 in vivo*

To test whether Hippo signaling is also required for *etv2* induction *in vivo*, we treated zebrafish embryos with HIPPOi starting at the shield stage (6 hours post fertilization [hpf]) and analyzed *etv2* expression by *in situ* hybridization at the 5-somite stage (∼11.5 hpf) (**Fig. S13b,c**). HIPPOi treatment increased the proportion of embryos that displayed low *etv2* expression, relative to controls (30.2% HIPPOi vs 6.4% control, posterior view; 54.7% HIPPOi vs 8.5% control, anterior view; n = 47 embryos for control, 53 for HIPPOi treated; *P* = 2.53 × 10^−3^, posterior view; *P* = 2.65 × 10^−6^, anterior view; Fisher’s exact test) (**Fig. S13c**). Bulk RNA-seq of zebrafish embryos treated with HIPPOi over the same time window showed the expected upregulation of the YAP/TAZ targets *ccn1* and *amotl2*, confirming Hippo inhibition (**Fig. 5d**). EC regulators *etv2* and *lmo2* were significantly reduced in HIPPOi-treated embryos, while markers of alternative mesenchymal fates *tagln* and *alcam* were significantly upregulated (**Fig. 5d**). Thus, Hippo signaling appears to be important for maximal *etv2* expression in zebrafish embryos, and may control a similar bifurcation between EC and mesenchymal fates *in vivo*.

## Discussion

Complex gene programs unfold at each step of human development, and mapping the regulatory wiring of each emerging cell-type remains a defining challenge of developmental and stem cell biology. Here we exploit the ability to differentiate hPSCs into large numbers of a given cell-type *in vitro* to discover evolving gene programs at each step of differentiation. A benefit of this *in vitro* differentiation system is that one can dynamically watch when and how differentiation becomes derailed when these gene programs are compromised, with temporal resolution that would be challenging to accomplish *in utero*. Analysis of gene programs during human EC differentiation revealed a surprising requirement for the Hippo mechanosignaling pathway. Hippo pathway inhibition suppressed the expression of an upstream transcriptional program controlled by *ETV2*, thus diverting cells away from the EC lineage and towards a mesenchymal fate. Conversely, Hippo pathway activation enhanced EC differentiation. Our results reveal a mechanosignaling pathway at the crux of EC specification and demonstrate how a detailed map of gene programs can enhance stem cell differentiation protocols.

Lateral mesoderm progenitors choose among multiple downstream fates^63^, and differentiation into ECs is thought to be principally instructed by VEGF signaling^37,64^. Given the preeminence of VEGF, it was thus surprising to us that if Hippo signaling is inhibited, lateral mesoderm fails to acquire EC identity even despite the presence of VEGF *in vitro*. Hippo inhibition likewise blunts *etv2* expression in zebrafish embryos. This thus reveals an additional regulatory layer in addition to VEGF, defining an <u>AND</u> logic gate whereby VEGF <u>AND</u> Hippo signaling are simultaneously required to specify ECs (**Fig. 5e**). This may relate to the longstanding curiosity that *Vegf* is widely expressed in the developing embryo^68^, yet vasculogenesis only occurs at discrete locations. It will be interesting to explore whether this contingent requirement for VEGF and Hippo signaling might confine where vasculogenesis occurs. Perhaps the density, size, and shape of lateral mesoderm cells, or the stiffness of their microenvironment, could bias cells towards either an EC or mesenchymal fate. At later stages of development, Hippo signaling allows mature ECs to sense the physical forces of blood flow,^51–55^ but our findings assign an earlier role for Hippo signaling in imparting endothelial identity. This builds on a past observation that *YAP1* knockdown moderately promoted EC differentiation^69^, and expands the role for Hippo signaling in cell fate decisions beyond its known roles in the blastocyst, mesenchyme, intestine, and other developmental contexts.^70–72^

Considerable attention has been devoted to improving stem-cell differentiation through extracellular signaling manipulations^1^, yet we discovered that Hippo signaling levels vary between single lateral mesoderm cells within differentiating cultures (**Fig. S10a**). Such variability in Hippo signaling levels among single cells likely accounts for why pharmacological activation of Hippo signaling improves EC differentiation purity. In any culture, single cells often vary in their mechanical environments and Hippo signaling levels, based on local differences in cell density or position within a cell sheet or colony^73–75^. Perhaps these often neglected mechanical differences might impede directed differentiation, and neutralizing such differences can maximize homogeneity. Modulating Hippo signaling in addition to traditional extracellular signaling pathways thus provides an additional knob to control differentiation heterogeneity *in vitro* that may be generally applicable to other differentiation contexts. Taken together, this demonstrates how CC-Perturb-seq can reveal novel regulators that can be modulated to improve *in vitro* differentiation methods.

Our CC-Perturb-seq experiments elucidate the transcriptional effects of a myriad of other EC differentiation regulators in addition to the Hippo pathway (**Table S2**), including VEGF-associated signaling components such as *PLCG1* and *PRKCD*, BMP pathway components such as *BMPR1A* and *SMAD1/5*, and the primitive-streak transcription factors *EOMES* and *MIXL1*. By organizing single-cell transcriptomes into gene programs with cNMF and mapping their responses to perturbation across differentiation, we connected these regulators to coordinated changes in developmental identity and cellular activity. This program-level view distinguished regulators with similar effects on final EC output but different effects on intermediate cell states and alternative fates. For example, knocking down BMP pathway genes or mesodermal transcription factors (*EOMES* or *MIXL1*) both impaired subsequent EC differentiation, but led to deviations into different types of fates. Thus, a shared failure to acquire endothelial identity can arise through distinct disruptions of differentiation. This screen thus serves as a launching pad to explore additional mechanisms driving EC differentiation by characterizing the dynamic regulatory logic of EC regulators at each stage of differentiation. The CC-Perturb-seq method will also be generally applicable across cell types and systems (*e.g.*, **Fig. S2**), and in parallel studies we have applied it to map regulators in valvular interstitial cells^76^ and keratinocytes.^77^

In summary, this framework for mapping the regulatory wiring of cell differentiation—combining genome-wide genetic screens with high-content molecular mapping—identifies signaling pathways that act during differentiation *in vitro* and development *in vivo*. Expanding this approach to many additional cell types will provide a foundation for understanding human development, understanding the etiology of developmental diseases, and developing improved differentiation protocols for basic research and cell therapy.

## Methods

### 1. Development and validation of CC-Perturb-seq

#### Overview

CC-Perturb-seq was developed to combine the scalability and low per-cell cost of split-pool combinatorial indexing with efficient direct detection of CRISPR guide RNAs. The method adapts the SPLiT-seq workflow of Rosenberg et al.^33^ by including a gRNA-specific reverse-transcription primer during the first round of cellular barcoding. Consequently, Pol III-transcribed gRNAs and cellular mRNAs acquire the same three-round combinatorial cell barcode, permitting perturbation identities and transcriptomes to be linked within individual cells. Following amplification of the barcoded cDNA, gene-expression and gRNA-derived molecules are separated by SPRI size selection. The gRNA-enriched fraction is subjected to strand selection, single-stranded DNA circularization and inverse PCR, thereby enriching guide-derived molecules while retaining their cellular barcodes and unique molecular identifiers. Unlike CROP-seq-based capture^78^, which detects a Pol II-transcribed reporter containing the guide sequence, circular capture directly detects the Pol III gRNA transcript and can be adapted to different gRNA scaffolds or expression vectors by changing the guide-specific reverse-transcription primer.

#### General CC-Perturb-seq protocol

##### Split-seq with direct gRNA capture and circularization

The following general protocol describes the home-built CC-Perturb-seq workflow used in this study, including direct gRNA capture, three rounds of combinatorial indexing and circularization-based enrichment of guide-derived amplicons.

##### Cell fixation and permeabilization

Cells were prepared as single-cell suspensions and maintained on ice during fixation. Cryopreserved samples were thawed at 37 °C, diluted gradually in pre-warmed complete growth medium and centrifuged at 300 × *g* for 5 min. Up to 4 × 10^6^ cells were washed in PBS supplemented with Protector RNase inhibitor (Sigma–Aldrich) and Enzymatics RNase inhibitor and passed through a 40-µm cell strainer. Larger strainers were used where required by cell size.

Cells were fixed in 1% formaldehyde (Thermo Fisher Scientific, 28906) for 10 min on ice and permeabilized with 0.2% Triton X-100 for 3 min on ice. Formaldehyde was quenched with Tris and the cells were washed by centrifugation at 500 × *g* for 5 min at 4 °C. Cells were subsequently treated with 2 mM bis(sulfosuccinimidyl)suberate (BS3; Thermo Fisher Scientific, A39266) for 20 min on ice, followed by a second 3-min permeabilization with 0.2% Triton X-100. The reaction was quenched with Tris, and cells were washed, resuspended in 0.5× PBS containing both RNase inhibitors and filtered again through a 40-µm strainer. DMSO was added gradually to approximately 5% (v/v), and aliquots were frozen at −80 °C using a controlled cooling rate of approximately −1 °C min^−1^.

##### Combinatorial cellular barcoding

For experiments targeting approximately 1 × 10^6^ profiled cells, approximately 4 × 10^6^ fixed cells were loaded across 96 wells for the first round of barcoding. Low-binding plastics, polypropylene tubes and a swinging-bucket centrifuge were used throughout to minimize cell loss. All cell-handling buffers were supplemented with Protector and Enzymatics RNase inhibitors.

The first barcode was introduced during reverse transcription using 96 well-specific phosphorylated primers containing an 8-nucleotide barcode. Each primer mixture contained poly(dT), random-hexamer and direct-capture primers targeting the gRNA transcript. Cells were added to 40-µl reverse-transcription reactions containing Maxima H Minus reverse transcriptase (Thermo Fisher Scientific, EP0753), reverse-transcription buffer and dNTPs. Reverse transcription was performed at 50 °C for 10 min, followed by three cycles of 8 °C for 12 s, 15 °C for 45 s, 20 °C for 45 s, 30 °C for 30 s, 42 °C for 2 min and 50 °C for 3 min, and a final incubation at 50 °C for 5 min.

Cells were pooled, supplemented with Triton X-100 to 0.1% and collected at 200 × *g* for 10 min at 4 °C. The pellet was resuspended in NEBuffer r3.1 containing RNase inhibitors. The second barcode was introduced by ligation in a 96-well plate using pre-annealed barcode–linker duplexes and T4 DNA ligase (New England Biolabs, M0202M). Reactions were incubated at 37 °C for 30 min, followed by addition of a complementary blocking oligonucleotide and a further 30-min incubation at 37 °C.

Cells were pooled, filtered and redistributed into a third 96-well plate containing pre-annealed, 5′-biotinylated barcode–linker duplexes. The third ligation was performed at 37 °C for 30 min and terminated using a third-round blocking oligonucleotide and EDTA. The resulting barcodes comprised three rounds of split-pool indexing. For million-cell-scale experiments, the barcoded pool was divided into at least 16 separately indexed sublibraries, each containing no more than approximately 62,500 cells.

##### Cell lysis, cDNA capture and amplification

After the third barcoding round, cells were washed in PBS containing 0.1% Triton X-100 and RNase inhibitors, counted using YOYO-1 staining and aliquoted into sublibraries. Cells were lysed by adding an equal volume of 2× lysis buffer containing 20 mM Tris, 400 mM NaCl, 100 mM EDTA and 4.4% SDS, together with Proteinase K (Thermo Fisher Scientific, 25530049). Lysates were incubated at 65 °C for 60 min with intermittent shaking and were either processed immediately or stored at −80 °C.

Proteinase K was inactivated with phenylmethylsulfonyl fluoride. Biotinylated barcoded cDNA was captured using Dynabeads MyOne Streptavidin C1 beads (Thermo Fisher Scientific, 65001) for 60 min at 25 °C. Beads were washed in high-salt binding and washing buffer followed by Tween-containing storage buffer. Template switching was performed on bead-bound cDNA using Maxima H Minus reverse transcriptase, the Split_TSO oligonucleotide, dNTPs, PEG 8000 and RNase inhibitor. Reactions were incubated at 25 °C for 30 min and then at 42 °C for 90 min.

Template-switched cDNA was amplified directly from the beads using KAPA HiFi HotStart ReadyMix (Roche, KK2602) and the Split_Partial-TSO and Split_TruSeq-Read2 primers. PCR consisted of five initial cycles using an annealing temperature of 65 °C, followed by 4–12 cycles using an annealing temperature of 67 °C. The number of cycles was selected according to the number of cells and RNA content of each sublibrary, with a target total cDNA yield of 50–200 ng to avoid overamplification.

Amplified material was subjected to a 0.7× SPRI separation. The bead-bound fraction was retained for construction of gene-expression libraries, whereas the supernatant was retained for gRNA-library construction. Gene-expression cDNA was eluted from the 0.7× beads. gRNA-enriched cDNA was recovered from the supernatant by increasing the SPRI ratio to 1.2×.

##### Gene-expression library construction

Up to 50 ng of gene-expression cDNA per sublibrary was fragmented at 32 °C for 10 min and subjected to end repair and A-tailing at 65 °C for 30 min using the Enzymatics WGS fragmentation system. Fragmented cDNA was size-selected using a 0.6×–0.8× double-sided SPRI procedure. Annealed sequencing adapters were ligated at 20 °C for 15 min using WGS ligase, followed by a 0.8× SPRI cleanup.

Each sublibrary was amplified with a unique combination of indexed Split_P5 and Split_P7 primers. Six to twelve PCR cycles were used according to the amount of fragmented cDNA entering the reaction. Libraries were subjected to a final 0.6×–0.8× double-sided SPRI size selection, quantified using a Qubit dsDNA High Sensitivity assay and assessed using a Bioanalyzer. The expected gene-expression library distribution was centred at approximately 400–500 bp.

##### gRNA-library construction by circularization

The gRNA-enriched cDNA fraction was amplified using a feature-specific PCR that introduced a 5′ phosphate on one strand and a 5′ biotin on the complementary strand. PCR was performed using KAPA HiFi HotStart ReadyMix and the Split_5Phos_Read1_TSO and Split_5Biotin_Read2 primers. Five initial cycles with a 65 °C annealing temperature were followed by two or three cycles with a 67 °C annealing temperature. The expected direct-capture product was approximately 296 bp. Products were purified using a 1.2× SPRI cleanup, quantified and pooled across up to eight sublibraries, with approximately 2 µg of PCR product used for each circularization pool.

Biotinylated PCR products were captured on Streptavidin C1 beads. After washing, the non-biotinylated single-stranded DNA was released by two consecutive incubations in freshly prepared 0.2 M NaOH at 37 °C for 5 min. Eluates were neutralized with HCl and Tris and purified using a Zymo DNA Clean & Concentrator-5 column.

Purified single-stranded gRNA DNA was circularized using CircLigase ssDNA ligase (Biosearch Technologies, CL4115K) in the presence of ATP, MnCl₂ and betaine. Circularization was performed at 60 °C for 10 h, followed by enzyme inactivation at 80 °C for 10 min. For all gRNA libraries in this study, the Split_circ_scaffold oligonucleotide was annealed to the circularized product and the resulting duplex was re-linearized with DraI at 37 °C for 1 h, followed by heat inactivation at 65 °C for 20 min. DNA was purified again using a Zymo DNA Clean & Concentrator-5 column.

Final gRNA libraries were amplified using KAPA HiFi HotStart ReadyMix and the Split_circ_gRNA_P5 and Split_circ_gRNA_P7 primers. PCR consisted of five cycles with a 65 °C annealing temperature followed by ten cycles with a 67 °C annealing temperature. The expected final product was approximately 351 bp. Libraries were purified using a 0.9× SPRI cleanup, quantified by Qubit and assessed using a Bioanalyzer.

##### Sequencing

For the general protocol, gene-expression and gRNA libraries were pooled at a molar ratio of 6–10:1 with 5% PhiX. Recommended sequencing depths were approximately 30,000–50,000 gene-expression read pairs per cell and 5,000 gRNA read pairs per cell. The recommended read configuration was 66 cycles for gene-expression Read 1, at least 64 cycles for gRNA Read 1, 8 cycles each for the i7 and i5 indexes, and 86 cycles for Read 2. Experiment-specific achieved or targeted depths are reported in the corresponding subsections.

#### CC-Perturb-seq data processing and analysis

Paired-end gene-expression and guide-enrichment FASTQ files were processed using a custom Snakemake workflow.^79^ Read-level quality was assessed with fastp;^80^ these analyses generated quality-control reports, whereas the original untrimmed FASTQ files were used for quantification.

Libraries were generated using home-built split-pool combinatorial-indexing chemistry. In Read 2, nucleotides 1–10 encoded the unique molecular identifier (UMI), and three 8-nucleotide combinatorial barcode segments occupied nucleotides 11–18, 49–56 and 79–86, separated by invariant linker sequences. Cell barcodes were corrected using the corresponding barcode whitelist and replacement tables. Gene-expression and guide-enrichment libraries were processed independently and linked using sample-specific pairings defined in the sample manifest.

Gene-expression reads were pseudoaligned and quantified using kallisto|bustools.^81^ Quantification was performed using the mature, nascent and ambiguous RNA (nac) workflow, with forward-strand specificity and fractional assignment of reads pseudoaligning to multiple genes (--mm). The transcriptome index and gene annotations were generated from GENCODE release 43 on the GRCh38.p13 genome assembly and included both mature and nascent transcript annotations.^82^ UMI counts assigned to mature, nascent and ambiguous RNA were retained as separate layers and summed to generate the total gene-by-cell count matrix.

Guide-enrichment reads were quantified independently using the KITE workflow^81^ against the study-specific guide library, generating a cell-by-guide UMI count matrix. The guide reference included the sequence and identifier of each targeting and control guide. BUS-file statistics were collected using bustools inspect. Gene-expression and guide-library sequencing saturation was evaluated by repeating quantification after subsampling to 10%, 25%, 50%, 75% and 100% of the available reads.

Before cell calling, barcodes with fewer than 100 total gene-expression UMIs were removed. Cell-associated barcodes were then called independently within each gene-expression sublibrary using the inflection point estimated by barcodeRanks in DropletUtils.^83^ A barcode was classified as cell associated when its total gene-expression UMI count was greater than or equal to the sublibrary-specific inflection-point threshold.

Following cell calling, gene-expression quality control was performed separately for each differentiation day. Cells with total UMI counts or numbers of detected genes outside day-specific lower or upper thresholds defined using four normalized median absolute deviations were excluded. Cells with more than 25% of gene-expression UMIs assigned to mitochondrial genes or more than 5% assigned to annotated ribosomal genes were also removed. Barcode-index groups with consistently lower gene-expression quality were excluded in their entirety; these comprised barcode indices 3 and 16 at D0 and barcode indices 18 and 20 at D1.

Guide-count matrices were aligned to the quality-controlled gene-expression matrices by exact corrected cell-barcode identity. The gene-expression barcode set was treated as authoritative: cells without corresponding guide reads were retained with zero guide counts.

For downstream analyses, guide-to-cell assignment was rerun independently for D0, D1, D2 and D3 from the raw guide-count matrices using the SCEPTRE R package.^84^ Because the experiment was conducted at high multiplicity of infection, SCEPTRE was run in high-MOI mode using its mixture-model guide-assignment strategy. For each guide, this procedure models the guide UMI-count distribution and assigns the guide by thresholding fitted posterior probabilities of guide membership. The resulting binary assignments allowed zero, one or multiple guides to be assigned to an individual cell.

The five-UMI guide calls generated during the initial preprocessing workflow were used only to summarize guide-detection quality-control metrics, including guide UMIs and provisional guide multiplicity per cell. They were not used as the final perturbation assignments. For all downstream knockdown, differential-expression and gene-program analyses, the provisional fixed-threshold matrix was replaced by the SCEPTRE-derived guide-assignment matrix. Raw guide UMI counts were retained alongside the final binary assignments.

Genes were annotated using the GENCODE v43 annotation database. Gene symbols, Ensembl identifiers, gene biotypes, chromosome assignments and HGNC identifiers were propagated to the feature metadata where available. Mitochondrial genes were identified from gene symbols beginning with MT-, and ribosomal and cell-cycle genes were annotated using curated reference lists. Total gene-expression UMIs, number of detected genes, mitochondrial and ribosomal transcript fractions, total guide UMIs and number of assigned guides were calculated for each cell.

Cells were assigned to experimental wells and biological samples by extracting the round-1 barcode from each corrected combinatorial cell barcode, validating this barcode against the corresponding barcode table and mapping it to the experiment-specific plate map. Biological sample, replicate, differentiation day, plate, well and other available plate-map annotations were propagated to the cell metadata.

The resulting AnnData objects contained the total gene-expression count matrix; separate mature, nascent and ambiguous RNA layers; GENCODE v43 gene annotations; cell-level quality-control metrics; raw guide UMI counts; SCEPTRE-derived guide assignments; and sample, replicate, plate and well metadata. Normalization, dimensionality reduction, cell-state annotation and perturbation-effect testing were performed in subsequent analysis steps rather than during this preprocessing stage.

#### Methods for initial experiments to validate CC-Perturb-seq

##### Comparison of circular capture and CROP-seq capture in CRISPRi hiPSCs

To compare direct circular capture with reporter-based CROP-seq capture while holding the cell type and gRNA vector constant, WTC11-CRISPRi hiPSCs carrying a pooled gRNA library expressed from a CROP-seq vector were processed in parallel using the split-pool combinatorial-indexing workflow described above. In the circular-capture condition, a scaffold-specific gRNA reverse-transcription primer was included during the first barcoding round, and the resulting guide-derived cDNA was separated from gene-expression cDNA by SPRI size selection and processed by strand selection, single-stranded DNA circularization and guide-enrichment PCR. In the CROP-seq-capture condition, no direct gRNA reverse-transcription primer was included, and guide identities were recovered from the Pol II-transcribed guide-reporter molecule generated by the CROP-seq vector. Gene-expression libraries were otherwise prepared using the same split-pool barcoding and library-construction workflow.

Gene-expression and guide libraries were processed independently and linked using corrected combinatorial cell barcodes. For comparisons of guide recovery at matched sequencing depth, guide-library reads were subsampled to the same assigned-read depth before UMI quantification. Guide-capture efficiency was summarized as the number of guide UMIs per cell and the fraction of cells with at least one detected guide. Transcriptome complexity was evaluated using total transcriptome UMIs and detected genes per cell. Transcriptome saturation was assessed by repeating quantification after subsampling the gene-expression FASTQ files to 10%, 25%, 50%, 75% and 100% of the available reads and calculating the median transcriptome UMI and detected-gene counts at each depth.

To assess whether improved guide detection affected measurement of CRISPRi activity, on-target expression was evaluated for genes represented in the pilot library. For each target, normalized expression of the intended target gene was compared between cells assigned a corresponding targeting guide and control cells without a guide targeting that gene. Target genes were ordered by control expression for visualization. Statistical significance was corrected across the tested target genes, and associations with an adjusted *q* value of 0.10 or less were highlighted.

##### Validation of circular capture in teloHAECs

The comparison between circular capture and CROP-seq capture was repeated in telomerase-immortalized human aortic endothelial cells carrying gRNAs expressed from a CROP-seq vector. Cells were processed using the same split-pool combinatorial-indexing workflow and the same circular-capture or reporter-capture strategies described for the hiPSC experiment. Because individual cells could contain more than one detected guide, guide-capture efficiency was summarized using the UMI count of the most abundant guide in each cell. Transcriptome recovery was assessed using the total gene-expression UMI count per cell. Gene-expression and guide libraries were processed using the same barcode-correction and quantification procedures used for the hiPSC benchmarking experiment.

##### Compatibility of circular capture with Parse Evercode v3 barcoding

To test whether circular capture could be incorporated into the Parse Evercode v3 split-pool workflow, hiPSCs and K562 cells carrying distinct pooled gRNA libraries were processed using a hybrid Parse–CC-Perturb-seq protocol. The hiPSCs carried a library of approximately 17,000 gRNAs at a nominal multiplicity of infection of approximately 5, whereas K562 cells carried a library of approximately 97,000 gRNAs at a nominal multiplicity of infection of approximately 7. The two cell populations were fixed separately using the Parse Biosciences cell-fixation workflow and were combined at an approximately 1:1 ratio only after fixation.

The mixed fixed-cell suspension was adjusted to approximately 8,400 cells in 14 µl per Round 1 well and loaded into an Evercode WT v3 Round 1 plate. Immediately before reverse transcription, designated wells were supplemented with either the CS1 gRNA-specific RT primer, the DC1 gRNA-specific RT primer, both primers or no gRNA-specific primer. CS1 targeted the CS1 direct-capture sequence, whereas DC1 targeted a sequence present in a broader range of commonly used gRNA scaffolds. Wells without a gRNA-specific primer served as negative controls for guide capture and as controls for evaluating whether the added primers altered transcriptome recovery.

Following Round 1 reverse transcription, cells were processed through the standard Parse WT v3 Round 2 and Round 3 ligation-based barcoding reactions. Cells were then pooled and divided into eight sublibraries containing approximately 13,000 cells each. Two sub-libraries were carried forward for cDNA capture, template switching, amplification and sequencing-library preparation.

After cDNA amplification, the standard Parse post-amplification SPRI purification was modified to separate gene-expression and guide-derived products by size. A 0.7× SPRI cutoff was used to retain the larger gene-expression cDNA, and the shorter guide-derived cDNA in the supernatant was recovered using a 0.7×–1.2× double-sided SPRI selection. The gene-expression fraction was fragmented, end-repaired, adapter-ligated and amplified with a unique dual index according to the Parse WT v3 workflow. The guide-enriched fraction was processed using the downstream CC-Perturb-seq workflow, including guide-specific re-amplification, biotin-based strand selection, isolation of single-stranded DNA, CircLigase-mediated circularization and guide-enrichment PCR. Final gene-expression and guide libraries were assessed by Qubit and Bioanalyzer before pooling and sequencing. The experimental workflow—Parse Round 1–3 barcoding followed by CC-Perturb-seq size fractionation and circular guide enrichment—is documented in the Parse compatibility study. 250912 CC-Perturb-seq with Pars…

##### Processing and analysis of the Parse v3 compatibility experiment

Gene-expression and guide-library reads were initially processed using Parse Biosciences Pipeline v1.4. Gene-expression reads were processed using the whole-transcriptome workflow, and the STAR option was used for guide-reference alignment. Guide UMIs were additionally quantified using UMI-tools. In this analysis, a single read was sufficient to support an observed UMI, after which UMIs with closely related sequences were collapsed. UMI-tools-derived counts were used for the guide-capture comparisons because the Parse guide-counting workflow required at least two reads per UMI and was more conservative at the available guide-library sequencing depth.

A combined reference was constructed from the hiPSC and K562 guide libraries. Guides that could not be assigned unambiguously to one of the two source libraries were excluded from guide-pool-specific analyses, yielding a filtered reference containing 113,275 unique gRNA sequences. Guide counts were summarized as a cell-by-guide UMI matrix and linked to the gene-expression matrix using corrected cellular barcodes.

Gene-expression counts were normalized to 10,000 counts per cell and log-transformed as log(1 + *x*). Highly variable genes were selected using minimum mean expression of 0.0125, maximum mean expression of 3 and minimum dispersion of 0.25. Total transcriptome UMI count was regressed out, values were scaled with a maximum absolute value of 10, and principal-component analysis was performed. A 20-nearest-neighbor graph was constructed using the first 30 principal components, followed by UMAP and Leiden clustering at resolution 0.2. These settings reproduce the analysis workflow used to generate the K562–hiPSC benchmarking panels.

Clusters were assigned as K562 or hiPSC on the basis of lineage-marker expression. K562 identity was evaluated using *GATA1*, *KLF1*, *HBB*, *HBA1* and *HBA2*, whereas hiPSC identity was evaluated using *SOX2*, *NANOG*, *POU5F1* and *LIN28A*. Mean marker scores were calculated from library-size-normalized, log-transformed expression, and each Leiden cluster was assigned to the cell type with the higher mean marker score. Cells without detectable expression of either marker set were designated unassigned and excluded from cell-type-specific guide-pool analyses.

Transcriptome recovery was compared among the no-guide-primer, CS1, DC1 and dual-primer Round 1 conditions using total transcriptome UMIs per cell. Sequencing-saturation curves were obtained from the Parse pipeline subsampling reports by plotting median transcript counts per cell against mean assigned reads per cell.

Guide-capture efficiency was evaluated using three related metrics. First, total guide UMIs and the UMI count of the most abundant guide were calculated for each cell. The top-guide metric was used to compare capture among primer conditions while reducing the dependence of total guide counts on the number of guides carried by each cell. Second, guide UMIs were summed separately over the hiPSC and K562 guide pools. For a transcriptome-assigned K562 cell, K562-pool UMIs were defined as matched-pool guide UMIs and hiPSC-pool UMIs as cross-pool guide UMIs; the definitions were reversed for transcriptome-assigned hiPSCs. Third, guide-pool specificity was calculated for each cell as

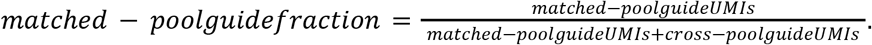

For the guide-pool mixing analysis, individual cell–guide observations supported by fewer than three UMIs were removed before summing guide counts by source library. Total hiPSC-pool and K562-pool guide UMIs were then compared within each cell and related to the independently assigned transcriptomic cell identity. The ≥3-UMI filtering rule and the use of separately fixed cells mixed only before barcoding were intended to distinguish cross-pool signal from low-count sequencing or UMI errors.

### 2. Genome-wide CRISPRi screen to identify and validate regulators of hiPSC-EC differentiation

#### Overview

A genome-wide CRISPRi screen was performed to find essential genes for endothelial cell differentiation. In this screen, the frequencies of CRISPRi guides were compared between successfully differentiated and undifferentiated cells in the post-differentiation population, as endothelial cells can be sorted via MACS based on CD144 expression, an endothelial cell marker. This screen also allows for the comparison of guide frequencies between cell populations before and after hiPSC differentiation into endothelial cells.

By assessing frequency of guides, we can identify genes that may potentially activate or inhibit differentiation: for example, if guides targeting a certain gene in this screen are found at a higher frequency in the unsuccessfully differentiated group compared to the successfully differentiated group, this may indicate that specific gene is involved in an essential process, therefore cells infected with guides targeting that gene will either die or fail to differentiate. Alternatively, guides for genes found to have higher frequencies in the successfully differentiated group may indicate that inhibition of those genes promotes the differentiation process.

WTC11-CRISPRi Human Induced Pluripotent Stem Cells (hiPSCs) were infected using a genome-wide CRISPRi library containing 123,168 guides corresponding to 20,528 target genes^85^, and 6,470 guides as safe and non-targeting controls. Cells were then differentiated over a period of three days into arterial endothelial cells using our published differentiation protocol.^4^ Three groups of cells were collected over this process: infected “baseline” WTC11-CRISPRi cells before differentiation (D0), CD144+ endothelial cells (D3 ECs), and CD144- non-endothelial cells (D3 non-ECs). Analyzing differentiation efficiency via FACS, 71.9% of cells successfully differentiated into endothelial cells.

Two approaches were used to identify and rank hits. First, MAGeCK v0.5.9.5 robust ranking aggregation^86^ was applied to the guide-count data. Of the 20,528 genes targeted, 239 candidate regulators had FDR < 0.05: 233 negative hits, for which targeting guides were enriched in the D3 non-EC fraction, and 6 positive hits, for which targeting guides were enriched in the D3 EC fraction. Second, a Mann–Whitney U-test with Benjamini–Hochberg correction identified 90 negative and 6 positive hits at adjusted P < 0.05. The union of the two analyses comprised 236 negative and 8 positive regulators selected for further validation.

#### WTC11-CRISPRi hiPSC cell culture

WTC11-CRISPRi cells (hiPSCs harboring doxycycline-inducible CRISPRi)^87^ were cultured in 6-well plates (Falcon) coated with Matrigel hESC-Qualified Matrix (Corning) in mTeSR Plus media (STEMCELL Technologies). Plates were prepared by diluting 180 µL Matrigel in 12 mL DMEM/F-12 medium (Gibco), adding 2 mL of diluted Matrigel to each well, and incubating the plates at 37 °C in 5% CO₂ for at least 1 h. Cells were passaged using Accutase (STEMCELL Technologies). At 70–85% confluence, medium was aspirated and 1 mL Accutase was added per well. Cells were incubated at 37 °C for 3 min, diluted with 1 mL room-temperature mTeSR Plus medium, collected and centrifuged at 200 × g for 3 min. Cells were resuspended in 1 mL mTeSR Plus containing 10 µM Y-27632 (ROCK inhibitor) and counted using a Countess instrument (Invitrogen). For routine maintenance, 100,000 cells were transferred to each prepared well in 2 mL medium containing 10 µM Y-27632.

#### sgRNA library design and cloning

To build a genome-wide library, we first obtained the sgRNA protospacer sequences from the Dual sgRNA CRISPRi libraries of Jonathan Weissman’s lab^85^ (Addgene #187246, #187247, #187248). This resulted in 123,168 sgRNAs targeting 20,528 genes (6 sgRNAs/promoter), 2680 safe-targeting and 3790 non-targeting sgRNAs, for a total of 129,638 total guides. A pool of sgRNA oligonucleotides containing the guide protospacer sequence flanked by partial human U6 promoter (TATCTTGTGGAAAGGACGAAACACCG) and partial scaffold (GTTTAAGAGCTATGCTGGAAACAGCATAG) sequences were synthesized by Agilent Technologies.

The guide pool was amplified with a forward primer (GGCTTTATATATCTTGTGGAAAGGACGAAACACCG) and a reverse primer (CTTATTTAAACTTGCTATGCTGTTTCCAGCATAGCTCTTAAAC), extending the Gibson arms for subsequent cloning step into sgOpti-CS vector.^88^ Plasmid backbone was prepared by digesting with BsmBI and ClaI (New England Biolabs), followed by purification with 1× AMPure XP SPRI bead clean-up. Gibson assembly was performed for 1 hour at 50 °C using Gibson Assembly Master Mix (New England Biolabs), 50 ng backbone, and 70 ng of purified PCR-amplified sgRNA pool in a final volume of 40 µL. The cloned plasmid was purified with 0.7× AMPure XP SPRI, electroporated into Endura competent cells (Lucigen #602422), and cells were then expanded in liquid culture for 18 hours at 30 °C. Plasmid library was then obtained with the Nucleobond Xtra Midi EF kit (Macherey–Nagel #740420.50). To sequence-validate sgRNAs and determine their relative abundance, we PCR amplified and sequenced the sgRNAs from the plasmid library on Illumina MiSeq standard v2 50-cycle kit. Analysis of the sequencing data showed that all designed sgRNAs were represented, with a 3.3-fold ratio between the 90th and 10th percentiles of guide abundance.

#### Lentivirus generation

HEK293T cells were seeded at 550,000 cells per well in 6-well plates and incubated for 24 h. A DNA mixture containing 900 ng psPAX2 packaging vector (Addgene #12260), 360 ng pMD2.G VSV-G envelope vector (Addgene #12259) and 1.2 µg purified plasmid library was prepared in 192 µL Opti-MEM and transfected using 5.8 µL X-tremeGENE HP DNA Transfection Reagent (MilliporeSigma, 06366236001). Viral supernatant was harvested 48 h later, passed through a 0.45-µm filter and stored immediately at −80 °C.

#### Lentiviral infection of WTC11-CRISPRi hiPSCs

WTC11-CRISPRi cells were transduced at a multiplicity of infection (MOI) of 1, maintaining a coverage of more than 500 transduced cells per sgRNA. Transduction was performed by adding lentivirus and polybrene (MilliporeSigma, TR-1003-G) to the culture medium at a final polybrene concentration of 8 µg/mL for 24 h. The medium was then aspirated and replaced with medium containing 0.5 µg/mL puromycin (Sigma-Aldrich) and 1 µg/mL doxycycline (Selleck, S5159) for 24 h. This medium was subsequently replaced with fresh medium containing 0.5 µg/mL puromycin and 1 µg/mL doxycycline for an additional 24 h, yielding a total of 48 h of puromycin selection and the first 48 h of CRISPRi induction. Doxycycline was maintained in all subsequent culture and differentiation media to sustain dCas9–KRAB expression.

#### Optimal differentiation seeding density test

Before conducting large-scale experiments, we performed test differentiations to determine the optimal seeding density for the batch of hiPSCs used at the time. We tested seeding densities ranging from 150,000 to 350,000 cells per well in 6-well plates and differentiated the cells into endothelial cells. Based on these results, we selected the seeding density that yielded the highest differentiation efficiency for the large-scale experiments. The cell numbers were then scaled proportionally to the surface area of the culture vessels used in the larger-scale setup.

#### Differentiation and cell harvest

Following puromycin selection and the first 48 h of doxycycline treatment, infected WTC11-CRISPRi hiPSCs were passaged using Accutase and seeded at the optimal density determined above in mTeSR Plus medium containing 10 µM Y-27632 and 1 µg/mL doxycycline. Cells were maintained in this medium for an additional 24 h, completing a total of 72 h of doxycycline-mediated CRISPRi induction before differentiation. Arterial EC differentiation was then initiated, with 1 µg/mL doxycycline maintained in all differentiation media and the medium replaced every 24 h for 3 d. On D3, cells were washed with PBS, incubated with 1 mM EDTA at 37 °C for 15 min and dissociated by trituration in differentiation medium. The resulting cell suspension was processed for flow cytometry and CD144-based magnetic-activated cell sorting (MACS) as described below.

CDM2 media^4^ was prepared using 225 mL IMDM (Life Technologies), 250 mL Ham’s F-12 Nutrient Mix with GlutaMAX Supplement (Life Technologies), 0.7 µg/mL insulin (Sigma), 15 µg/mL transferrin (Roche), 5 mL Chemically Defined Lipid Concentrate (Gibco), 500 U/mL penicillin-streptomycin (Gibco), 450 µM thioglycerol (Sigma), and 1 mg/mL polyvinyl alcohol(Sigma). Detailed instructions to prepare CDM2 are available.^89^

Artery EC differentiation was performed as described previously, with differentiation factors added to CDM2 basal medium. On each day of differentiation, cells were exposed to different media to drive sequential differentiation into primitive streak (day 0-1 interval; 24 hours), lateral mesoderm (day 1-2 interval; 24 hours), and artery ECs (day 2-3 interval; 24 hours). Doxycycline was added to the medium to induce CRISPRi expression.^4^

| D0-1 Media (Primitive streak induction) | D1-2 Media (Lateral mesoderm induction) | D2-3 Media (Artery EC induction) |
|---|---|---|
| 60mL CDM2 media<br>30ng/mL Activin A (STEMCELL)<br>40ng/mL BMP4 (Peprotech)<br>6 µM CHIR99021 (R&D Systems)<br>20 ng/mL FGF2 (R&D Systems)<br>1 µg/mL Doxycycline (Selleck) | 60mL CDM2 media<br>40ng/mL BMP4<br>2.5 µM GDC0941 (Selleck)<br>100 ng/mL VEGF (STEMCELL)<br>1 µM XAV939 (R&D Systems)<br>10 µM Forskolin (R&D Systems)<br>2 µM SB505124 (Selleck)<br>200 µg/mL AA2P (Cayman Chemicals)<br>1 µg/mL Doxycycline | 60mL CDM2 media<br>15ng/mL Activin A<br>2.5 µM GDC0941<br>100 ng/mL VEGF<br>1 µM XAV939<br>200 µg/mL AA2P<br>1 µg/mL DMH1 (STEMCELL)<br>1 µg/mL Doxycycline |

#### Flow cytometry

One million cells were set aside for flow cytometry to confirm successful differentiation. Cells were centrifuged at 300 × *g* for 3 minutes, supernatant aspirated, and pellet resuspended in 270 µL PBS + 1% BSA. Each sample was then equally split into three FACS tubes. In the first tube, an isotype antibody mix was added to 1:50 dilution for APC-iso (BioLegend) and 1:500 dilution for FITC-iso (BioLegend)). In the second tube, a targeting antibody mix was added to 1:100 dilution for APC-CD144 (BioLegend) and 1:200 dilution for FITC-CD31 (BioLegend)). In the third tube, PBS + 1% BSA was added to serve as an unstained control. All tubes were covered and incubated at 2–8 °C for 30 minutes. Cells were centrifuged and washed with PBS + 1% BSA. This washing step was repeated again, and the final product was resuspended in 300 µL PBS + 1% BSA. We also included hiPSCs as negative control and TeloHAEC as positive control for antibody staining. These samples were then analyzed using the MA900 Multi-Application Cell Sorter (Sony) to determine the frequency of APC-CD144 and FITC-CD31 double positive population.

#### MACS sorting

Cells were magnetically labeled with CD144 MicroBeads (Miltenyi Biotec 130-097-857) according to the manufacturer’s instructions. Briefly, the day 3 cell suspension was centrifuged at 300 × *g* for 5 minutes, then resuspended with 80 µL MACS buffer per 10 million cells. Then 20 µL of MicroBeads were added and incubated at 2–8 °C for 15 minutes. After incubation, 1 mL MACS buffer was added to wash and centrifuged again at 300 × *g* for 10 minutes. Cell pellet was resuspended in 500 µl MACS buffer and magnetic separation of cells was performed within LS Column (Miltenyi Biotec) on a MACS Separator and MultiStand (Miltenyi Biotec). CD144+ labeled ECs and non-labeled cells (non-endothelial cells) were collected and centrifuged at 300 × *g* for 5 minutes. Supernatant was carefully aspirated and cell pellets were frozen at −20 °C for later genomic DNA extraction.

#### Genomic DNA extraction and amplicon sequencing

Extraction of genomic DNA for each sorted cell sample was performed using the QIAamp Blood Maxi Kit (QIAGEN) according to the manufacturer’s spin protocol with a maximum of 20 million cells per spin column. Following extraction, each genomic DNA sample was split into separate PCR replicates in a 96-well plate (Eppendorf) with each PCR reaction containing 6 µg of genomic DNA. Genomic region containing the gRNA was amplified using Q5 High-Fidelity 2× Master Mix (New England Biolabs, no M0492L) and 0.5 µM of each amplification primer containing 8 bp sample index and Illumina adapters: AATGATACGGCGACCACCGAGATCTACACNNNNNNNNCGATTTCTTGGCTTTATATATCTTGTG; CAAGCAGAAGACGGCATACGAGATNNNNNNNNCGGTGCCACTTTTTCAAGTTG. Genomic amplicon PCR was carried out with the following thermocycling parameters: 98 °C for 3 min and then 28 cycles of 98 °C for 10 s, 62 °C for 20 s, 72 °C for 25 s, followed by a final 72 °C extension for 2 min. Equal volumes of PCR replicates were pooled together and DNA was purified by a double-sided SPRI bead cleanup (Beckman) (first, a left-handed cleanup at 0.5×, then a right-handed cleanup at 1.2×). 5 µL of each sample was then run on a 2% agarose gel (Invitrogen) to confirm a clear band of 210 bp. DNA concentration was measured using the Qubit 1× dsDNA HS Assay Kit (Invitrogen, no Q33231). DNA amplicon libraries were sequenced on an Illumina Nextseq 550 System (High Output v2.5, 75-cycle kit) using a custom sequencing primer: CGATTTCTTGGCTTTATATATCTTGTGGAAAGGACGAAACACCG for Read 1 and AAGGCTAGTCCGTTATCAACTTGAAAAAGTGGCACCG for Index 1.

#### Data Analysis

Raw sequencing reads were demultiplexed using the bcl2fastq program (Illumina) to generate FASTQ files and trimmed to contain the unique gRNA sequences. These processed reads were then aligned to the designed gRNA sequences from the library using ^90^ to obtain read counts for each gRNA in each sorted sample.

We calculated the gene fold change for each gene based on the frequency of gRNA reads in the CDH5+ versus CDH5- populations. This was calculated as follows:

1. Calculate the fold change of control guides between CDH5+ and CDH5- populations. Safe-targeting and non-targeting guides were used to estimate the control fold change between the CDH5+ and CDH5− fractions. This control factor accounted for global shifts in library composition caused by perturbations that altered survival or differentiation. For each biological replicate, fold changes for the safe-targeting and non-targeting guide sets between the CDH5+ and CDH5− fractions were calculated. The mean of these two control fold changes was used to normalize target-gene effects.

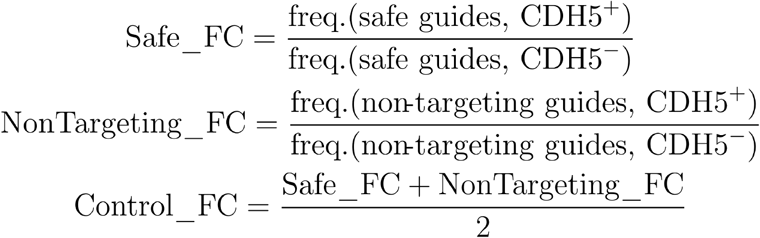
2. Calculate the fold change for all target genes between CDH5+ and CDH5− populations.
  a. For each gene, the frequency of guides targeting that gene in the CDH5+ population was divided by the frequency within the CDH5- population.
  b. To normalize this value to control guides, the value in (a) was then divided by the control guide fold change in Step 1.
  c. The base-2 logarithm of the normalized fold change was then calculated. 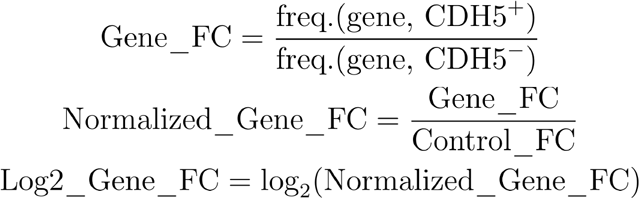

To calculate the P value and adjusted P value for each gene between the CDH5+ and CDH5− fractions, a Mann–Whitney U-test with Benjamini–Hochberg correction^91^ was performed on the guide-frequency ratios for all guides targeting each gene versus the ratios for all negative-control guides.

### 3. CRISPRi screen to validate regulators of hiPSC differentiation into arterial ECs

#### Overview

The focused validation screen contained the 239 genes significant by MAGeCK together with five additional genes identified by the Mann–Whitney U-test but not by MAGeCK, 30 randomly selected non-hit genes expressed at >20 TPM in at least one differentiation time point, and 23 additional expressed non-hit genes with reported roles in endothelial biology, proliferation, cancer or cell-cycle regulation. The final panel contained 297 target genes. The 1,926-sgRNA library included 1,458 guides for the 244 candidate regulators, 180 guides for the 30 random non-hits, 168 guides for the 23 additional non-hits, and 60 safe-targeting plus 60 non-targeting controls. Depending on the number of targeted promoters, genes were represented by 6–12 sgRNAs.

Screen quality was assessed by flow cytometry and by reproducibility of gene-level guide-enrichment effects. The two biological replicates yielded 84.4% and 72.1% CD144⁺/CD31⁺ endothelial cells, respectively (Fig. S1b). Gene-level effects were reproducible between the validation-screen replicates (R² = 0.85; **Fig. S1c**) and between the genome-wide and validation screens (Pearson’s *r* = 0.72; **Fig. S1d**).

#### Cell culture, sgRNA library cloning and lentivirus production

Unless noted otherwise, all procedures were performed as described above for the genome-wide screen. WTC11-CRISPRi hiPSCs were cultured as described above. The validation sgRNA library was synthesized and cloned into the sgOpti-CS vector as described above. Lentivirus was produced as described above, except that HEK293T cells were seeded on plates coated with 50 µg/mL poly-D-lysine (Sigma Millipore), virus was produced in three 6-well plates, and viral titer was measured using Lenti-X GoStix (Takara).

#### Lentiviral infection

Two biological replicates were infected independently. For each replicate, 11 Matrigel-coated 15 cm plates (Falcon) were seeded with 2,265,625 WTC11-CRISPRi hiPSCs per plate (equivalent to 150,000 cells per well of a 6-well plate; 22 plates and ∼48 million cells in total) in mTeSR Plus + 10 µM RI and incubated for 24 hours. Cells were then transduced with lentivirus diluted 1:1,280 in mTeSR Plus with polybrene (266 µL virus and 204 µL polybrene in 340 mL media; 23 mL per plate) for 24 hours. Puromycin selection and doxycycline induction were performed as described above (30 mL media per plate), except that puromycin + doxycycline media was replaced daily for two days.

#### Differentiation and cell harvest

For each biological replicate, infected hiPSCs were seeded in four Matrigel-coated 10 cm plates (Falcon) at 2.1 million cells per plate in 12 mL mTeSR Plus + 10 µM RI + 1 µg/mL doxycycline. After 24 hours, two plates per replicate were dissociated with Accutase, counted and pelleted as the “baseline” (pre-differentiation) sample, and pellets were stored at −20 °C for gDNA extraction. The remaining two plates were differentiated and harvested as described above (12 mL media per plate). After harvest, cells were resuspended in 5 mL PBS + 1% BSA, passed through a 70 µm cell strainer (Falcon) and counted.

#### Flow cytometry, MACS sorting, library preparation, and data analysis

For each biological replicate, 1 million cells were analyzed by flow cytometry and the remaining cells were MACS-sorted on CD144 as described above. Genomic DNA was extracted, and sgRNA amplicons were amplified, purified and sequenced as described above, with the following changes: each gDNA sample was split into seven 100 µL PCR reactions, each containing < 2.5 µg gDNA, 50 µL Q5 2× Master Mix (NEB) and 5 µL each of unique Illumina i5/i7 primers, with thermocycling at 98 °C for 30 s; 28 cycles of 98 °C for 15 s, 62 °C for 15 s and 72 °C for 16 s; and 72 °C for 2 min. 10 µL of each reaction was pooled by condition and biological replicate (six samples) before the double-sided SPRI cleanup. Data analysis was conducted as for the genome-wide screen.

### 4. CC-Perturb-seq to characterize regulators of hiPSC differentiation into arterial ECs

#### Target selection and gRNA library design

The CC-Perturb-seq panel matched the focused validation screen and comprised the 239 MAGeCK-significant genes plus 58 additional genes: five genes identified only by the Mann–Whitney U-test, 30 randomly sampled non-hit genes expressed at >20 TPM in at least one differentiation time point, and 23 additional expressed non-hit genes with reported functions in endothelial biology, proliferation, cancer or cell-cycle regulation. The final panel comprised 297 target genes.

The pooled library contained 1,926 sgRNAs: 1,806 targeting guides distributed across the 297 genes, with 6–12 sgRNAs per gene depending on the number of targeted promoters, together with 60 safe-targeting and 60 non-targeting control sgRNAs. Guide sequences and cloning followed the focused validation-library design described above. Briefly, pooled oligonucleotides containing each protospacer flanked by partial human U6 promoter and gRNA-scaffold sequences were synthesized, PCR-amplified to add Gibson-assembly homology arms and cloned into the sgOpti-CS vector.^92^ The assembled library was electroporated into Endura competent cells, expanded in liquid culture and purified by midi preparation. Plasmid-library representation was assessed by amplicon sequencing before lentivirus production, and the resulting pooled lentiviral library was used for the CC-Perturb-seq experiment.

#### Cell culture

WTC11-CRISPRi hiPSCs harboring doxycycline-inducible dCas9–KRAB were maintained on hESC-qualified Matrigel in mTeSR Plus and passaged using Accutase, as described above. Two biological replicates were transduced independently with pre-titered focused-library lentivirus at high multiplicity of infection and subjected to puromycin selection. The delivered MOI was calibrated by droplet digital PCR. CRISPRi was induced with 1 µg mL⁻¹ doxycycline for 3 d before differentiation, and doxycycline was maintained in all subsequent culture and differentiation media.

Following selection and CRISPRi induction, 200,000 cells were seeded per well of a 6-well plate and differentiated using the same three-day arterial EC differentiation protocol described above, with differentiation medium replaced every 24 h. Cells were collected immediately before differentiation as the D0 hiPSC sample and subsequently every 24 h. D1, D2 and D3 corresponded to primitive streak, lateral mesoderm and arterial endothelial-cell stages, respectively. The two biological replicates were cultured, differentiated and harvested independently.

#### CC-Perturb-seq library preparation and sequencing

Cells from each biological replicate were harvested at D0, D1, D2 and D3, yielding eight biological samples. At each time point, cells were dissociated into single-cell suspensions and fixed, permeabilized and stored according to the general CC-Perturb-seq protocol above. All eight samples were multiplexed during Round 1 barcoding, with non-overlapping reverse-transcription wells assigned to each replicate–time-point sample. Within each replicate, Round 1 wells were allocated across D0:D1:D2:D3 in a 2:1:1:2 ratio. Reverse transcription included poly(dT), random-hexamer and guide-specific capture primers. Cells were then pooled and processed jointly through the second and third ligation-based barcoding rounds.

Following completion of the three barcoding rounds, the pooled cells were divided into three independently indexed sublibraries containing approximately 50,000 cells each. Cells were lysed, and biotinylated barcoded cDNA was captured on streptavidin beads, template-switched and PCR-amplified as described above. A 0.7× SPRI separation partitioned amplified material into a bead-bound gene-expression fraction and a gRNA-enriched supernatant fraction. Gene-expression libraries were generated by fragmentation, end repair, A-tailing, adapter ligation and indexed PCR. Guide-derived cDNA was amplified by feature-specific PCR using the guide-specific RT product, strand-selected, circularized with CircLigase, re-linearized with DraI and amplified by inverse PCR using sublibrary-specific indexed primers.

Gene-expression and gRNA libraries were pooled at a molar ratio of approximately 6–10:1 with 5% PhiX. Gene-expression libraries were sequenced to approximately 10,000 paired-end reads per cell using 150 cycles for Read 1, 8 cycles for each index read and 150 cycles for Read 2 on a NovaSeq X flow cell. Guide libraries were sequenced to approximately 2,000 paired-end reads per cell.

#### Data processing and analysis

Gene-expression and guide-enrichment FASTQ files were processed using the canonical CC-Perturb-seq pipeline described under “CC-Perturb-seq data processing and analysis.” For this experiment, the pipeline used the home-built barcode structure, GENCODE release 43 on GRCh38.p13 and the experiment-specific guide reference. Cell calling and day-specific gene-expression quality control followed the thresholds described above. Sample, biological replicate and differentiation day were recovered from Round 1 barcodes and the plate map. Guide assignment was rerun independently for D0–D3 using SCEPTRE high-MOI mixture-model assignment; fixed five-UMI calls were retained only for guide-detection quality control. The final AnnData objects contained total, mature, nascent and ambiguous RNA counts; feature annotations; cell-level quality-control metrics; raw guide UMI counts; SCEPTRE guide assignments; and experimental metadata.

#### On-target knockdown analysis

Differential gene-expression testing and estimation of on-target knockdown were performed using SCEPTRE v0.10.3^84^. Analyses were conducted independently for D0, D1, D2 and D3 using the raw gene-expression UMI matrix and raw guide UMI matrix. Data were imported using the SCEPTRE high-MOI model, and guide-to-cell assignments were inferred with the mixture-model assignment method. For the on-target analysis used to calculate knockdown efficiency, discovery pairs were constructed by matching each intended gRNA target to the corresponding GENCODE v43 response gene. Guides targeting the same gene were combined using the union integration strategy; thus, a cell was classified as perturbed for a target when at least one guide assigned to that target was present. The control group for each target consisted of the complement of this set—that is, cells without an assigned guide targeting the gene being tested, including cells carrying guides targeting other genes.

Tests were two-sided and used SCEPTRE’s conditional randomization test with the skew-normal resampling approximation and the default high-MOI adjustment model. These covariates were selected for adjusting the analysis model: the logarithm of the number of detected response genes, the logarithm of total gene-expression UMIs, the logarithm of the number of detected guides, the logarithm of total guide UMIs and biological-sample identity as a batch covariate:

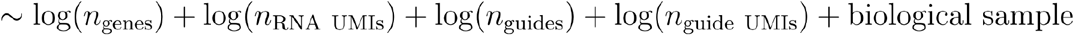

SCEPTRE pair-level quality control used a mitochondrial-transcript threshold of 0.25, followed by the calibration check and discovery analysis. Covariates were selected to yield well-calibrated null tests, as assessed from calibration Q–Q plots. Only response–target pairs passing SCEPTRE quality control were retained. Conditional-randomization P values were adjusted by the Benjamini–Hochberg procedure separately within each day across all quality-control-passing on-target pairs; BH-adjusted P ≤ 0.05 was considered significant.

For each target and time point, relative target-gene expression was obtained from the SCEPTRE fold_change estimate, with a value of 1 representing expression in the corresponding complement-control population.

Thus, a fold change of 0.40 corresponded to an estimated knockdown efficiency of 60%, whereas negative knockdown values indicated estimated expression above the control reference. Fold-change standard errors were taken directly from the SCEPTRE v0.10.3 se_fold_change output. Where confidence intervals were displayed, approximate 95% Wald intervals were calculated on the log₂-fold-change scale by applying the delta method to the SCEPTRE fold-change estimate and its standard error; these model-based intervals were reported separately from the conditional-randomization *P* values.

#### Defining a high-confidence regulator set

Among the 239 genes assigned as hits in the genome-wide CRISPRi screen, we defined 174 as confident regulators based on validation in this CC-Perturb-seq experiment. Confident regulators are those where (1) perturbing the gene changed the fraction of endothelial cells on Day 3 in CC-Perturb-seq (Fisher’s exact test, BH FDR < 0.05; **Table S2**), in the same direction as in the genome-wide screen, and (2) the RefSeq-curated transcription start sites (TSSs) did not include another TSS within 1 kb that had its own genome-wide screen effect in the same direction and at least as large (|log2 fold-change| ≥ that of the regulator).

### 5. Gene program inference and annotation

#### Gene program inference

Gene programs were inferred using a GPU-accelerated implementation of consensus Non-negative Matrix Factorization (cNMF; https://github.com/ymo6/torch_based_cNMF), adapted from the original CPU implementation.^28^ In place of the coordinate descent solver used in the original package, our implementation employs a hierarchical alternating least squares (HALS) solver^93^ for parallelizing efficient NMF factorization on GPU (https://github.com/ymo6/nmf-torch).

Unnormalized, post-QC mRNA UMI counts were used as input to the cNMF pipeline, which performs TP10K depth normalization and per-gene unit-variance scaling before NMF. The 121,616 cells from D0–D3 were concatenated into one expression matrix. No highly variable gene filter was applied; all 17,538 expressed genes were used. Factorization was performed in full-batch mode on one NVIDIA H100 80-GB GPU. For each candidate K, 100 NMF replicates were initialized with distinct random seeds. We evaluated K = 10, 20, 30, 40, 50, 60, 70, 80, 90, 100, 150, 200, 250 and 300. K = 50 was selected from the combined stability, reconstruction-error, explained-variance and biological-enrichment metrics shown in Fig. S4a, at the point where additional components provided diminishing biological and statistical gains. The selected solution used a consensus density threshold of 2.0.

The resulting gene-program matrix had dimensions 50 programs × 17,538 genes, with each program representing a set of coordinately regulated genes. The corresponding 121,616 cells × 50 programs usage matrix represented each cell state as a weighted combination of programs capturing both cellular identity and cellular activity.

#### Gene program evaluations

Each of the 50 gene programs in the selected K = 50 solution was evaluated for statistical performance and enrichment of biological signals using standardized tasks described previously in Schnitzler et al.^34^ The per-program evaluations included:

● Reconstruction error between the original and reconstructed matrices, measured with Frobenius loss
● Stability, measuring the consistency of inferred gene sets across replicates with silhouette score
● The amount of variance in normalized count data explained by each gene program
● Gene-set enrichment using the top-loaded genes ranked by their score within a gene program
● Unique regulators for each gene program, identified by comparing program scores of perturbed cells to non-targeting controls

The final set of gene programs (K=50) was selected at the inflection point where the curves for the total number of enriched gene sets and the total number of unique regulators per program set began to plateau while remaining relatively stable (**Fig. S4**).

#### Measure reconstruction error

Reconstruction error quantifies how far the inferred cell usage (W) and gene program (H) matrices recapitulate the original expression matrix (X). We compute this using the Frobenius norm of the residual, which sums the squared deviations across all gene-by-cell entries.

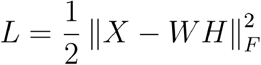

#### Assessing stability among NMF replicates

Because NMF is a non-convex optimization problem, different initializations can converge to different local minima, yielding variable gene programs across replicates. To quantify the consistency of programs recovered across initializations, we use the silhouette score,^94^ which measures how tightly replicate programs cluster together relative to programs from other components. Higher silhouette scores indicate that a given program is reliably recovered across independent runs, while low or negative scores flag unstable components that may reflect initialization artifacts rather than robust biological signals.

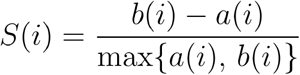

#### Computing variance explained per gene program

To quantify the contribution of each program to expression variability, we computed a dimensionally consistent rank-one reconstruction for each program following Schnitzler et al.^34^ Let X ∈ Rⁿˣᵖ denote the normalized cell-by-gene matrix and let hₖ ∈ R₊¹ˣᵖ denote the k-th program row vector. The least-squares per-cell usage coefficient was

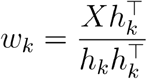

The rank-one reconstruction using program k alone was

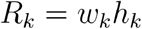

where wₖ ∈ Rⁿ and Rₖ ∈ Rⁿˣᵖ. The fraction of variance explained by program k was calculated as

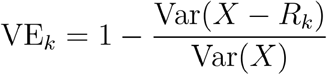

where Var(·) was evaluated over all cell-by-gene entries. Larger VEₖ values indicate that the program captures a greater share of transcriptional variation.

#### Gene-set enrichment per program

For each program, we tested the top 300 program genes for enrichment against curated gene-set libraries using a one-sided Fisher’s exact test. The background set for each test comprised the union of all genes in the library excluding the gene-set under evaluation, ensuring that enrichment was assessed relative to the library’s gene universe rather than the full transcriptome. We queried two libraries accessed through the Enrichr catalogue^95^ via gseapy^96^, Reactome 2022^97^ for pathway enrichment and GO Biological Process 2023^98^ for Gene Ontology enrichment. Within each program, p-values across all gene-sets were adjusted for multiple testing using the Benjamini–Hochberg procedure to control the false discovery rate (FDR), and gene-sets were considered significantly enriched at FDR < 0.05.

#### Identification of gene program regulators

To identify regulators whose perturbation altered program usage, analyses were performed separately at D0–D3 using the K = 50 cNMF usage matrix selected at density threshold 2.0. Exact zero usages were replaced by the day-specific floor and rows were renormalized; program outcomes were then subjected to the internal quantile floor and centered log-ratio transformation described below. For each target, all assigned guides were combined using a union indicator. The adjustment model included percent mitochondrial counts, biological sample, total RNA UMI counts, guides per cell, guide UMI counts and detected genes. Count-like covariates were transformed as log(1 + x), biological sample was dummy-encoded, constant columns were removed, retained numeric columns were standardized and an intercept was included. The observed target–program effect was the ordinary least-squares coefficient of the target indicator. Target propensities were estimated using L2-regularized logistic regression, and two-sided empirical P values were obtained from 10,000 Bernoulli conditional-randomization samples using a plus-one correction. Storey q values were calculated globally across all target–program tests from D0–D3, and q < 0.05 was considered significant.

#### Gene program regulator calibration

To evaluate CRT calibration, we constructed frequency-matched NTC pseudo-targets by grouping safe-targeting and non-targeting guides into six-guide sets matched to the guide-frequency-bin signatures of real targets. Each pseudo-target was tested over the same full day-specific cell population and with the same CLR outcomes, covariates, propensity-model procedure and 10,000 Bernoulli resamples used for real targets. The analysis was not restricted to cells carrying only control guides.

We compared NTC pseudo-target P values with expected uniform quantiles using quantile–quantile (Q–Q) plots. Under adequate calibration, NTC pseudo-targets should follow the identity line, whereas real targets are expected to show an excess of small P values when true perturbation effects are present.

#### CRT implementation details

##### Overview

We tested whether CRISPR perturbation of each target regulator was associated with altered usage of transcriptional programs identified by consensus non-negative matrix factorization (cNMF). The analysis was performed separately at each iPSC-to-endothelial-cell differentiation time point (d0, d1, d2, and d3), using the K = 50 cNMF solution selected at density threshold 2.0. For a given day, N denotes the number of retained cells, K the number of programs, and G the number of guide RNAs. The analysis used the empirical, two-sided Bernoulli conditional randomization test (CRT). The CRT follows the model-X conditional-independence framework, in which the perturbation indicator is resampled from an estimated conditional distribution given measured covariates while the observed outcomes and covariates are held fixed.

##### Program-usage preprocessing and centered log-ratio outcome

For cell i and program k, let u_ik_^raw^ ≥ 0 denote the cNMF usage value. Because program usage is compositional and rows are normalized to sum to one, program effects were modeled after a centered log-ratio (CLR) transformation. The current iPSC-EC implementation applies two sequential flooring operations before the CLR transformation.

First, exact zeros in each day-specific usage matrix are replaced by a data-adaptive floor. Let Q+0.01 denote the first percentile computed over strictly positive usage entries for that day. The initial floor is

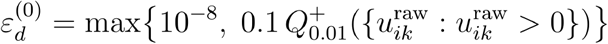

Only exact zero entries are replaced at this stage. The resulting values are row-normalized:

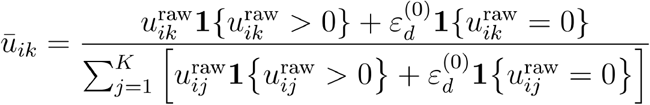

Second, programDE applies its internal quantile floor before taking logarithms. In the current implementation, the quantile level is eps_quantile = 10^−4^, so

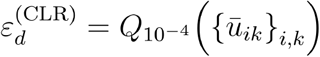

followed by

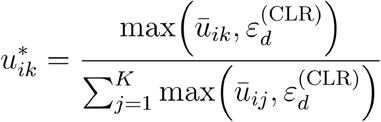

The response for program k in cell i is then

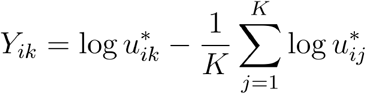

Thus, an estimated perturbation coefficient is interpreted on the CLR scale: it represents a change in the log ratio of one program to the geometric mean of all programs, rather than an absolute change in an unconstrained expression measurement. The realized values of both floors are data dependent and can differ by day. The fixed implementation settings are the 10^−8^ lower bound, the 0.1 multiplier applied to the first percentile of positive usages, and the internal 10^−4^ usage quantile.

##### Perturbation indicators and covariates

Guide assignments were converted to a binary cell-by-guide matrix A, with Aij = 1 when guide j was detected in cell i. For target regulator g, all guides mapped to that regulator were combined into a union perturbation indicator

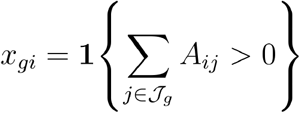

where J_g_ is the set of guides targeting g. A cell was therefore considered treated for target g if at least one guide assigned to g was present; cells carrying multiple guides targeting the same regulator were counted once in the union indicator.

The six-source covariate specification used for the full-day iPSC-EC analysis comprised percent mitochondrial counts (pct_counts_mt), biological sample (biological_sample), total RNA UMI counts (total_counts), guides per cell (guides_per_cell), guide UMI counts (guide_umi_counts), and the number of genes detected per cell (n_genes_by_counts). Count-like variables were transformed as log(1 + x), whereas percent mitochondrial counts were retained on their original scale. The biological sample was represented by indicator variables with one reference level omitted. Constant columns were removed within each day, all retained numeric columns were standardized, and an intercept was included in the model matrix. Consequently, the modeled covariate vector contained

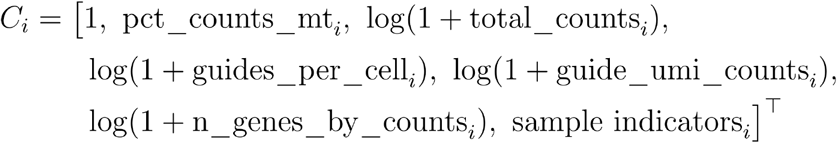

after standardization of the non-intercept columns.

##### Observed effect statistic

For each target g and program k, the observed effect was the ordinary least-squares coefficient of the target union indicator in the covariate-adjusted linear model

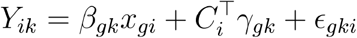

The null hypothesis was the conditional-independence statement

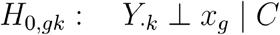

and the test statistic was the absolute value of the estimated target coefficient. Equivalently, define the covariate residual-maker matrix

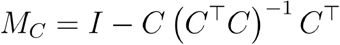

Then the observed coefficient can be written as

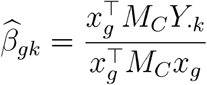

A positive coefficient indicates that the program increased relative to the geometric mean of the remaining programs after covariate adjustment; a negative coefficient indicates a relative decrease.

##### Bernoulli conditional randomization test

For each target regulator, programDE estimated the conditional probability of receiving at least one guide for that target using an L2-regularized logistic regression:

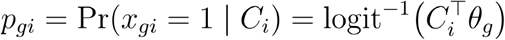

The current implementation uses the lbfgs solver, inverse regularization strength C_logistic_ = 1, and a maximum of 200 iterations. Estimated probabilities are clipped to the interval [10^−6^, 1 − 10^−6^] to avoid numerical degeneracy.

Conditional on the fitted propensity scores, B = 10,000 synthetic perturbation vectors were generated independently under the null:

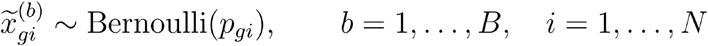

For each synthetic vector, the same covariate-adjusted OLS coefficient was recomputed for every program, yielding a null coefficient for each target-program pair. The empirical two-sided CRT p-value was

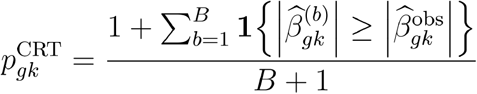

The plus-one correction^99^ gives a valid finite-resampling p-value and prevents zero p-values. With B = 10,000, the minimum attainable empirical p-value is 1/10,001 ≈ 9.999 × 10^−5^.

##### Efficient Bernoulli resampling with the index sampler

A direct implementation would require an N × B binary matrix of Bernoulli draws, which is computationally inefficient. programDE instead uses an exact sparse index sampler. For each cell i, the number of null resamples in which the cell is assigned to the treated group is first drawn as

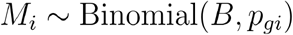

Conditional on M_i_, the sampler chooses M_i_ distinct resample identifiers uniformly without replacement from {1, …, B}:

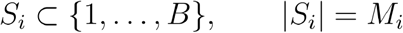

The synthetic treatment is set to one for resamples in Si and zero otherwise. This procedure is distributionally identical to drawing B independent Bernoulli variables for cell i. For any binary pattern z with m treated resamples,

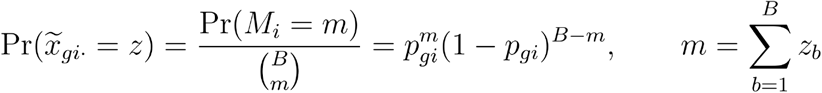

which is exactly the probability of that pattern under B independent Bernoulli(pgi) draws. Sampling is independent across cells.

The selected cell indices are stored in a compressed sparse representation. An indptr array identifies the segment corresponding to each resample, and an indices array stores only the cell indices treated in that resample. This avoids storing the dense N × B matrix and is particularly efficient when target propensities are small.

The OLS statistic is also evaluated without repeatedly solving a regression. The implementation precomputes

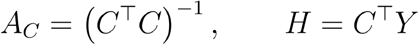

For an observed or resampled treatment vector z, define

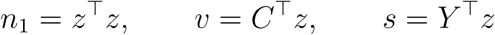

The vector of treatment coefficients across all K programs is then

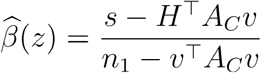

where division is by the scalar denominator. Thus, each null replicate requires sums over the treated cell indices and matrix-vector operations with precomputed quantities, rather than a new least-squares fit for every target-program-resample combination.

##### Construction of frequency-matched NTC pseudo-targets

The non-targeting-control (NTC) calibration curve was constructed from synthetic NTC pseudo-targets. These are computational guide groups rather than biological pseudogenes. Importantly, each NTC pseudo-target was tested over the same full day-specific cell population, using the same CLR outcomes, covariates, propensity-model procedure, Bernoulli sampler, and B = 10,000 resamples as the real targets. The analysis was not restricted to NTC-only cells; cells carrying an NTC guide together with one or more real-targeting guides remained in the common analysis population. Both non-targeting and safe-targeting guides are used and grouped together as NTC guides.

##### Guide frequencies and frequency bins

For guide j, its cell-level frequency was

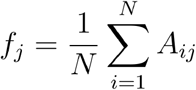

The frequency distribution of real-targeting guides, rather than the NTC guides, was used to define L = 10 quantile bins. Let b(j) denote the frequency-bin assignment of guide j. The same real-guide-derived bin edges were then applied to both real and NTC guides. This makes the bin labels comparable across guide classes and prevents the NTC frequency distribution from defining its own matching scale.

##### Real-target bin signatures

A guide frequency signature was constructed for each real target. Guides for a target were sorted by increasing frequency. For target g with six targeting guides, the signature was

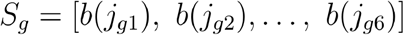

The signature records the multiplicity of guide-frequency bins represented among the six guides. For example, a signature [1, 1, 3, 4, 4, 7] requires two guides from bin 1, one from bin 3, two from bin 4, and one from bin 7. Similar guide frequency signatures were constructed for targets with more than six guides.

##### Frequency-matched NTC group construction

Within each ensemble replicate, NTC guides were shuffled separately within frequency bins. A real-target signature was sampled at random, and an NTC group was formed by taking the required number of currently unused NTC guides from each bin specified by that signature. If the available NTC guides could not satisfy a sampled signature, another signature was attempted. Selected guides were removed from the within-ensemble pools, so an NTC guide was used at most once within a given ensemble. Group construction continued until no additional feasible six-guide groups could be formed; max_groups=None was used, so groups were not deliberately truncated. This procedure was preferred over drawing guides with replacement as this allows all frequency-matched NTC guides to be used.

Ten independently randomized ensemble replicates were generated (n_ensemble = 10) using initial group-construction seed 7, with the ensemble index added to the seed. NTC guides can therefore reappear in different ensemble replicates, but not in multiple pseudo-targets within the same ensemble. The ensemble procedure reduces dependence on any single arbitrary partition of the NTC guide set and samples pseudo-targets across the empirical distribution of real-target frequency signatures.

For NTC group q with guide set Qq, the pseudo-target union indicator was

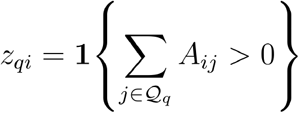

A separate propensity model for the NTC union indicator was fit for each group, followed by the same Bernoulli index-sampler CRT and the same two-sided OLS statistic used for real targets. The base seed for NTC-group CRT resampling was 23, with a group-specific seed derived from the ensemble and group identifiers.

##### NTC pseudo-target QQ calibration curve

For each NTC group, the CRT produced one empirical p-value for each of the K = 50 programs. P-values were concatenated across all NTC groups, all programs, and all ten ensemble replicates. If the pooled collection contains m p-values and p(1) ≤ … ≤ p(m) are their ordered values, the expected uniform quantile for rank r is

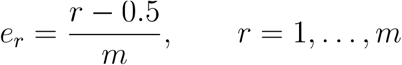

The NTC curve plots

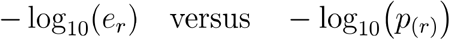

The real-target curve was constructed analogously from all non-NTC target-program p-values, using its own number of points when calculating expected quantiles. Under adequate calibration, the NTC pseudo-target curve should lie near the identity line. Upward deviation indicates an excess of small p-values, whereas downward deviation indicates conservative p-values.

The NTC curve was used as a negative-control calibration diagnostic; it was not used to transform, replace, or otherwise recalibrate the empirical p-values of real targets. Because NTC guides are reused across ensemble replicates and p-values across programs share cells and covariates, the pooled QQ points are correlated. The curve should therefore be interpreted as a descriptive assessment of null behavior rather than as a set of independent observations for a formal goodness-of-fit test.

##### Current implementation parameters

| Component | Current value | Implementation role |
|---|---|---|
| **Differentiation time points** | d0, d1, d2, d3 | Analyzed separately |
| **cNMF solution** | K = 50 | Number of program outcomes per day |
| **cNMF density threshold** | 2.0 | Selected usage solution |
| **CRT resamples** | B = 10,000 | Empirical Bernoulli null draws per target or NTC group |
| **Empirical p-value resolution** | 1/(B + 1) = 1/10,001 | Minimum attainable p-value |
| **Real-target CRT base seed** | 123 | Default gene-specific resampling seed base |
| **Propensity model** | L2 logistic; lbfgs; C = 1; 200 iterations | Estimates P(x = 1 \| C) |
| **Propensity clipping** | [10<sup>-6</sup>, 1 − 10<sup>-6</sup>] | Numerical stability |
| **Initial zero floor** | max(10<sup>-8</sup>, 0.1 × first percentile of positive usages) | Replaces exact zero cNMF usages before row normalization |
| **Internal CLR floor** | 10<sup>-4</sup> quantile of preprocessed usage | Floors small usages before CLR and renormalization |
| **Covariate sources** | Pct_counts_mt; biological_sample; total_counts; guides_per_cell; guide_umi_counts; n_genes_by_counts | Six-source adjustment model |
| **Count-covariate transform** | log(1 + x) | Applied to all count-like covariates |
| **NTC labels** | safe-targeting; non-targeting | Both safe-targeting and non-targeting guides are treated as NTC |
| **NTC group size** | 6-12 | Inferred from guide list to match the real-target’s number of guides |
| **Guide-frequency bins** | 10 quantile bins | Defines real-target signatures and NTC matching |
| **NTC ensembles** | 10 | Independent random matched groupings |
| **NTC group seed** | 7 | Initial group-construction seed |
| **NTC CRT seed** | 23 | Base resampling seed for NTC groups |
| **NTC maximum groups** | None | Uses all feasible non-overlapping groups within each ensemble |

##### Zero-replacement-floor sensitivity analysis

We repeated CRT using initial zero-replacement floors equal to 0.1-fold, 1-fold, or 10-fold the current day-specific value while holding all other analysis settings fixed. Alternative floors primarily changed the numerical scale of CLR beta estimates but preserved effect rankings, directions, major discoveries, and NTC calibration, supporting the current floor while indicating that absolute beta magnitudes should be interpreted as conditional on the flooring choice.

### 6. Characterizing enhancers and transcription factor motifs during hiPSC differentiation into arterial ECs

#### Overview

We annotated gene programs with predicted transcription factor motifs by (i) collecting 10x Multiome data at four hiPSC–arterial EC differentiation time points (D0, D1, D2 and D3); (ii) linking enhancers to target genes using the scE2G model^36^; (iii) identifying motif instances in each enhancer using ChromBPNet^35^ followed by TF-MoDISco^100^ v2.0.7 and Fi-NeMo^101^; (iv) calculating gene–TF motif scores by summing the signed products of motif-hit coefficients and scE2G scores across linked peaks; and (v) performing one-sided Mann–Whitney U-tests comparing the top 300-loaded genes in each program with the remaining genes.

#### 10x Multiome data collection

WTC11 hiPSCs were differentiated using the arterial EC differentiation protocol described above and collected at five consecutive time points, D0–D4, in five biological replicates, yielding 25 replicate–time-point samples. Samples were cryopreserved after collection and subsequently processed together for paired single-nucleus chromatin-accessibility and gene-expression profiling. Each replicate–time-point sample was labeled with a unique cell-multiplexing oligonucleotide (CMO), enabling biological-replicate and differentiation-time-point identities to be recovered after pooling. Although samples were collected at D0–D4, only D0–D3 were used for the scE2G, ChromBPNet and gene-program analyses presented in this study, matching the four-stage CC-Perturb-seq differentiation time course.

Cryopreserved samples were thawed at 37 °C, diluted into pre-warmed medium and centrifuged at 200 × g for 3 min. D0 hiPSCs were recovered in mTeSR Plus, whereas samples from subsequent differentiation time points were recovered in CDM2 medium. Cells were washed twice in PBS containing 0.04% BSA, resuspended in the same buffer and passed through a 70-µm Flowmi cell strainer. Up to 5 × 10^5 cells from each sample were used for nuclei isolation.

Cells were centrifuged at 300 × g for 5 min at 4 °C, resuspended in 100 µl chilled 0.5× lysis buffer and incubated for 2 min on ice. The lysis buffer contained 10 mM Tris-HCl, pH 7.4, 10 mM NaCl, 3 mM MgCl2, 0.1% Tween-20, 0.1% Nonidet P-40 substitute, 0.01% digitonin, 1% BSA, 1 mM DTT and 1 U µl−1 RNase inhibitor. Lysis was terminated by adding 1 ml chilled wash buffer containing 10 mM Tris-HCl, pH 7.4, 10 mM NaCl, 3 mM MgCl2, 0.1% Tween-20, 1% BSA, 1 mM DTT and 0.2 U µl−1 RNase inhibitor. Nuclei were immediately centrifuged at 500 × g for 5 min at 4 °C and resuspended in 180 µl wash buffer.

Nuclei from each sample were labeled separately using an anchor–barcode–co-anchor CMO hashing strategy. Each nuclei suspension was mixed with 20 µl of the corresponding 10× CMO anchor–barcode solution and incubated on ice for 5 min, followed by addition of 20 µl 10× co-anchor solution and a second 5-min incubation on ice. Labeled nuclei were washed with 1 ml chilled wash buffer, centrifuged at 500 × g for 5 min at 4 °C, resuspended in 300 µl wash buffer and passed through a 40-µm Flowmi cell strainer. Nuclei were centrifuged again, resuspended in 200 µl wash buffer and counted. Equal numbers of nuclei from the 25 samples were pooled, using the sample with the lowest recovery to determine the pooling depth; approximately 16,000 nuclei were contributed by each sample. The pooled nuclei were concentrated and divided across two Chromium channels for paired single-nucleus ATAC and gene-expression profiling using the Chromium Next GEM Single Cell Multiome ATAC + Gene Expression workflow.

ATAC and gene-expression libraries were generated according to the 10x Genomics Multiome workflow, with additional modifications to recover CMO-derived products. During cDNA amplification, 1 µl of a 2.5 µM hashtag-oligonucleotide additive primer was added to each reaction, and cDNA was amplified for five cycles. Amplified cDNA was subjected to a 0.6× SPRIselect separation. The bead-bound, mRNA-derived cDNA fraction was processed using the standard 10x gene-expression library workflow, whereas the supernatant containing the shorter CMO-derived products was retained for CMO-library preparation.

CMO products were recovered by increasing the final SPRIselect ratio to 3.2× in the presence of 1.8 volumes of isopropanol, followed by two washes with 80% ethanol and elution in Buffer EB. The recovered CMO products were quantified using Qubit and assessed using an Agilent Bioanalyzer High Sensitivity assay. CMO libraries were indexed by PCR using the same Dual Index TT Set A used for the corresponding gene-expression libraries. PCR was performed for 12 cycles, followed by a 1.6× SPRIselect purification and an additional 1× left-sided SPRIselect cleanup. ATAC libraries were indexed using Single Index Kit N Set A. Final ATAC, gene-expression and CMO libraries were assessed by Qubit, Bioanalyzer and KAPA library quantification. Gene-expression and CMO libraries were sequenced together on an Illumina NextSeq instrument, producing mixed GEX–CMO FASTQ files that were separated computationally as described below. ATAC libraries from the two Chromium channels were sequenced separately on an Illumina NextSeq instrument.

#### 10x Multiome data processing and analysis

##### Library demultiplexing

Gene-expression (GEX) and Cell Multiplexing Oligo (CMO) reads for each channel were delivered in the same FASTQ files. CMO reads were separated from GEX reads with splitcode v0.31.677^102^ by matching Read 2 against a panel of 25 CMO barcodes with Hamming distance ≤2. Matching read pairs were written to CMO FASTQs and all remaining pairs to GEX FASTQs. The 3′ 28 bp of each CMO Read 1, containing the cell barcode and UMI, were retained for quantification.

##### Quantification

Gene expression was quantified using kallisto|bustools (kb-python)^81^ against GENCODE v43, using the nac workflow. CMO tags were quantified per channel with the kb-python KITE workflow (kb-python v0.29.1), indexing the 25 CMO barcode sequences and counting against the 10x multiome cell barcode onlist.

##### scRNA QC

Barcodes were filtered based on UMIs (>=1000 and <=10000), >=200 genes detected, percentage of mitochondrial counts (<=median +2.5 Median Absolute Deviations (MADs)). Doublets were called with Scrublet^103^ (scanpy v1.12.2^104^), and Leiden clusters^105^ in which >40–50% of cells were doublets by both Scrublet^103^ and CMO hashing were excluded.

##### Sample demultiplexing

CMO counts were centered log-ratio (CLR)-normalized per cell, and a CMO was called positive if its CLR value exceeded the 95th percentile of that CMO’s own distribution. Barcodes positive for exactly one CMO were singlets and assigned to their respective timepoints, positive for more than one CMO were doublets and excluded, positive for none were negatives. We rescued negatives by assigning the timepoint based on their Leiden cluster’s majority vote.

Dimensionality reduction and clustering (scRNA): Counts were normalized with PFlog (scclr v0.1.0)^106^. The top 2,000 highly variable genes were used for PCA, followed by k-nearest-neighbor graph construction, Leiden clustering and UMAP^107^ using scanpy v1.12.2.

##### scATAC-seq preprocessing and quality metrics

Fragments were imported with snapATAC2 (v2.9.0)^108^ and restricted to the set of QC-d barcodes from RNA-seq modality. Per cell fragment count and TSS enrichment were calculated using ArchR^109^ (createArrowFiles, minTSS=0, minFrags=0). Aggregate TSS enrichment profiles were computed by treating both ends of each fragment as 1 bp insertion sites, and counting strand aware distances to all TSSs within 2,000 bp in 1 bp bins, using 20531 TSSs (scE2G CollapsdGeneBounds.hg38.intGENCODEv43, deduplicated by gene symbol, chr1-22, X and Y). Insertions near more than one TSS are counted once per TSS. Each profile was divided by its own flanking background, the mean insertion density over the outer 100 bp on each side so that flanks equal 1, and the curve indicates enrichment displayed as an 11 bp centered rolling mean (plotTSSEnrichment with default 200, 100 and 11 bp default values, ArchR).

#### Linking enhancers to target genes with scE2G

Candidate cis-regulatory element (CRE)-to-gene links were identified from 10x Multiome data for each time point following the scE2G pipeline^36^ (https://github.com/EngreitzLab/scE2G). Peaks were called using pseudobulk fragment files with MACS2^110^ (-f BED -g hs -p 0.1 --shift -75 --extsize 150 –nomodel --keep-dup all --call-summits). The 150,000 strongest peaks per time point were extended 250 bp upstream and downstream from their summits to generate uniform 500-bp peaks using makeCandidateRegions.py from the ABC model.^111^ Peaks overlapping blacklisted regions^112^ were excluded, and ±500-bp TSS-proximal regions were included, yielding 156,236 peaks at D0, 156,216 at D1, 156,073 at D2 and 154,835 at D3. For each time point and CRE–gene pair, scE2G scores were quantile-normalized across CREs and high-confidence links were retained at E2G.Score.qnorm ≥ 0.177.

#### Training ChromBPNet on four-time-point 10x Multiome data

##### 1. Data pre-processing

The peak sets generated by scE2G were used as input to ChromBPNet. Following the ChromBPNet preprocessing guidelines^35^ and using bedtools v2.30.0 slop and intersect^113^, blacklisted regions were extended by 1,057 bp, overlapping peaks were removed and peaks were reformatted to narrowPeak format. This yielded 156,186 peaks at D0, 156,163 at D1, 156,022 at D2 and 154,787 at D3. For each time point and fold, GC-content-matched nonpeak background regions were generated with chrombpnet prep nonpeaks using predefined train-validation-test folds.

##### 2. Tn5 bias model training

All models were trained using 5-fold cross-validation. Given the time course experimental design, and to reduce computation, we assumed that the Tn5 bias is timepoint-independent. Therefore, we trained a Tn5 sequence bias model on D0 (iPSC) ATAC data using a deduplicated pseudobulk fragment file.

Bias models were trained across a range of bias threshold factors (0.5, 0.6, 0.7, 0.8, 0.9, 1.0) for each of the 5 cross-validation folds using chrombpnet v1.0.1 bias pipeline, and selected the bias threshold was 0.8 for folds 0, 1 and 3, 0.7 for fold 2, and 0.9 for fold 4.

##### 3. Full ChromBPNet model training

We trained 20 bias-factorized ChromBPNet models using the default architecture across four time points and five folds. The chrombpnet pipeline received the time-point-specific pseudobulk fragment file through -ifrag and the corresponding fold-specific bias model. The bias-corrected models were used for all downstream analyses. Per-nucleotide contribution scores were computed for all peaks and each time-point–fold combination with chrombpnet contribs_bw and averaged across the five folds.

##### 4. Identifying transcription factor motif instances with ChromBPNet

For each time point, TF-MoDISco^100^ was applied to fold-averaged ChromBPNet contribution scores (modisco motifs -n 500000 -w 500). Patterns were clustered and annotated across time points with MotifCompendium using motif_compendium.py --ref-db MotifCompendium-Database-Human.meme.txt --threshold 0.95 [https://github.com/kundajelab/MotifCompendium]. Each motif cluster was assigned a tf_motif_id using the matched TF name for a single reference match or the compendium annotation for a multi-TF match. Fold-averaged contribution profiles were extracted in 1,000-bp windows using finemo extract-regions-chrombpnet-h5 --region-width 1000. Motif instances were called with finemo call-hits -l 0.8 -b 500 and filtered within each motif and time point at cwm_similarity ≥ 0.8. This yielded 731,482 hits for 162 motifs at D0, 893,216 hits for 167 motifs at D1, 1,063,546 hits for 170 motifs at D2 and 1,167,947 hits for 165 motifs at D3.

#### Calculating a score for motif enrichment in each gene program

To identify TF motifs enriched in scE2G peaks linked to the top 300-loaded genes in each cNMF program, we used TF-MoDISco pattern signs, Fi-NeMo hit_coefficient_global values and scE2G enhancer-to-gene predictions (scE2G.Score.qnorm). Only enhancer elements with isSelfPromoter=False were included.

For each timepoint d we used only the Fi-NeMo motif instances that matched the reference CWM with cwm_similarity >=0.8, and, among overlapping instances (>3bp overlap), kept the one with the highest hit similarity. For the downstream analysis, only positive TF-MoDISco patterns were used. Next, for each timepoint d we computed a gene-TF motif score representing the cumulative contribution of a given TF motif across all scE2G peaks linked to a given gene. In particular, the score was defined as follows:

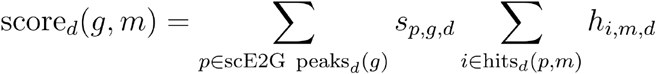

Here, *s_p_*_,*g*,*d*_ represents the quantile-normalized peak to gene score from scE2G, which captures how likely this peak *p* is to regulate this gene *g*. ℎ*_i_*_,*m*,*d*_ corresponds to the hit_coefficient_global (Fi-NeMo output) for an instance *i* of a motif *m* at timepoint *d*. This score reflects specificity of TF’s CWM towards explaining the local contribution score at a given instance after competition with other motifs.

Next, for each cNMF program, we used the top 300 loaded genes. For each motif m and timepoint d, we selected the genes carrying the motif in at least one of the scE2G linked peaks. To test whether a TF motif was preferentially associated with the program’s top loaded genes, we performed a one-sided Mann-Whitney U test. We tested whether gene-TF motif scores were higher for motif carriers (min 10 carriers) among the top 300 genes, as compared to all remaining genes carrying the motif (min 10 carriers). P-values were adjusted for multiple testing using the Benjamini-Hochberg (BH) procedure at FDR of 0.05.

### 7. Effects on EC differentiation percentage and statistical analysis

D3 cells were classified as belonging to an EC or non-EC transcriptional state according to the *CDH5*-based cell-state annotation used in the analysis. For each guide, the D3 EC percentage was calculated as the proportion of guide-carrying D3 cells classified as EC. Cells carrying both the targeting guide being evaluated and any control guide were excluded, ensuring that targeting-guide and control groups were mutually exclusive; target carriers containing other targeting guides remained in the analysis. Gene-level EC percentages were summarized as the unweighted arithmetic mean across all guides targeting that gene. The EC differentiation effect was calculated relative to 120 control guides, comprising 60 safe-targeting and 60 non-targeting guides, as

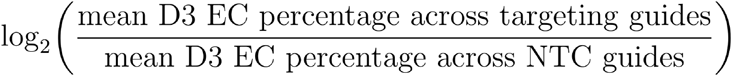

Statistical significance was assessed using a Welch’s t-test comparing the mean EC percentage across all guides targeting each gene with that of the control guides. P values were adjusted across all 297 tested genes using the Benjamini–Hochberg procedure. Significance was denoted as * FDR < 0.05, ** FDR < 0.01 and *** FDR < 0.001. Each guide was treated as one replicate and was not weighted by its cell count.

EC proportion on day 3, Fisher exact test: For each target gene, D3 cells were divided into target-carrier and target-specific control groups. The target-carrier group included cells carrying at least one sgRNA targeting the tested gene. The control group included cells carrying at least one non-targeting control sgRNA but no sgRNA targeting the tested gene. Cells carrying both target and non-targeting sgRNAs were therefore retained in the target group and excluded from the control group. The numbers of arterial endothelial cells and non-endothelial cells in the two groups were compared using a two-sided Fisher’s exact test. Raw P-values were adjusted across all tested genes using the Benjamini–Hochberg procedure.

### 8. Carrier-seeded neighborhood enrichment analysis

For each perturbed gene, target-carrier seed cells were defined as cells carrying at least one guide targeting that gene and no control guide. The target-specific control population comprised cells carrying at least one control guide and no guide targeting the tested gene. Cells carrying both the tested target and a control guide were excluded from both groups. Cell-state neighborhoods were defined independently within each day using the 300 nearest cells selected from all same-day cells in Euclidean distance across z-scored usage of the 50 cNMF programs. The seed cell itself was excluded from its neighborhood. UMAP coordinates were used only to display the resulting enrichment scores and were not used to define neighbors.

For each seed-cell neighborhood, we counted target carriers and target-specific control cells and compared their local ratio with the corresponding ratio among all cells from the same day. Local carrier enrichment was calculated as

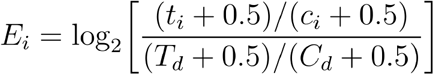

where tᵢ and cᵢ are the numbers of target carriers and control cells within neighborhood i, and T_d and C_d are their total same-day counts. Positive values indicate local enrichment of target carriers relative to their same-day background frequency, whereas negative values indicate depletion. Statistical significance was assessed using a two-sided Fisher exact test comparing target and control counts inside versus outside each same-day neighborhood, followed by Benjamini–Hochberg correction across seed-cell neighborhoods for each target. Because seed-cell neighborhoods overlap, these adjusted P values were interpreted as descriptive local-visualization diagnostics rather than as independent biological replicates.

For visualization, all cells were shown in gray on the integrated D0–D3 UMAP, and target-carrier seed cells were overlaid at their own UMAP coordinates. Point color represents local carrier enrichment, point size represents the fraction of neighboring cells carrying the tested perturbation, and a black outline indicates neighborhood FDR (<0.05). Shared UMAP limits and color scales were used across targets to support direct comparison.

### 9. Pharmacological manipulation of Hippo signaling during hPSC differentiation

#### Human Pluripotent Stem Cell Lines

Wild-type H1 hESCs^114^ and NCRM1 *ETV2^mCherry^* reporter hiPSCs^65^ have been described previously . In NCRM1 *ETV2^mCherry^* reporter hiPSCs, the stop codon in the endogenous *ETV2* locus was replaced with a *P2A-NLS-mCherry* reporter cassette, leaving the coding sequence of *ETV2* intact^65^.

All cells were cultured in standard incubator conditions (37 °C and 20% O_2_/5% CO_2_).

#### hPSC Culture

H1 hESCs were maintained as described previously.^89^ In brief, cells were cultured on TC-treated plates coated with Geltrex (Thermo Scientific, A4000046802) in mTeSR Plus medium (STEMCELL Technologies, 05825) supplemented with 1% penicillin/streptomycin (Fisher Scientific 15-140-163) (henceforth referred to as “mTeSR Plus”).

NCRM1 *ETV2^mCherry^* reporter hiPSCs were initially maintained as described previously^65^ in TeSR/E8 medium (STEMCELL Technologies, 05990) supplemented with 1% penicillin/streptomycin on TC-treated plates coated with Vitronectin (Thermo Fisher, A14700). They were then transitioned into mTeSR Plus medium supplemented with 1% penicillin/streptomycin on Geltrex-coated plates.

Undifferentiated cultures of H1 and NCRM1 *ETV2^mCherry^*hPSCs were maintained by changing mTeSR Plus media every two days. When partially confluent, hPSCs were dissociated into clumps with Versene (Thermo Scientific, 15040066), gently resuspended in mTeSR Plus medium, and replated onto Geltrex-coated plates.

#### Pharmacological modulation of the Hippo pathway during EC differentiation

hPSCs were differentiated into arterial ECs over 3 days as described previously.^4,89^ Seeding density impacts arterial EC differentiation efficiency^89^, and therefore to create a suboptimal baseline against which improvements could be detected, H1 hPSCs were seeded at 5.3 × 10⁴ cells/cm², equivalent to 2 × 10⁵ cells per well of a 12-well plate.

Hippo pathway inhibition was achieved using 1 μM TDI-011536 (Selleck Chemicals, E1314; HIPPOi), an inhibitor of LATS1 and LATS2 kinases.^58^ Functional activation of Hippo output was achieved using 1 μM NVP-IAG933 (Selleck Chemicals, E1490; HIPPOa), which inhibits YAP/TAZ–TEAD interactions.^59^

#### Modulation of surface stiffness during EC differentiation

First, *ETV2^mCherry^* hPSCs were seeded onto 10-cm dishes at 50,000 cells/cm² (2.805 × 10⁶ cells per dish), and then differentiated into primitive streak cells for 24 hours.^4,115^ Primitive streak cells were dissociated with Accutase (Thermo Fisher Scientific 00-4555-56) for 10 minutes at 37 °C, diluted 1:10 in DMEM/F-12, centrifuged at 500 × *g* for 5 minutes, and then resuspended in lateral mesoderm induction media^4^ lacking the PI3K inhibitor GDC-0941. Cells were then seeded on standard 6-well plates, or alternatively, 6-well plates containing PDMS gels ranging from 0.5 to 32 kPa stiffness (Advanced Biomatrix, 51907EA), at a density of either 250,000 cells per well (2.632 × 10^4^ cells/cm², referred to as “Sparse” in the text) or 750,000 cells/well (7.895 × 10^4^ cells/cm², referred to as “Dense” in the text). After 24 hours of lateral mesoderm induction, ETV2^mCherry^ expression was measured via flow cytometry as described below.

#### Flow cytometry

Cells were dissociated with 200–400 μL of TrypLE Express (Thermo Fisher, 12604013) per well for 10 minutes at room temperature. Cells were lightly triturated, and then 50 μL of cell suspension was transferred to a 96-well V-bottom plate (Thermo Fisher, 249570). 150 μL of FACS buffer (comprising PBS [Thermo Fisher, 10010049] + 2% FBS [Thermo Scientific A5670801] + 1% Penicillin/streptomycin + 0.2 mM EDTA [Thermo Fisher, AM9260G]) was added to each well to neutralize the TrypLE. Cells were centrifuged at 500 × *g* for 5 minutes, and then washed with 200 μL FACS buffer.

Two separate procedures were then used, depending on cell genotype. Cells derived from the ETV2^mCherry^ hPSC line were resuspended in FACS buffer containing 0.1 μg/mL DAPI (Thermo Scientific, D1306), and flow cytometry was performed. Cells derived from the H1 hPSC line were stained with an anti-CD144 FITC conjugated antibody (BD Biosciences, 560411), which was diluted 1:50 in FACS buffer. Antibody staining was performed for 30 minutes at 4 °C. After staining, cells were diluted with FACS buffer, centrifuged, washed with FACS buffer, and then resuspended in FACS buffer containing 0.1 μg/mL DAPI.

Flow cytometry was performed on a Beckman Coulter CytoFlex Analyzer. Data analysis was performed using FlowJo. Cells were first identified by gating along the forward scatter area and side scatter area axes. Doublets were eliminated by gating on forward scatter height and width. Live cells were identified as DAPI negative. To determine which cells were positive for ETV2^mCherry^, a negative control (undifferentiated ETV2^mCherry^ hPSCs) was used. To determine which cells were positive for CD144, a negative control (undifferentiated H1 hPSCs) was used.

#### Statistical analysis of Hippo-modulated arterial EC differentiation

For the differentiation-window experiment shown in **Fig. 3e**, Day 3 CD144-positive percentages were analyzed at the level of independently treated culture wells. The analysis included 33 evaluable wells representing eight conditions, including control, across four culture plates in two independent differentiation batches. All measured values were retained, including all six control wells. The four plotting-only zero values for Day 0–1 HIPPOi were excluded because these cultures died and CD144-positive purity was not measurable; they were not treated as measured zero percentages. Missing measurements were not imputed.

Percentages were analyzed on their original scale using an ordinary least-squares model equivalent to an additive two-factor ANOVA, with treatment condition and plate block as categorical fixed effects: purity ∼ condition + plate_block. Plate blocks were defined by the unique combination of differentiation batch and plate identifier, thereby accounting for plate-specific baseline differences within and between batches. No condition-by-plate interaction was included. The model used a common residual variance and had 22 residual degrees of freedom.

Each of the seven evaluable treatment conditions was compared with the shared control using contrasts obtained from the fitted model. Two-sided P values and simultaneous 95% confidence intervals were calculated using a single-step multivariate-t adjustment based on the fitted contrast covariance matrix and residual degrees of freedom, corresponding to a model-based Dunnett procedure. All seven treatment-versus-control comparisons were treated as one family, with family-wise error controlled at 0.05. Descriptive percentages represent observed well-level means; adjusted effects represent model-estimated treatment-minus-control differences in percentage points

#### Quantitative PCR

We performed RNA extraction, reverse transcription, and quantitative PCR (qPCR) as described previously.^115^ Briefly, cells were lysed using 350 or 700 μL of RLT Plus buffer then processed using the RNeasy Mini Plus Kits (Qiagen, 74136) to obtain RNA. 150–300 ng of RNA was reverse-transcribed into cDNA using the High Capacity cDNA Reverse Transcription Kit (Thermo Fisher, 4368813). cDNA was then diluted in Ultrapure DNase/RNase Free Distilled Water (Thermo Fisher, 10977023) at a 1:20–1:40 ratio. 10 μL qPCR reactions were prepared in a MicroAmp Optical 384-well plate (Thermo Scientific, 4326270) using 5 μL of SensiFAST SYBR Green Lo-ROX master mix (Bioline, BIO-94020), 4.6 μL of diluted cDNA, and 0.4 μL of gene-specific forward and reverse primer mix (5 μM). qPCR was performed on a QuantStudio 5 Real-Time PCR machine (Applied Biosystems, A34322). Gene expression was normalized to the reference gene *YWHAZ*.

#### Immunocytochemistry and imaging

Day 2 lateral mesoderm cells were washed once with PBS (Thermo Fisher 10010049) then fixed for 10 minutes at room temperature in 4% paraformaldehyde (Thermo Fisher Scientific PI28908). Cells were then washed for five minutes three times with PBS then permeabilized with permeabilization/blocking solution (0.1% Triton X-100 [Millipore Sigma T8787] + 5% donkey serum [Sigma-Aldrich D9663] in PBS) for one hour at 4 °C. Primary antibodies were diluted in permeabilization/blocking solution at specified dilutions (**Table S7**) and added to cells. Following an overnight incubation protected from light at 4 °C, cells were washed three times with PBS before secondary antibody dilutions (1:500 in permeabilization/blocking solution) were added. Following a one hour incubation at 4 °C, cells were washed twice with PBS then stained with 0.2 μg ml^−1^ DAPI + PBS for 10 min at room temperature. Finally, cells were washed twice with PBS then coated in a working volume of PBS while imaging. Imaging was performed using an Olympus FV3000 confocal microscope.

#### Preparation of Hippo-modulated cells for scRNA-seq

H1 hPSCs were differentiated towards arterial ECs for 3 days in 12-well plates. For each biological condition, three independent replicate wells were seeded on separate plates to account for plate-associated seeding variability. Separate replicate wells were harvested at each time point; cells were dissociated with TrypLE Express, neutralized with FACS buffer and centrifuged for fixation.

Single-cell suspensions were fixed and permeabilized using the Evercode Low Input Cell Fixation v3 kit (Parse Biosciences) according to the manufacturer’s instructions. Cells were counted, resuspended in Cell Prefixation Master Mix and passed through the supplied plate strainer. Cell Fixative Master Mix was added, and samples were incubated for 10 min on ice. Permeabilization Solution was then added for 3 min on ice, after which fixation and permeabilization were terminated by addition of Cell Fix and Perm Stop Master Mix. Fixed cells were frozen at −80 °C under controlled cooling and stored until library preparation.

Fixed cells were subsequently processed using the home-built split-pool combinatorial-indexing workflow described above for CC-Perturb-seq, with two modifications: no gRNA-specific reverse-transcription primer was included, and no gRNA-enrichment or circularization library was generated. Samples representing different treatments, collection times and biological replicates were assigned to non-overlapping wells during the first barcoding round. For Round 1 barcoding, fixed cells were distributed across a 96-well reverse-transcription plate.

Gene-expression libraries were sequenced on an Illumina NovaSeq X flow cell, using 150 cycles for Read 1, 150 cycles for Read 2, 8 cycles for Index 1, 8 cycles for Index 2, to an achieved depth of approximately 22,000 read pairs per cell. No guide RNA library was prepared or sequenced for this experiment.

#### Data processing and analysis of scRNA-seq dataset of Hippo-modulated cells

Paired-end gene-expression FASTQ files were processed using the same custom Snakemake workflow used for the CC-Perturb-seq gene-expression data. The guide-enrichment, KITE quantification, guide-to-cell assignment and SCEPTRE components of the CC-Perturb-seq workflow were omitted. Read-level quality was assessed with fastp, whereas the original untrimmed FASTQ files were retained for quantification.

Libraries were decoded according to the home-built split-pool read structure. In Read 2, nucleotides 1–10 encoded the unique molecular identifier (UMI), and three 8-nucleotide combinatorial barcode segments occupied nucleotides 11–18, 49–56 and 79–86, separated by invariant linker sequences. Cell barcodes were corrected using the corresponding barcode whitelists and replacement tables. Treatment condition, collection time, biological replicate, plate and well identity were recovered by mapping each corrected Round 1 barcode to the experiment-specific plate map. Gene-expression reads were pseudoaligned and quantified using kallisto|bustools with the mature, nascent and ambiguous RNA workflow, forward-strand specificity and fractional assignment of reads pseudoaligning to multiple genes. The transcriptome index and gene annotations were generated from GENCODE release 43 on the GRCh38.p13 genome assembly and included mature and nascent transcript annotations. UMI counts assigned to mature, nascent and ambiguous RNA were retained as separate layers and summed to generate the total gene-by-cell count matrix.

Before cell calling, barcodes with fewer than 100 total gene-expression UMIs were removed. Cell-associated barcodes were then called independently within each gene-expression sublibrary using the inflection point estimated by barcodeRanks in DropletUtils. A barcode was classified as cell associated when its total gene-expression UMI count was greater than or equal to the sublibrary-specific inflection-point threshold. For each called cell, we calculated the total gene-expression UMI count, number of detected genes and fractions of UMIs assigned to mitochondrial and ribosomal genes.

Quality control was subsequently performed separately within each treatment–collection-time group. Cells with unusually low or high total UMI counts or numbers of detected genes, or with elevated mitochondrial or ribosomal transcript fractions, were removed using condition-specific thresholds selected from the empirical distributions of these metrics. This condition-specific procedure accommodated differences in transcriptome complexity and mitochondrial and ribosomal RNA fractions across developmental stages and treatment conditions and avoided removal of particular cell states by a single global cutoff. The exact thresholds and the numbers of cells retained at each filtering step were recorded in the accompanying analysis notebook. After quality control, cells from all conditions and biological replicates were concatenated, while the raw UMI matrix and the mature, nascent and ambiguous count layers were preserved.

Downstream analyses were performed using Scanpy and associated Python packages in a Jupyter notebook. Quality-controlled counts were normalized for library size and transformed as log(1 + x). The first 50 principal components were computed using scanpy.pp.pca, and the nearest-neighbor distance matrix and neighborhood graph were constructed using scanpy.pp.neighbors with default parameters. Normalized expression values were used for exploratory visualization and marker-gene summaries, whereas raw UMI counts were retained for projection onto the fixed cNMF program space described below.

#### Projection of Hippo-modulated scRNA-seq dataset onto cNMF program space

To quantify the activity of the 50 cNMF programs inferred from the CC-Perturb-seq dataset in the independent Hippo-modulation scRNA-seq dataset, each quality-controlled cell was projected onto the fixed cNMF gene spectra. Raw UMI counts were matched to the reference spectra by gene symbol; duplicated gene symbols in the query matrix were summed. Counts were normalized to 10⁶ transcripts per cell using the total transcriptome UMI count. For each projection gene set, the corresponding columns of the query matrix and reference spectra were retained, and each program spectrum was normalized to sum to one across the selected genes.

Let X(*G*) denote the normalized cell-by-gene expression matrix for the genes in projection set G, and let H(*G*) denote the corresponding fixed 50-program cNMF spectra. Cell-level program usages were estimated by holding H(*G*) fixed and solving

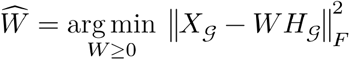

Projection was performed using the multiplicative-update implementation of non-negative matrix factorization in scikit-learn, with Frobenius loss, a maximum of 1,000 iterations, a convergence tolerance of 10⁻⁵ and a random seed of 0. The estimated usage coefficients were normalized within each cell to sum to one across the 50 programs.

#### Selection of genes for projection

We compared four gene sets: all 17,480 genes shared between the reference (CC-Perturb-Seq) and query (Hippo-modulation single cell) datasets; the union of the 300 highest-loading genes from each program, comprising 3,297 unique genes; and the union of the 200 most program-specific genes from each program, comprising 7,268 unique genes. For gene g and program k, program specificity was defined as

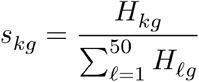

Within each program, genes were ranked first by program specificity and then by their absolute cNMF loading. The top 200 genes were retained for each program, and the union across all 50 programs was intersected with genes detected in the Hippo-modulation dataset.

Projection using all shared genes or the highest-loading genes produced diffuse assignment of some programs, particularly P35, across conditions in which the corresponding program-specific markers were weakly expressed. This behavior can arise when the independent query dataset contains expression variation that is not represented by the fixed 50-program reference basis. Because no new component can be learned during projection, residual variation outside the reference basis is approximated by combinations of the closest available programs, potentially inflating their estimated usage broadly across conditions. Restricting the fit to genes that distinguish among the reference programs reduced this leakage from broadly shared or poorly represented expression variation. The top-specific-200 projection was therefore used for all downstream analyses.

#### CLR transformation of projected program usages

Normalized program usages were converted to base-2 centered log-ratio (CLR) scores. For normalized usage u(i,k) of program k in cell i, values were bounded below at ε = 10⁻⁶ and renormalized:

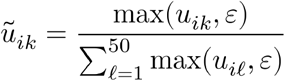

The final program score used for comparison was

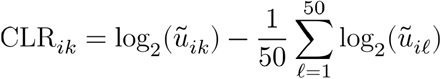

### 10. In vivo validation of Hippo pathway in zebrafish

#### Treatment with HIPPOi

Zebrafish embryos were exposed to either DMSO or hippo inhibitor (TDI-011536, 10uM) from shield stage (approximately 6 hours post fertilization (hpf)) to the 5-somite stage (11.5 hpf). At the end of the treatment, embryos were stored in Trizol for total RNA isolation, or fixed in 4% paraformaldehyde overnight at 4oC to study the etv2 expression using whole mount in situ hybridization (WISH)^116^. After WISH, the etv2 expression in posterior and anterior regions were scored categorically as low, medium or high.

#### Preparation for bulk RNA-seq

For bulk-RNAseq, Trizol saved samples (10 embryos per sample and n=5 per group) were lysed and processed for total RNA extraction using the Direct-zol kit (R2050; Zymogen) according to the user manual. 40 µg/µl concentration of total RNA was used for bulk-RNA sequencing and bioinformatic processing was performed by Plasmidsaurus (Plasmidsaurus, Arcadia, CA).

#### Bulk RNA-seq processing

Quality of the fastq files was assessed using FastQC v0.12.1. Reads were then quality filtered using fastp v0.24.0 with poly-X tail trimming, 3’ quality-based tail trimming, a minimum Phred quality score of 15, and a minimum length requirement of 50 bp. Quality-filtered reads were aligned to the reference genome using STAR aligner v2.7.11 with non-canonical splice junction removal and output of unmapped reads, followed by coordinate sorting using samtools v1.22.1. PCR and optical duplicates were removed using UMI-based deduplication with UMIcollapse v1.1.0. Alignment quality metrics, strand specificity, and read distribution across genomic features were assessed using RSeQC v5.0.4 and Qualimap v2.3, with results aggregated into a comprehensive quality control report using MultiQC v1.32. Gene-level expression quantification was performed using featureCounts (subread package v2.1.1) with strand-specific counting, multi-mapping read fractional assignment, exons and three prime UTR as the feature identifiers, and grouped by gene_id. Final gene counts were annotated with gene biotype and other metadata extracted from the reference GTF file. Sample-sample correlations for sample-sample heatmap and PCA were calculated on normalized counts (TMM, trimmed mean of M-values) using Pearson correlation. Differential expression was done with edgeR v4.0.16 using standard practice including filtering for low-expressed genes with edgeR::filterByExprwith default values. Functional enrichment, when available, is performed using gene set enrichment analysis with gseapy v0.12 using the MSigDB Hallmark gene set.

## Data Availability

Data is available on the IGVF Data Portal (https://data.igvf.org) and NCBI Gene Expression Omnibus (https://www.ncbi.nlm.nih.gov/geo/) at the following accessions:

Genome-wide CRISPRi Screen https://data.igvf.org/analysis-sets/IGVFDS9629VXQK/

Validation CRISPRi Screen https://data.igvf.org/analysis-sets/IGVFDS4708PCVX/ iPSC-arterial EC CC-Perturb-seq:

D0 https://data.igvf.org/analysis-sets/IGVFDS0463XTUH/

D1 https://data.igvf.org/analysis-sets/IGVFDS0991DBQZ/

D2 https://data.igvf.org/analysis-sets/IGVFDS4402ZGWJ/

D3 https://data.igvf.org/analysis-sets/IGVFDS2307REVJ/

scE2G predictions:

D0 https://data.igvf.org/prediction-sets/IGVFDS0171LUDN/

D1 https://data.igvf.org/prediction-sets/IGVFDS0128AJSL/

D2 https://data.igvf.org/prediction-sets/IGVFDS6804MAES/

D3 https://data.igvf.org/prediction-sets/IGVFDS9759CZLY/

ChromBPNet predictions, TF-MoDISco motifs, and Fi-NeMo motif hits:

D0 https://data.igvf.org/prediction-sets/IGVFDS5990HMDG/

D1 https://data.igvf.org/prediction-sets/IGVFDS7390PXAM/

D2 https://data.igvf.org/prediction-sets/IGVFDS9734VFMO/

D3 https://data.igvf.org/prediction-sets/IGVFDS5542PJNE/

Parse v3 CC-Perturb-seq, iPSC-K562 mix GSE352893

Small molecule Hippo scRNA-Seq GSE352939

Zebrafish RNA-Seq GSE350146

## Code Availability

cNMF workflow functions: https://github.com/EngreitzLab/cNMF_benchmarking/tree/main at commit 96d283ffb0e5d0d4e05565b70c33c095338049df

CRT: https://github.com/edtnguyen/programDE/tree/main at commit 11cfa3f5d9c8d400c15cd15e1e550fc11fe0f4f9

ABC^111^: https://github.com/broadinstitute/ABC-Enhancer-Gene-Prediction/main at commit 92ac50360231a6bcfd654f3147839d078be445d9 scE2G: https://github.com/EngreitzLab/scE2G at commit 9cadb5245ae1f26bb435f301a5fe305b04e1a3ae

ChromBPNet: https://github.com/EngreitzLab/ipsc-ec-chrombpnet at commit 4eca49c9b6ebc678968d3ce47e26dd23a84ac0ad

Code to reproduce selected figures: https://github.com/edtnguyen/cc-perturbseq-manuscript-analysis/ at commit 0ee11f3fc87e20852167a4873faf326adc10c024

## Acknowledgements

We thank members of the Engreitz and Loh labs for helpful discussions; Vivian Hecht for advice on applying ChromBPNet; Alex Rosenberg and Joey Pangallo for technical discussions; Adam Klie and Hannah Carter for sharing analysis tools; Stanford University and the Stanford Research Computing Center for providing computational resources and support as part of the Sherlock High-Performance Compute Cluster; and Stanford Genomics Service Center for DNA sequencing services. The Stanford Institute for Stem Cell Biology & Regenerative Medicine and the Stanford Stem Cell FACS Core Facility provided infrastructure support.

## Funding

This study was supported by NIH-NHGRI UM1HG011972 (J.M.E.), NIH-NHLBI R01HL164811 (J.M.E.), NIH-NHLBI P01HL180323 (J.M.E.), NIH-NHGRI U01HG013176 (J.M.E. and A.K.), the Applebaum Foundation (J.M.E.), Novo Nordisk Fonden NNF21SA0072102 (J.M.E.), the Breakthrough T1D Center of Excellence (K.M.L.), the Stanford Ludwig Center (K.M.L.), NIH-NHLBI R01HL153005 (S.S.), and the Anonymous, Fickel, and Gilbert families (K.M.L.). T.C.N. acknowledges support from a fellowship from the American Heart Association and The Children’s Heart Foundation (AHA 917090). H.D.K. and A.M. acknowledges support from the Stanford Bio-X PhD fellowship. H.D.K acknowledges support from NIH-NIGMS (5T32GM141828). D.A. acknowledges support from AHA/CHF Postdoctoral Fellowships (821920 and 23POSTCHF1019753) and the Stanford Maternal and Child Health Research Institute (MCHRI) Postdoctoral Fellowship. O.P. acknowledges support from the Swiss National Science Foundation (SNSF) through a Postdoc.Mobility fellowship (grant number P500-3_235165). T.Z. acknowledges support from NIH NIGMS (5T32GM007790) and a Predoctoral Fellowship from the American Heart Association (26PRE1565731). R.Z. acknowledges support from NIH NHGRI (T32 HG000044), an Impetus Grant, the Rosenkranz Foundation, and the Hevolution Foundation. M.U.S. acknowledges the support of an NSF Graduate Research Fellowship (DGE-1656518) and a graduate fellowship award from Knight-Hennessy Scholars at Stanford University. X.R.M. acknowledges support from NIH-NHGRI (2T32HG000044-21). K.M.L. was supported by the Packard Foundation Fellowship and The Anthony DiGenova Endowed Faculty Scholar Fund.

## Author Contributions

T.C.N., H.D.K., D.A., K.M.L. and J.M.E. designed the study and experiments.

T.C.N., T.Z., R.Z., S.D.C., N.T. and J.M.E. developed, optimized and tested the CC-Perturb-seq method.

D.A., C.J.M., C.M. and J.M.E. conducted and analyzed the genome-wide and validation CRISPRi screens to identify regulators.

D.A., C.J.M., O.P., E.M., H.Y.K. and X.R.M. generated and analyzed the 10x Multiome data.

L.T.A. and K.M.L. developed the differentiation protocol.

T.C.N. conducted the CC-Perturb-seq experiments during hiPSC-to-arterial EC differentiation.

T.C.N., O.P., Y.A.M., R.G. and J.M.E. processed and analyzed the CC-Perturb-seq data.

T.C.N. and T.Z. developed data-processing and analysis methods for CC-Perturb-seq.

Y.A.M., O.P., R.G. and H.Y.K. developed and applied gene-program analysis and annotation tools.

H.D.K. and K.M.L. designed and conducted experiments to characterize the Hippo pathway.

H.D.K., T.C.N., A.M., K.M.L. and J.M.E. conducted and analyzed single-cell RNA-seq experiments with Hippo activators and inhibitors.

S.P.R.B. and S.S. conducted the zebrafish experiments.

V.O. provided *ETV2^mCherry^* reporter hPSCs.

L.G., N.S. and A.K. supervised aspects of the study.

T.C.N., H.D.K., O.P., Y.A.M., K.M.L. and J.M.E. wrote the manuscript with input from all authors.

K.M.L. and J.M.E. supervised the study.

## Conflict of Interest Statement

T.C.N. and J.M.E. are listed as inventors on a patent application related to CC-Perturb-seq. J.M.E. has received speaking honoraria from GSK plc and Roche Genentech. K.M.L. is listed as an inventor on patent applications filed by Stanford University on hPSC differentiation into ECs.

## Supplementary Information

### Supplementary Note 1

**Circular capture improves guide RNA detection and can be integrated into Parse Evercode v3 barcoding**

We developed circular capture Perturb-seq (CC-Perturb-seq) to enable scalable CRISPR screens using split-pool combinatorial-indexing single-cell RNA sequencing. During the first round of reverse transcription, a guide RNA (gRNA)-specific primer labels Pol III-transcribed gRNAs with the same combinatorial cellular barcode used to label cellular mRNAs. Following amplification of the barcoded cDNA, guide-derived molecules are separated from gene-expression cDNA by SPRI size selection, strand-selected, circularized and enriched by inverse PCR. In contrast to CROP-seq-based capture, which detects a Pol II-transcribed reporter containing the guide sequence, circular capture directly detects the Pol III gRNA transcript. Guide detection can therefore be adapted to different gRNA scaffold architectures by changing the guide-specific reverse-transcription primer (**Fig. S2a**).

We first compared circular capture with CROP-seq capture in CRISPRi hiPSCs expressing gRNAs from the same CROP-seq vector. At matched guide-library sequencing depth, circular capture recovered a median of 192 guide UMIs per cell, compared with 30.5 guide UMIs per cell using CROP-seq capture, corresponding to a 6.3-fold increase (**Fig. S2b**). At least one guide was detected in 99.0% of cells processed by circular capture, compared with 75.3% of cells processed by CROP-seq capture (**Fig. S2c**). Median transcriptome recovery was similar between the two methods, with 14,200 transcriptome UMIs per cell for circular capture and 13,600 transcriptome UMIs per cell for CROP-seq capture **(Fig. S2d**). Read-downsampling analyses likewise showed broadly comparable transcript and detected-gene recovery between the two capture strategies (**Fig. S2e,k**), indicating that the circular-capture procedure did not compromise transcriptome complexity. Improved guide detection also increased the measured magnitude of on-target CRISPRi knockdown, consistent with reduced perturbation-assignment error (**Fig. S2f**).

Circular capture also increased guide recovery in telomerase-immortalized human aortic endothelial cells. The median UMI count for the most abundant guide in each cell was 110 with circular capture and 15 with CROP-seq capture, corresponding to a 7.3-fold increase (**Fig. S2g**). Thus, improved guide detection was observed in multiple cell types and was not restricted to hiPSCs.

We next tested whether circular capture could be integrated into the commercial Parse Evercode v3 split-pool barcoding workflow^117^, which uses a similar molecular biology protocol derived from SPLiT-seq.^33^ We conducted a cell-mixing experiment to evaluate gRNA capture efficiency and cleanliness. hiPSCs carrying a library of approximately 17,000 gRNAs at a MOI of approximately 5 and K562 cells carrying a library of approximately 97,000 gRNAs at a MOI of approximately 7 were fixed separately using the Parse fixation workflow. The fixed cell populations were then mixed at an approximately 1:1 ratio and loaded into the Parse Evercode v3 Round 1 plate.

Immediately before Round 1 reverse transcription, designated wells were supplemented with the CS1 guide-specific RT primer (**Table S1**), the DC1 guide-specific RT primer (**Table S1**), both primers or no guide-specific primer (see **Methods**). CS1 recognizes the CS1 direct-capture sequence^118^, whereas DC1 is compatible with a broader range of commonly used gRNA scaffold architectures. Following the modified Round 1 reaction, cells proceeded through the standard Parse Round 2 and Round 3 ligation-based barcoding steps. After Round 3, barcoded cDNA was captured, template-switched and amplified. A modified SPRI size-selection procedure separated the larger gene-expression cDNA from the shorter guide-derived cDNA. The guide-enriched fraction then underwent amplification, strand selection, single-stranded DNA circularization and guide-enrichment PCR according to the CC-Perturb-seq workflow.

Both guide-specific RT primers supported efficient guide capture in K562 cells and hiPSCs. For K562 cells, the median number of guide UMIs derived from the K562 guide pool was 217 with CS1, 238 with DC1 and 301 with the dual-primer condition. For hiPSCs, the corresponding median numbers of guide UMIs derived from the hiPSC guide pool were 138, 162 and 175, respectively **(Fig. S2i**).

Because K562 cells and hiPSCs carried distinct guide libraries but were combined only after fixation, the experiment also assessed the potential for mixing. In K562 cells, the median fractions of guide UMIs derived from the K562 guide pool were 0.86 with CS1, 0.96 with DC1 and 0.92 with both primers. In hiPSCs, the median fractions derived from the hiPSC guide pool were 0.72, 0.93 and 0.83, respectively (**Fig. S2j**).

Together, these experiments establish circular capture as an efficient and adaptable gRNA-detection strategy for split-pool Perturb-seq. Circular capture substantially increased guide recovery relative to reporter-based CROP-seq capture without reducing transcriptome recovery. The circular-capture module is compatible with many standard gRNA expression constructs and with multiple combinatorial indexing protocols. CC-Perturb-seq therefore increases the scalability and efficiency of conducting perturbation experiments with combinatorial indexing readouts.

### Supplementary Figures

**Fig. S1.**
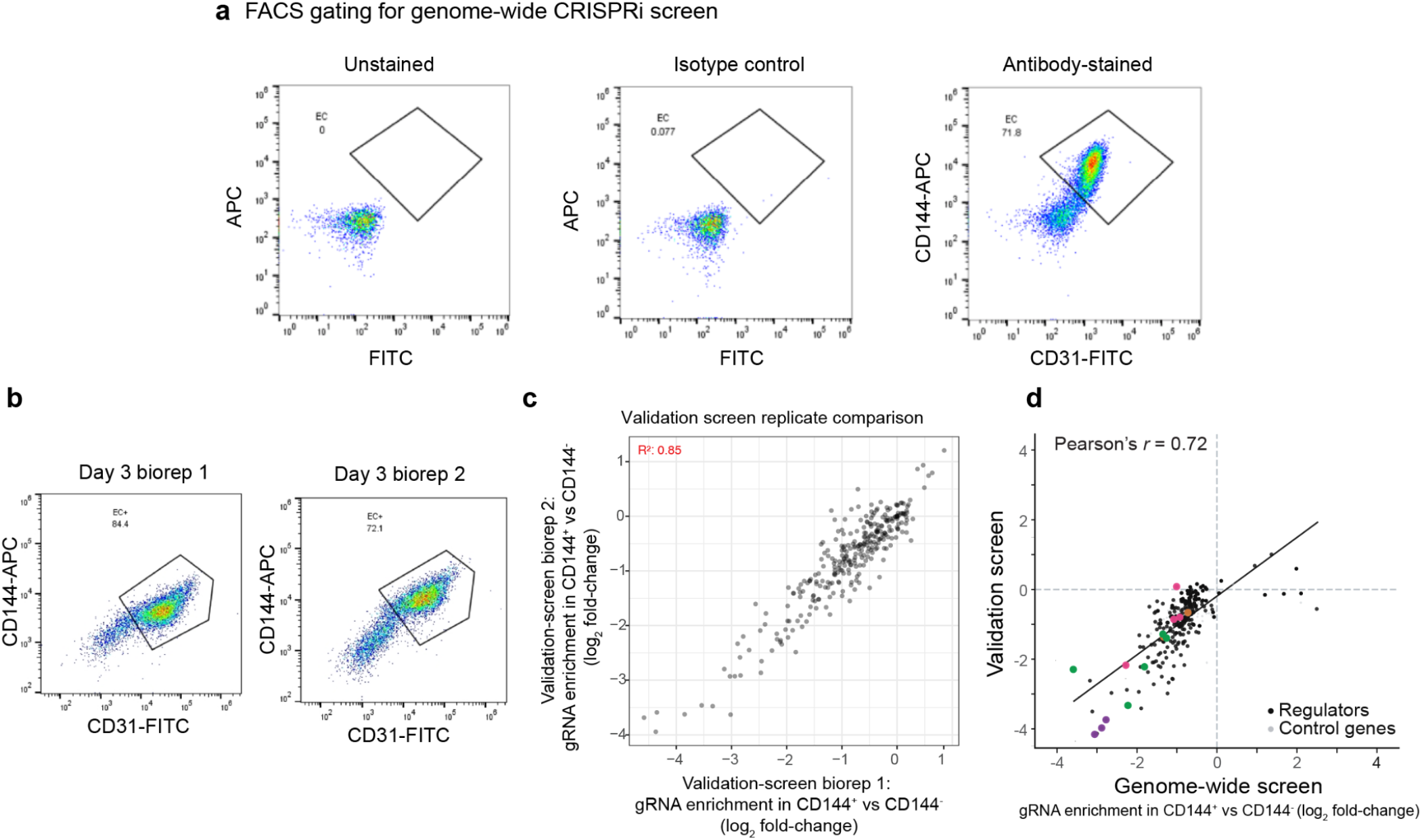
Genome-wide and focused CRISPRi screens for regulators of hiPSC–arterial EC differentiation. **a.** Representative flow-cytometry gates used in the genome-wide screen. Unstained and isotype-control samples define the negative region; antibody-stained cells are gated for CD144 and CD31 to quantify the endothelial population. **b.** CD144/CD31 flow-cytometry gates for the two biological replicates of the focused validation screen. **c.** Gene-level guide enrichment in CD144+ versus CD144− cells compared between validation-screen replicates. Each point represents one target gene; the identity line and correlation statistic summarize reproducibility. **d.** Comparison of gene-level effects in the genome-wide and focused validation screens. Axes show log₂ guide enrichment in CD144+ versus CD144− cells. Highlighted dots are colored by corresponding functional group in **Fig. 1**.

**Fig. S2.**
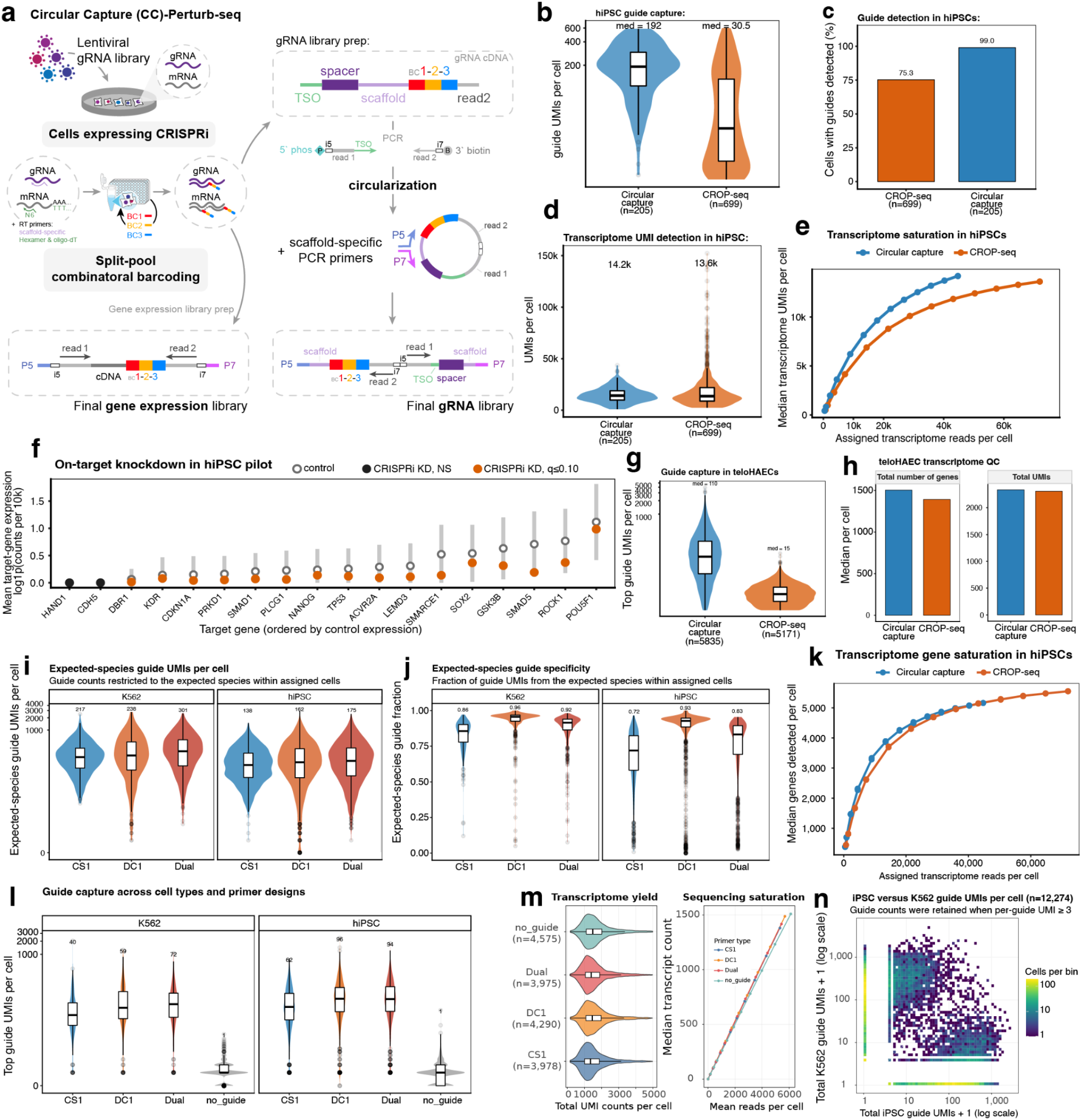
Development and benchmarking of CC-Perturb-seq. **a.** Schematic of CC-Perturb-seq. CRISPRi-expressing cells carrying a pooled lentiviral gRNA library are processed by split-pool combinatorial indexing. During Round 1 reverse transcription, cellular mRNAs are captured using poly(dT) and random-hexamer primers, whereas Pol III-transcribed gRNAs are captured using a scaffold-specific reverse-transcription primer. Following cDNA amplification, gene-expression and guide-derived molecules are separated by SPRI size selection. The guide-derived cDNA is strand-selected, circularized and amplified by inverse PCR to generate the final gRNA library. **b.** Head-to-head comparison of circular capture and CROP-seq capture in CRISPRi hiPSCs carrying gRNAs expressed from the same CROP-seq vector. Guide libraries were compared at matched sequencing depth. Violins show guide UMI counts per cell; embedded boxes indicate the median and interquartile range, and median values are shown above the distributions. **c.** Guide-detection rate in the hiPSC experiment described in **b**. Bars show the percentage of cells with at least one detected guide for CROP-seq capture and circular capture. **d.** Transcriptome recovery in the hiPSC experiment described in **b**. Violins show total gene-expression UMIs per cell for circular-capture and CROP-seq-capture libraries; embedded boxes indicate the median and interquartile range. **e.** Transcriptome sequencing saturation in the hiPSC experiment described in **b**. Curves show the median number of transcriptome UMIs recovered per cell as a function of assigned gene-expression reads per cell after read downsampling. **f.** On-target CRISPRi knockdown in the hiPSC experiment described in **b**. For each target gene, open points show mean normalized expression in control cells and filled points show mean expression in cells assigned a corresponding targeting guide. Target genes are ordered by their expression in control cells. Orange points indicate associations with *q* ≤ 0. 10; vertical bars show the corresponding confidence intervals. **g.** Independent comparison of circular capture and CROP-seq capture in TeloHAECs. Violins show the UMI count of the most abundant guide in each cell; embedded boxes indicate the median and interquartile range, and median values are shown above the distributions. **h.** Transcriptome recovery in the teloHAEC experiment described in **g**. **i.** Guide recovery in a hybrid Parse Evercode v3–CC-Perturb-seq compatibility experiment. hiPSCs carrying an approximately 17,000-gRNA library and K562 cells carrying an approximately 97,000-gRNA library were fixed separately, mixed after fixation and loaded into the Parse Round 1 plate. The Round 1 reverse-transcription reaction was supplemented with the CS1 primer, the DC1 primer or both primers; standard Parse Round 2 and Round 3 barcoding was followed by downstream CC-Perturb-seq guide enrichment. Violins show, separately for transcriptome-assigned K562 cells and hiPSCs, the total guide UMIs derived from the guide pool matching the assigned cell type. Embedded boxes indicate the median and interquartile range, and median values are shown above the distributions. **j.** Guide-pool specificity in the Parse compatibility experiment described in **i**. For each cell, specificity was calculated as the fraction of guide UMIs derived from the guide pool matching its transcriptome-assigned cell type. Violins show the resulting fractions for K562 cells and hiPSCs under the CS1, DC1 and dual-primer conditions. Embedded boxes indicate the median and interquartile range, and median values are shown above the distributions. **k.** Gene-detection saturation in the hiPSC experiment described in **b**. Curves show the median number of genes detected per cell as a function of assigned gene-expression reads per cell after read downsampling. **l.** Guide-capture efficiency across all four Round 1 primer conditions in the Parse compatibility experiment described in **i**. Violins show the UMI count of the most abundant guide in each transcriptome-assigned K562 cell or hiPSC under the CS1, DC1, dual-primer and no-guide-primer conditions. The no-guide-primer condition measures the background signal in the guide library. Embedded boxes indicate the median and interquartile range, and median values are shown above the distributions. **m.** Effect of adding guide-specific primers on transcriptome recovery in the Parse compatibility experiment described in **i**. Left, total transcriptome UMI counts per cell under the no-guide-primer, dual-primer, DC1 and CS1 conditions; labels indicate the number of cells in each condition. Right, median transcriptome UMI counts per cell as a function of mean assigned reads per cell after sequencing downsampling. Embedded boxes in the violin plots indicate the median and interquartile range. **n.** Guide-pool mixing in the Parse compatibility experiment described in **i**. Individual cell–guide observations supported by fewer than three UMIs were removed, after which UMIs were summed separately across the hiPSC and K562 guide pools within each cell. The two-dimensional histogram shows total hiPSC-pool versus K562-pool guide UMIs per cell on logarithmic axes. Color indicates the number of cells in each bin.

**Fig. S3.**
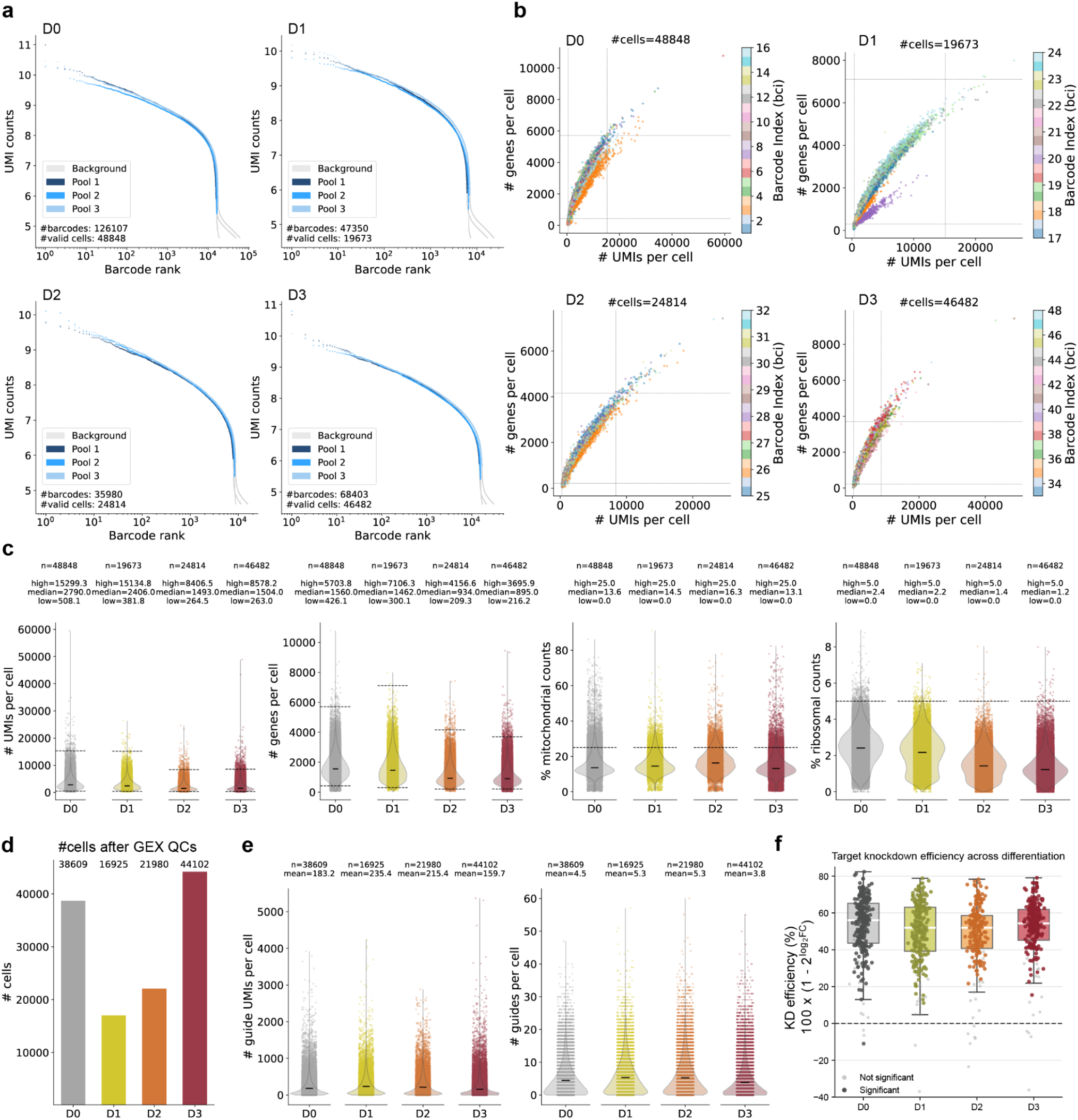
CC-Perturb-seq data processing and quality control. **a.** Barcode-rank plots and inflection-point cell calls for Days 0–3. Blue shades denote the three sublibraries and gray denotes background barcodes below the calling threshold. **b.** Total UMIs versus detected genes per cell, colored by first-round barcode index. Dashed lines show day-specific QC thresholds; low-quality barcode-index groups removed from Days 0 and 1 are indicated in the source analysis. **c.** Distributions of total UMIs, detected genes, mitochondrial fraction and ribosomal fraction before and after day-specific QC. Dashed lines show the thresholds defined by four normalized median absolute deviations for counts and by fixed 25% mitochondrial and 5% ribosomal cutoffs. **d.** Number of cells retained at each time point after gene-expression QC. **e.** Guide UMIs and provisional guide multiplicity per cell after gene-expression QC and before SCEPTRE assignment. **f.** Estimated target-gene knockdown by time point. Each point represents one target gene; boxes show the median and interquartile range. Dark points denote targets significant in the corresponding knockdown analysis and gray points denote non-significant targets

**Fig. S4.**
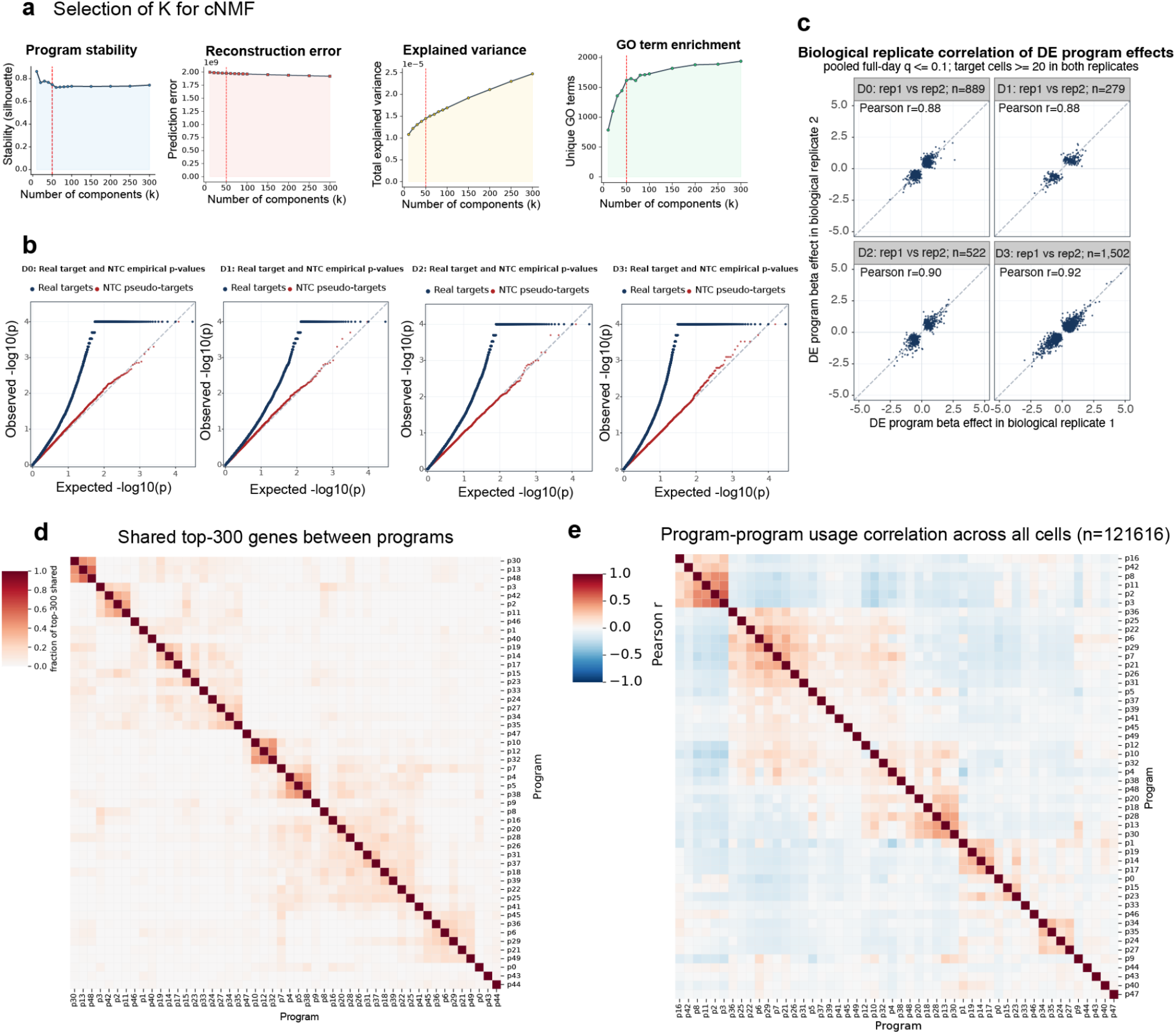
Selection, calibration and evaluation of the cNMF program solution. **a.** Program stability, reconstruction error, explained variance and cumulative GO-term enrichment across candidate values of K. K = 50 was selected as a balance between stable components, improved reconstruction and saturation of biological annotation. **b.** Day-specific Q–Q plots for real target–program tests and frequency-matched NTC pseudo-targets under the six-covariate conditional randomization test. The NTC curves assess null calibration; real targets show enrichment of small P values. **c.** Biological-replicate correlation of gene-level program effects at Days 0–3. Each point is a target–program pair passing the plotted inclusion criterion; Pearson correlations are shown in each panel. **d.** Pairwise overlap among the top 300 genes of each K = 50 program. Color indicates the number of shared genes; programs are shown in the same order on both axes. **e.** Pairwise Pearson correlation of program usage across all 121,616 CC-Perturb-seq cells. Red and blue indicate positive and negative correlation, respectively.

**Fig. S5.**
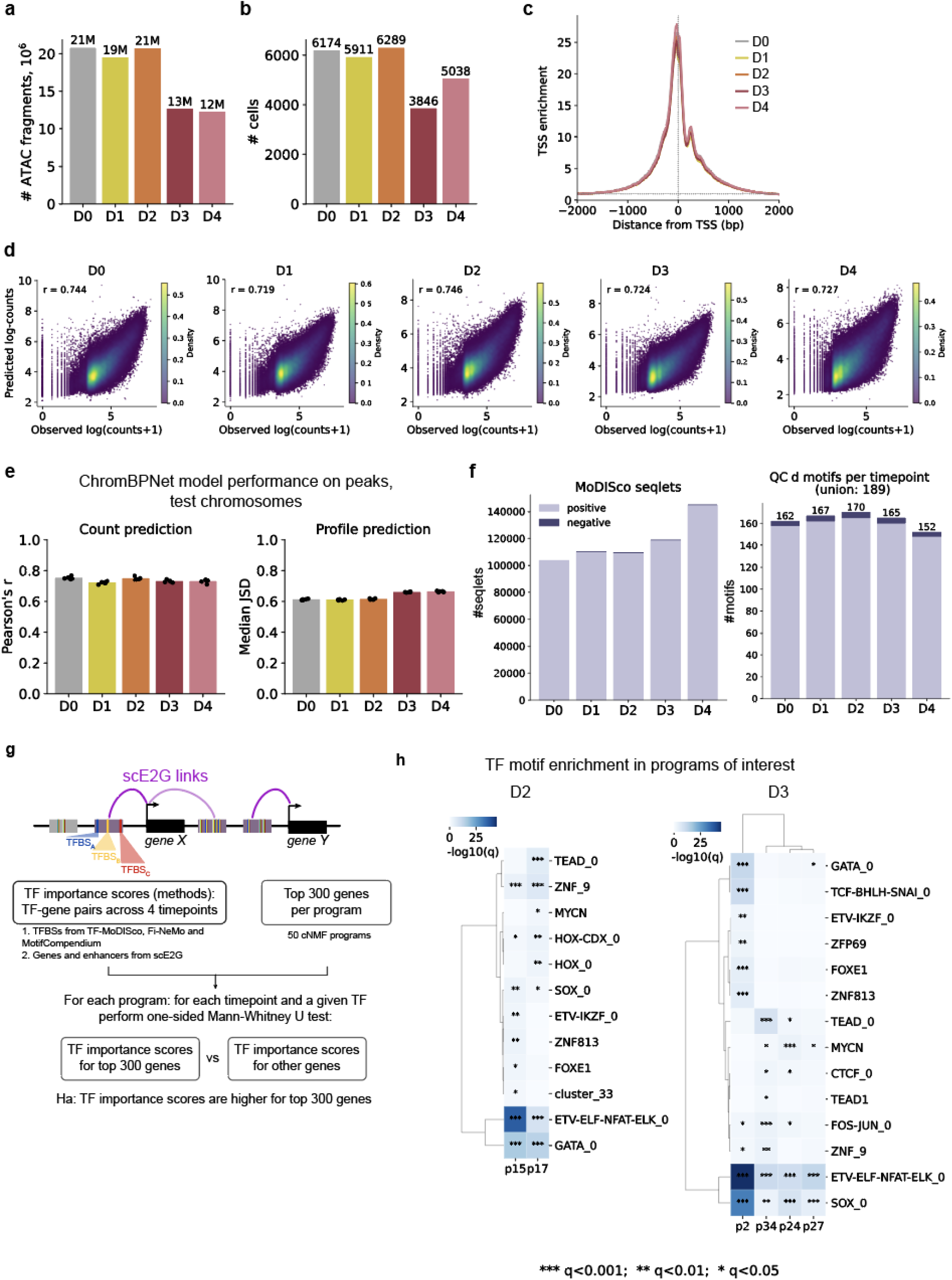
Multiome quality control, ChromBPNet performance and motif enrichment. **a.** Total pseudobulk ATAC fragments per differentiation time point. **b.** Number of cells per time point passing RNA-based quality control. **c.** Aggregate TSS-enrichment profiles for Days 0–4, normalized to the flanking insertion background. **d.** Observed versus ChromBPNet-predicted log counts in held-out peaks, pooled across five cross-validation folds for each time point. Pearson correlations are shown. **e.** ChromBPNet count-prediction correlation and profile-prediction Jensen–Shannon distance on held-out chromosomes. Points are folds and bars show the mean. **f.** Number of TF-MoDISco seqlets and Fi-NeMo motif instances passing the CWM-similarity threshold at each time point; darker segments denote negative contribution patterns. **g.** Analysis schematic linking scE2G enhancer–gene predictions and ChromBPNet/TF-MoDISco/Fi-NeMo motif scores to the top 300 genes of each cNMF program. **h.** Motif enrichment for selected Day 2 programs (P15 and P17) and Day 3 programs (P2, P24, P27 and P34). Rows are motif families or individual motif annotations; asterisks denote Benjamini–Hochberg-adjusted q-values (\**q* < 0.05, \*\**q* < 0.01, \*\*\**q* < 0.001).

**Fig. S6.**
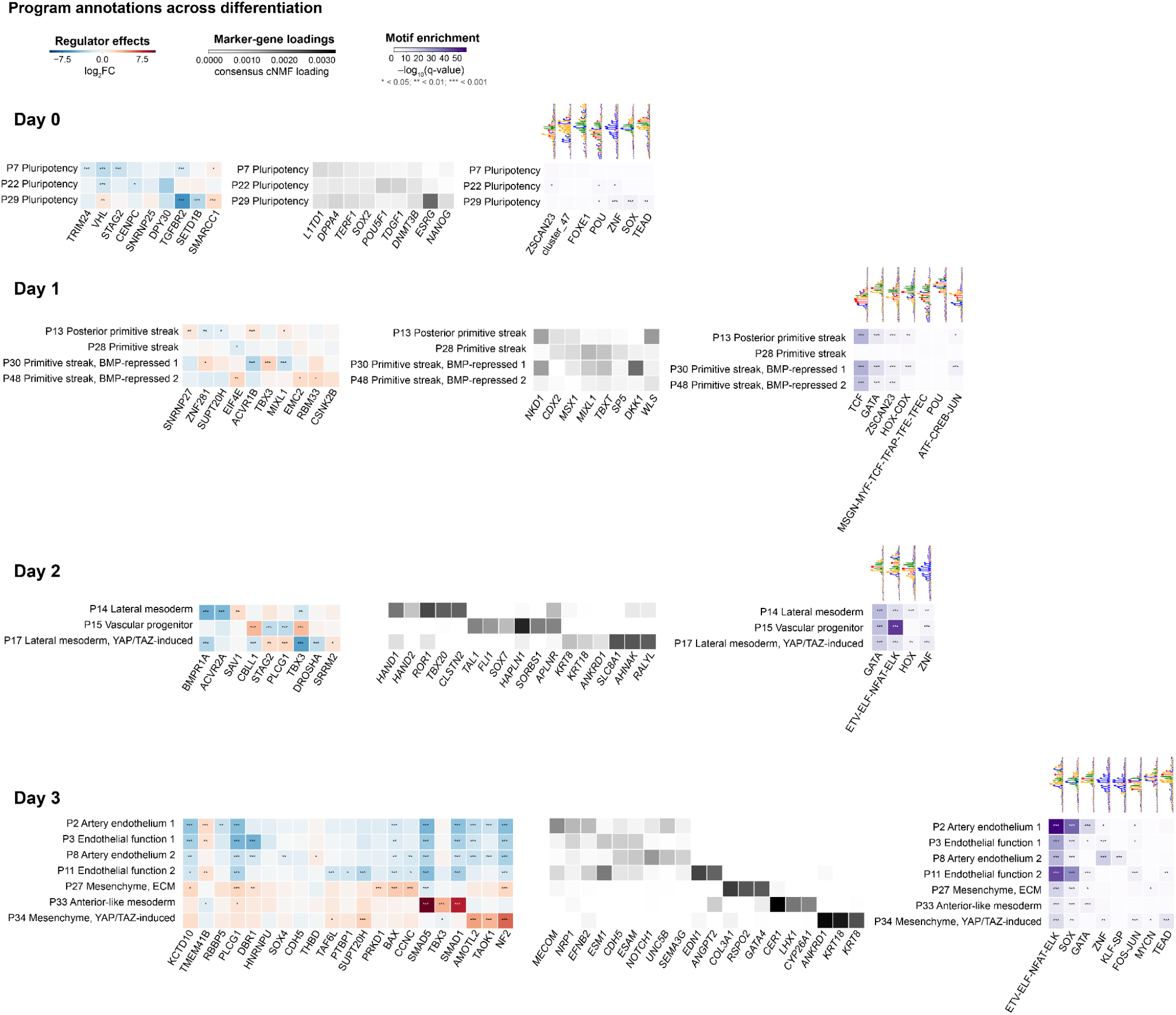
Integrated annotation of cNMF programs across hiPSC–arterial EC differentiation. Programs are grouped by the day at which they are most active (Days 0–3). For each day, rows are selected programs and the three aligned heatmaps show program-specific perturbation effects, representative gene loadings and transcription-factor motif enrichment in linked enhancers. In the perturbation heatmaps, red and blue indicate increased and decreased program activity after CRISPRi, respectively. In the loading heatmaps, darker gray indicates higher consensus cNMF loading. In the motif heatmaps, color indicates −log₁₀(q-value), with motif logos shown above the corresponding columns. Regulators, marker genes and motifs are unioned across programs within each day. Asterisks indicate the corresponding FDR or q-value threshold: * < 0.05, ** < 0.01 and *** < 0.001.

**Fig. S7.**
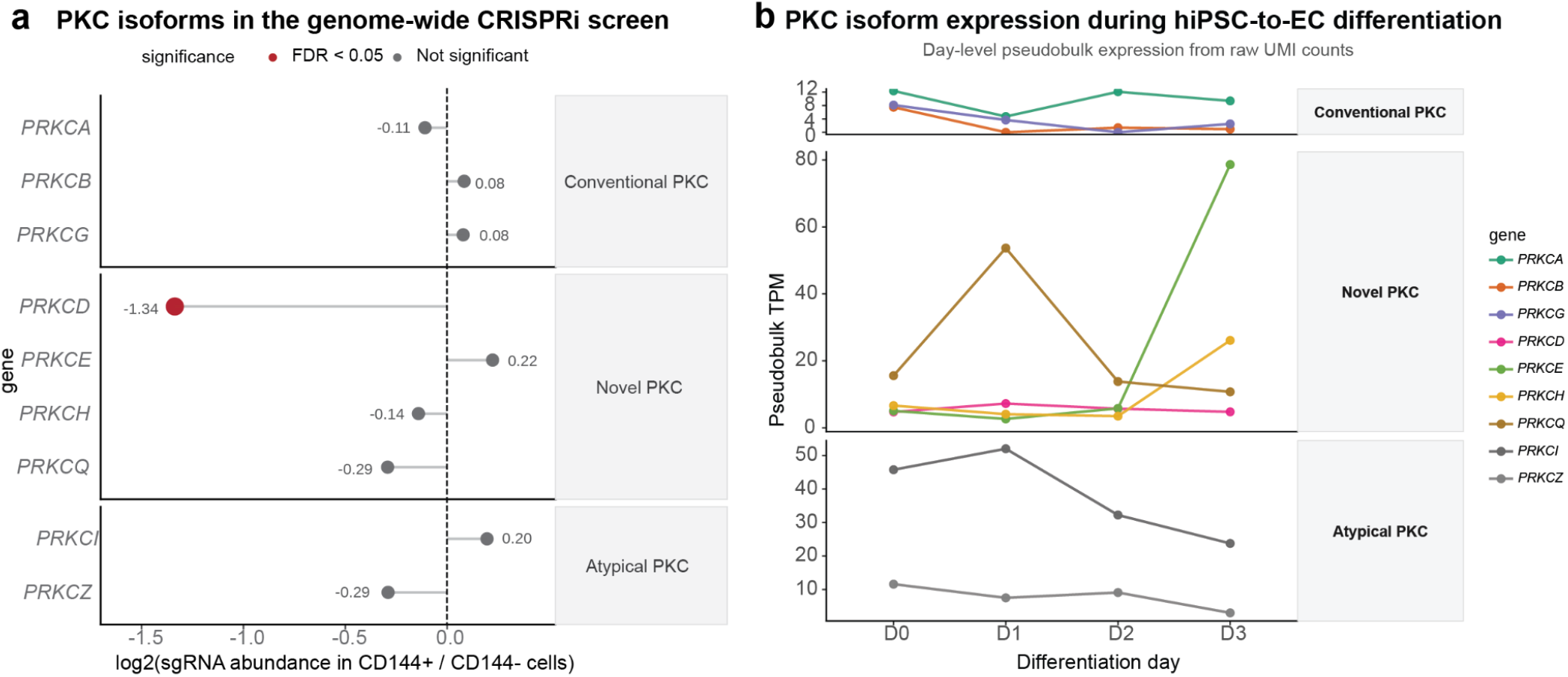
PKC isoform effects and expression during hiPSC-to-EC differentiation. **a.** Genome-wide CRISPRi screen effects for canonical PKC isoforms, shown as the log fold-change in sgRNA abundance in CD144-positive versus CD144-negative cells. Negative values indicate reduced EC differentiation after knockdown. Red points denote isoforms with directional MAGeCK FDR < 0.05; grey points are not significant. **b.** Day-level pseudobulk TPM of the same PKC isoforms across D0–D3 of differentiation, calculated from raw CC-Perturb-seq transcript counts using GENCODE v43 gene lengths.

**Fig. S8.**
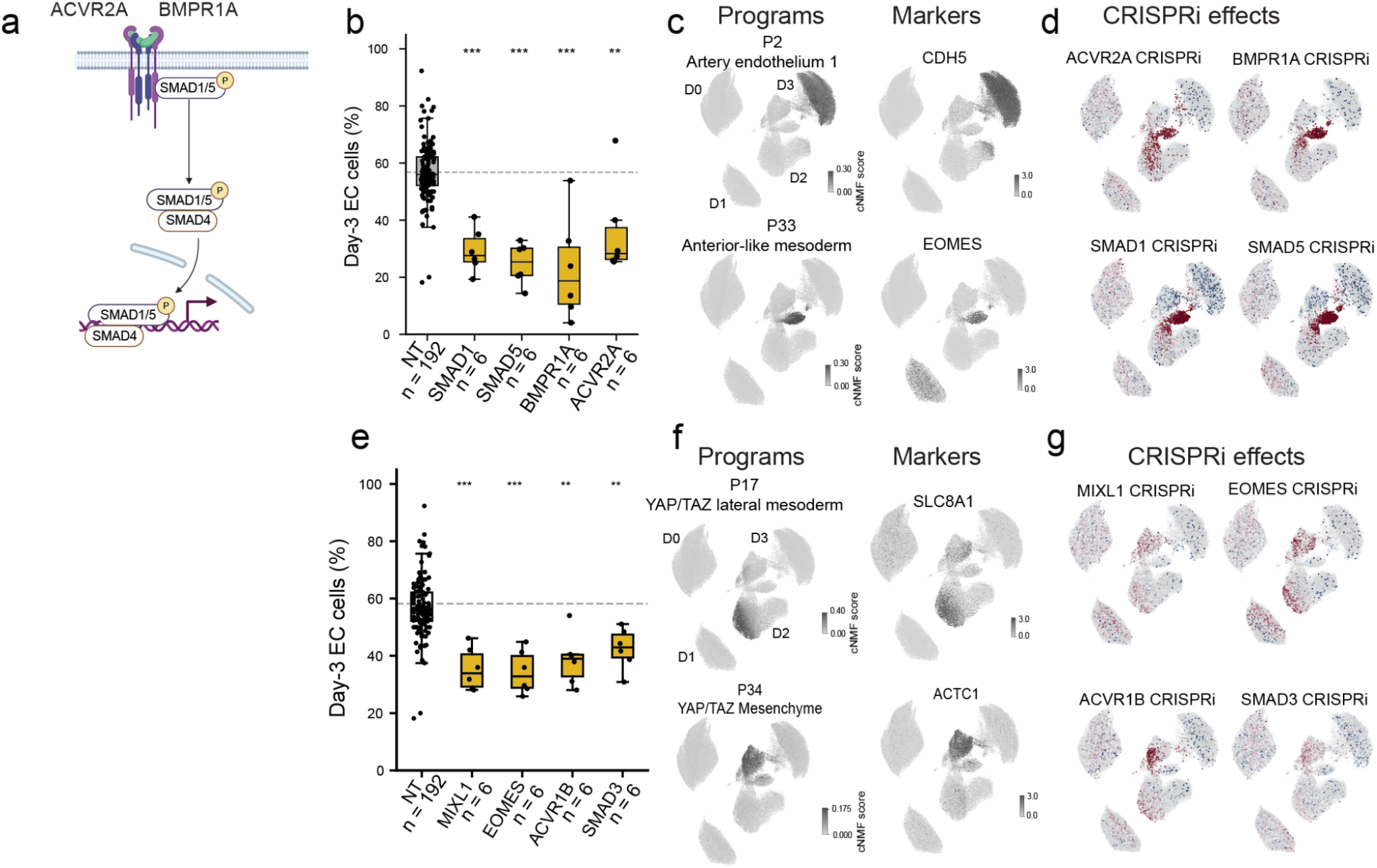
Distinct regulator groups alter EC differentiation through different program states. **a.** Schematic of BMP receptor signaling through SMAD1/5 and SMAD4. **b.** Day 3 EC fraction for non-targeting guides and guides targeting selected BMP-pathway components. Points represent guides; boxes show the median and interquartile range. **c.** UMAPs showing representative endothelial and primitive-streak/mesendoderm programs and marker genes in the CC-Perturb-seq data. **d.** Carrier-seeded neighborhood enrichment for ACVR2A, BMPR1A, SMAD1 and SMAD5 perturbations. **e.** Day 3 EC fraction for non-targeting guides and guides targeting EOMES, MIXL1, ACVR1B or SMAD3. **f.** UMAPs showing P17 and P34 together with representative marker genes SLC8A1 and ACTC1. **g.** Carrier-seeded neighborhood enrichment for MIXL1, EOMES, ACVR1B and SMAD3 perturbations.

**Fig S9.**
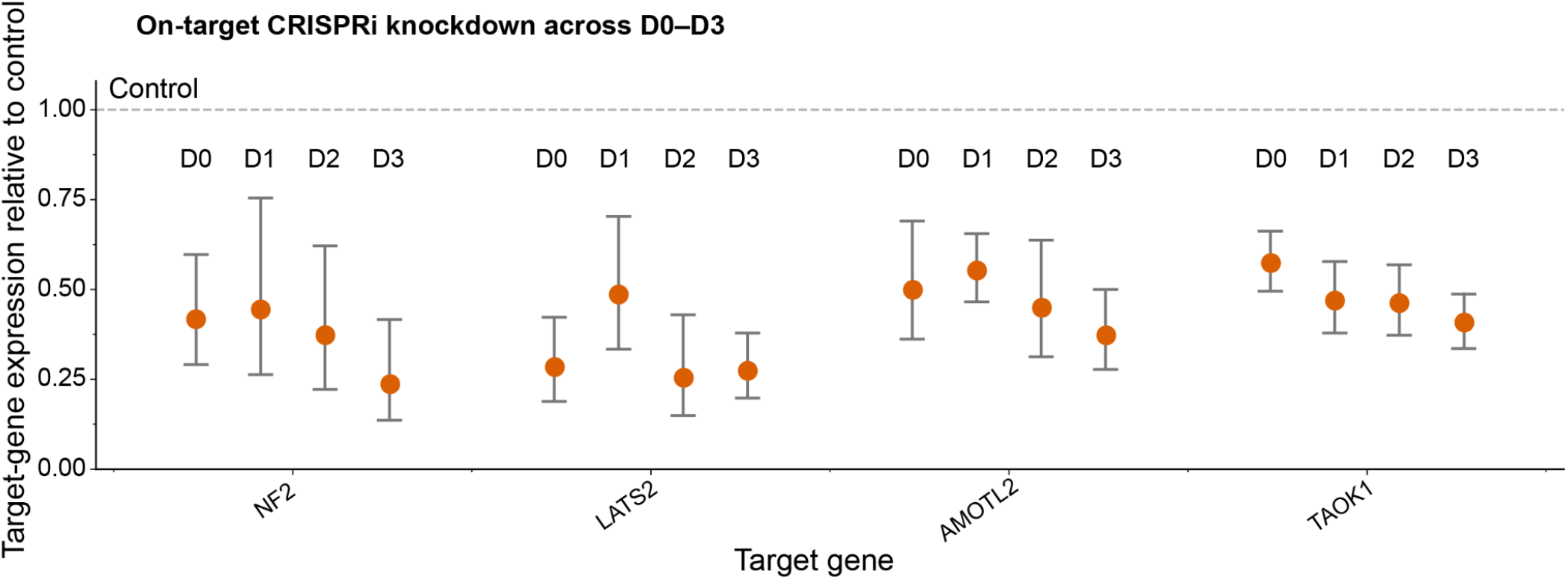
CRISPRi knockdown of Hippo pathway genes across differentiation days D0–D3. Target-gene expression is shown relative to the corresponding control population, with control expression normalized to 1. Error bars indicate approximate 95% model-based confidence intervals for the SCEPTRE fold-change estimate. SCEPTRE analyses were performed separately at each time point using high-MOI guide assignment and complement controls. All comparisons are significant with *q* < 0.05.

**Fig. S10.**
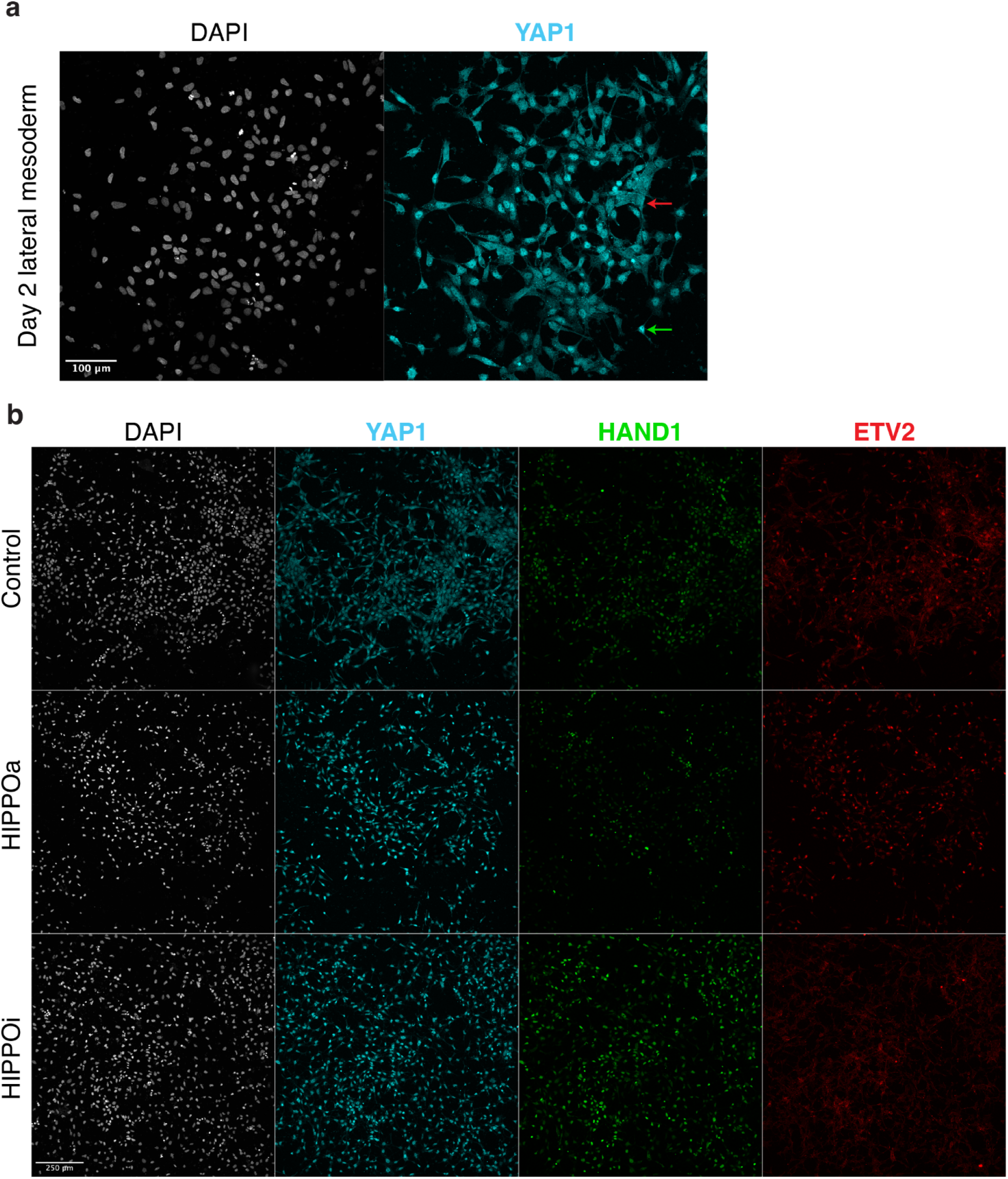
Immunostaining of YAP1, HAND1, and ETV2 in day 2 lateral mesoderm. **a.** Day 2 lateral mesoderm cells stained with DAPI (left) and anti-YAP1 antibody (right). Red arrow: cell with cytoplasmic YAP1. Green arrow: cell with nuclear YAP1. **b.** Day 2 lateral mesoderm cells treated with control lateral mesoderm media, lateral mesoderm supplemented with HIPPOa, or lateral mesoderm media supplemented with HIPPOi for 24 hours starting at day 1 of differentiation. Cells were fixed and costained for DAPI, YAP1, HAND1, and ETV2.

**Fig. S11.**
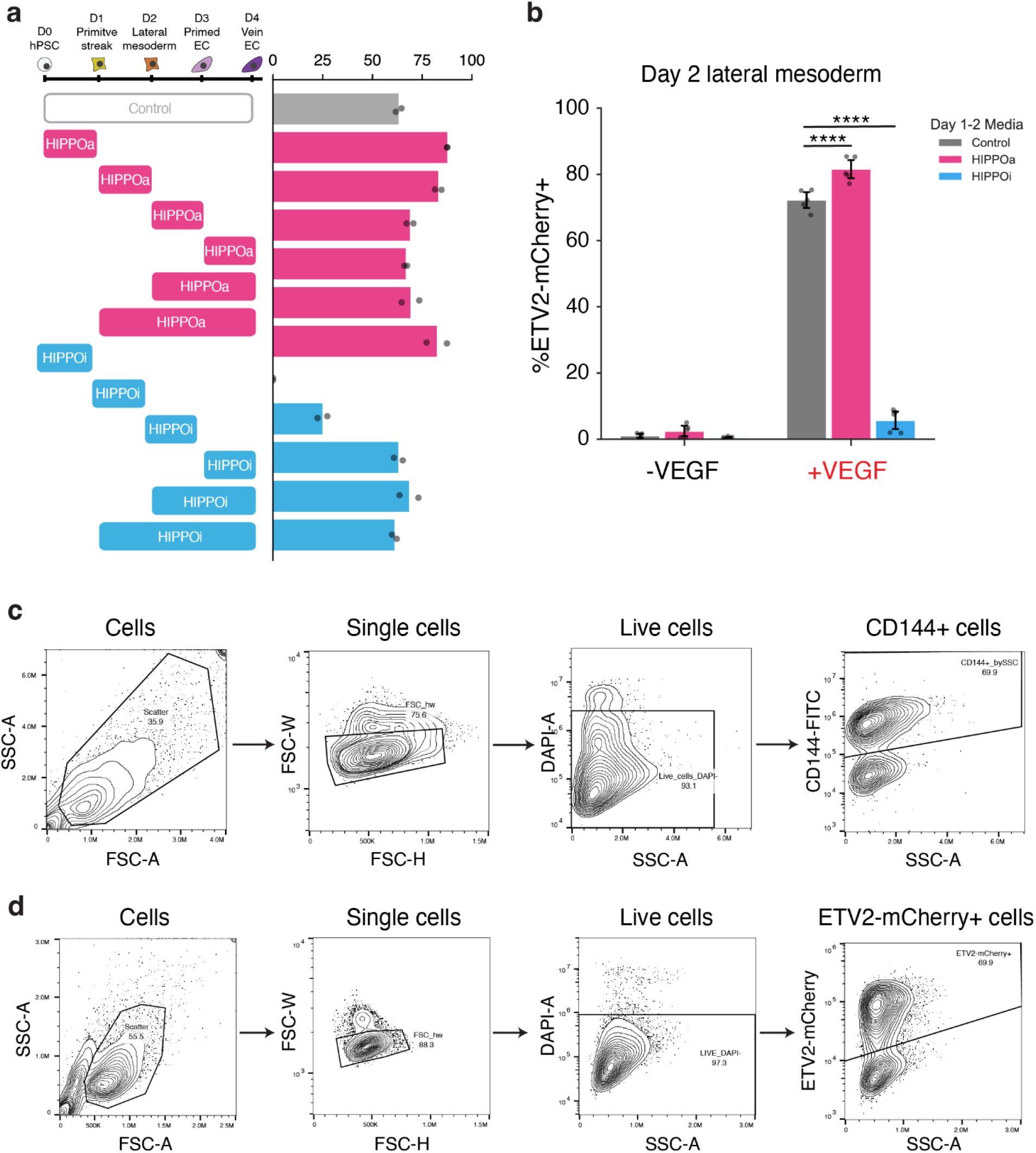
Effects of Hippo modulators on hPSC differentiation into ECs. **a.** Effects of HIPPOa or HIPPOi treatment at various stages of venous EC differentiation on EC purity assessed by flow cytometry of surface CD144. n=2 biological replicates from one experiment. **b.** Barplot showing purity of replicates from **Fig 4b**. **** = *P* < 10^-4^; Tukey’s Honestly Significant Difference test. n=5 biological replicates across two independent experiments. **c.** Representative gating scheme for artery differentiation experiments shown in **Fig. 3e**. **d.** Representative gating scheme for differentiation of ETV2-mCherry hPSCs into day 2 lateral mesoderm shown in **Fig. 4b**.

**Fig. S12.**
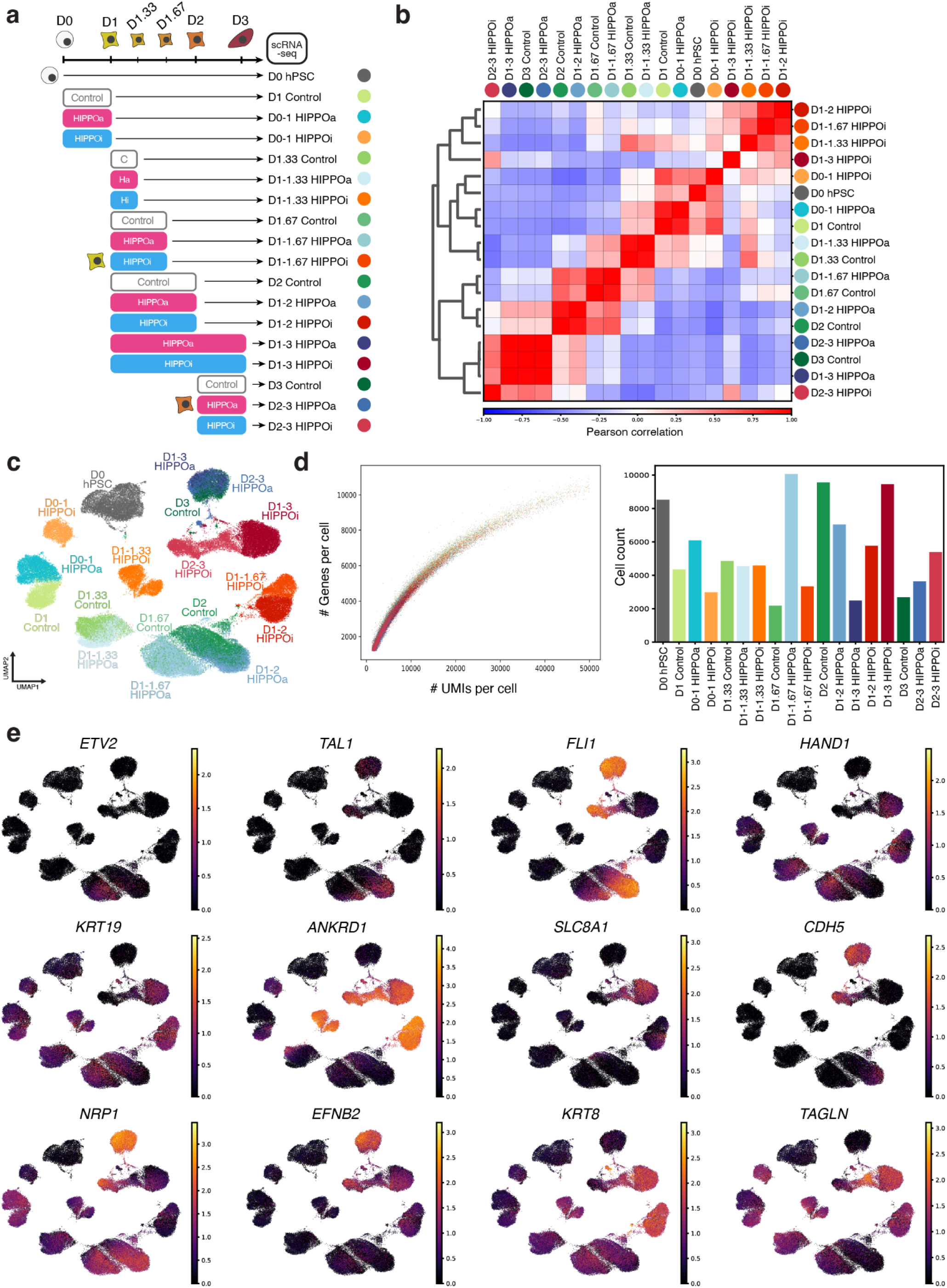
Extended scRNA-seq data from hPSCs differentiated in the presence of HIPPO pathway modulators. **a.** Schematic showing differentiation scheme and harvest timepoints. “C” = Control; “Ha” = HIPPOa; “Hi” = HIPPOi. **b.** Pearson correlation matrix between conditions in PCA space (50 PCs). **c.** UMAP plot of all conditions. **d.** Per-cell relationship between the number of detected genes and total gene-expression UMIs across experimental conditions (left), and the number of cells retained in each condition (right). **e.** UMAP plots showing expression of *ETV2, TAL1, FLI1, HAND1, KRT19, ANKRD1, SLC8A1, CDH5, NRP1, EFNB2, KRT8* and *TAGLN* across all conditions. Color bars show natural log normalized UMI counts.

**Fig. S13.**
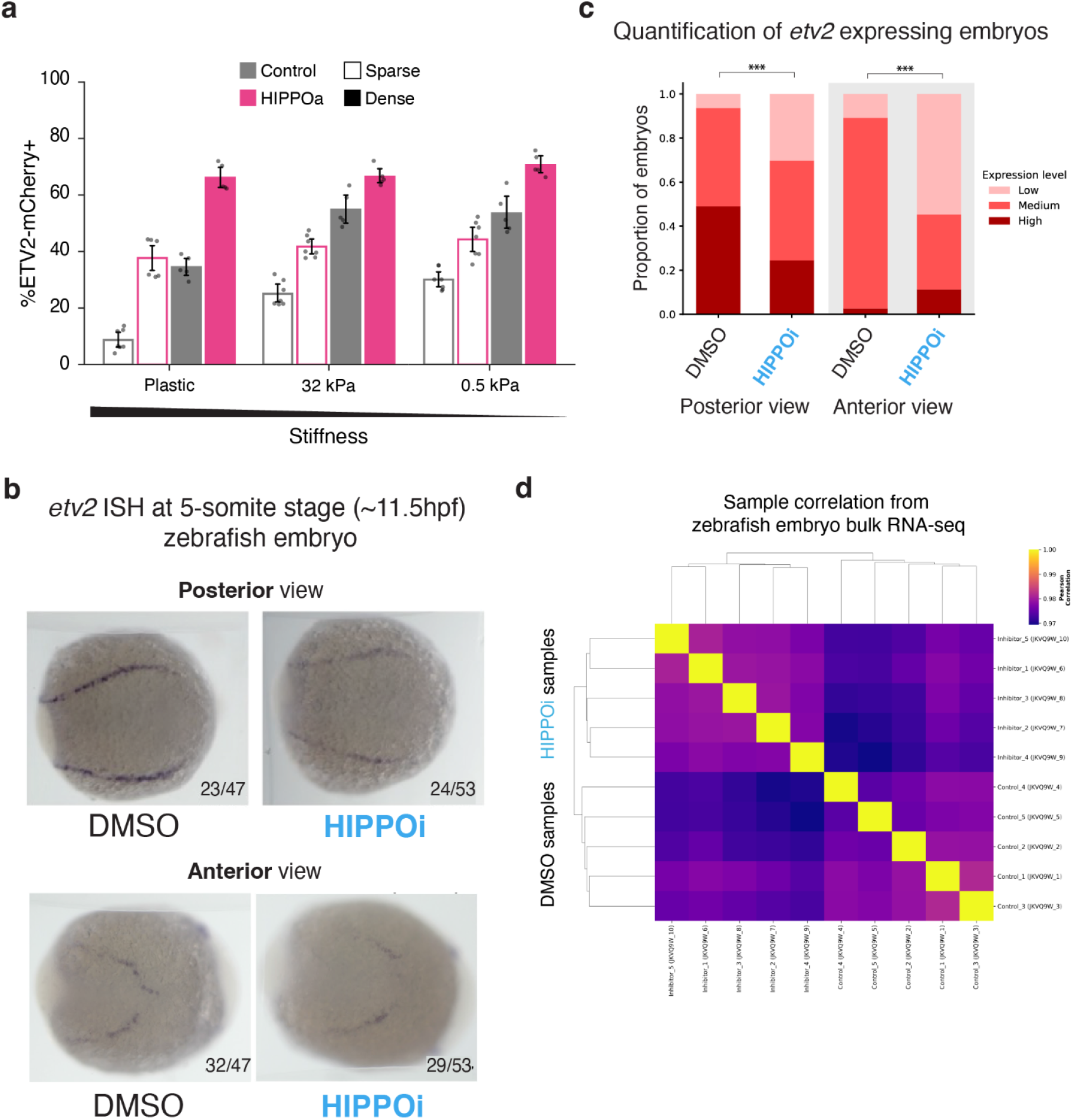
Extended data for mechanical and zebrafish experiments. **a.** Quantified ETV2-mCherry+ fraction by flow cytometry of cells replated on either plastic culture dishes or on gels of 32 kPa or 0.5 kPa stiffnesses at either sparse (2.6 × 10^4^ cells/cm^2^) or dense (7.8 × 10^4^ cells/cm^2^) cell density with or without HIPPOa treatment. Bars #1, 3, 5, 9 reflect the same data shown in **Fig. 5c**. Dots represent biological replicates harvested from three independent experiments. Replicates per experiment are as follows: n=1 for bars #1, 5, 7, 9, 11 from experiment #1; n=3 for control media conditions and n=2 for HIPPOa from experiment #2; n=3 for all conditions from experiment #3. **b.** *In situ* hybridization (ISH) of zebrafish embryos treated with DMSO or HIPPOi (10 μM TDI-011536) starting from the shield stage. Embryos analysed at 5-somite stage. Images representative of most frequent imaging phenotype (“low”, “medium”, or “high”). Left: posterior view; right: anterior view. **c.** Quantification of embryo ISH data shown in **b**. Embryos were classified as having “low,” “medium”, or “high” *etv2* expression. \*\*\**P* < 0.001, Fisher exact test. **d.** Heatmap of the Pearson correlation values between samples from the bulk RNA-seq experiment shown in **Fig. 5d**.

### Supplementary Tables

**Table S1.** CC-Perturb-seq sequences

**Table S2.** Summary of regulators of hiPSC-arterial_EC differentiation from CRISPRi differentiation screen

**Table S3.** Per-guide effects on EC differentiation efficiency from CC-Perturb-seq

**Table S4.** Program annotation table Annotation, temporal activity and perturbation-associated regulators of cNMF programs during hiPSC-to-endothelial differentiation

**Table S5.** Significant gene-perturbation effects on cNMF program activity across differentiation time points

**Table S6.** Zebrafish bulk RNA-seq

**Table S7.** qPCR primers and antibodies

